# Asymmetrical gene flow between coastal and inland dunes in a threatened digger wasp

**DOI:** 10.1101/2022.09.20.508247

**Authors:** Femke Batsleer, Matthieu Gallin, Moyra Delafonteyne, Daan Dekeukeleire, Filiep T’Jollyn, Pieter Vantieghem, An Vanden Broeck, Joachim Mergeay, Dirk Maes, Dries Bonte

## Abstract

Connectivity is a species- and landscape-specific measure that is key to species conservation in fragmented landscapes. However, information on connectivity is often lacking, especially for insects which are known to be severely declining. Patterns of gene flow constitute an indirect measure of functional landscape connectivity. We studied the population genetic structure of the rare digger wasp Bembix rostrata in coastal and inland regions in and near Belgium. The species is restricted to sandy pioneer vegetations for nesting and is well known for its philopatry as it does not easily colonize vacant habitat. It has markedly declined in the last century, especially in the inland region where open sand habitat has decreased in area and became highly fragmented. To assess within and between region connectivity, we used mating system independent population genetic methods suitable for haplodiploid species. We found more pronounced genetic structure in the small and isolated inland populations as compared to the well-connected coastal region. We also found a pattern of asymmetrical gene flow from coast to inland, including a few rare dispersal distances up to 200 to 300 km based on assignment tests. We point to demography, wind and difference in dispersal capacities as possible underlying factors that can explain the discrepancy in connectivity and asymmetrical gene flow between the different regions. Despite B. rostrata being a poor colonizer, gene flow between existing populations appeared not highly restricted, especially at the coast. Therefore, to improve the conservation status of B. rostrata, the primary focus should be to preserve and create sufficient habitat for this species to increase the number and quality of (meta)populations, rather than focusing on landscape connectivity itself.

## 2. Introduction

The decline in abundance and distribution of many insects has raised widespread public and political awareness on their biological value (Harvey et al. 2020; Didham et al. 2020; Wagner et al. 2021; Welti et al. 2021). As habitat loss and fragmentation have been identified as major drivers of this insect decline, a focus on connectivity conservation for (meta-)population persistence is essential and justified (Hanski et al. 1996; Hanski and Ovaskainen 2002; Haddad et al. 2015; Cardoso et al. 2020). Connectivity is a biological concept, in which fluxes of individuals between patches in heterogenous landscapes are determined by both landscape configuration and the species’ dispersal capacity. Patterns of gene flow reflect realized dispersal across multiple generations and can shed light on the functional landscape connectivity especially at large spatial scales (Kim and Sappington 2013; Hodgson et al. 2022; Maes and Van Dyck 2022). Responses to habitat loss and fragmentation are species- and context-dependent, because drivers of fragmentation can be diverse in identity, scale and intensity (Cheptou et al. 2017). Consequently, every landscape and its regional context is unique for each species and habitat connectivity remains difficult to generalize, even across different regions for the same species.

Dune habitats harbor a specific insect biodiversity with typical species of conservation interest (Maes and Bonte 2006; Provoost et al. 2011; De Ro et al. 2021). These sandy habitats in Belgium, both at the coast and inland, have gone through extensive—but different—landscape changes and fragmentation during the past decades or centuries. Coastal and inland dunes differ regarding their size, extent, history and nature of fragmentation, even though general levels of natural habitat fragmentation in Flanders (Belgium) are among the highest in Europe (European Environment Agency et al. 2011). Firstly, coastal dunes in Flanders are calcareous and form a narrow, linear system along the coast (Provoost et al. 2004; Decleer 2007). Coastal sandy habitats became fragmented at two scales. Primarily, urbanization from the interbellum period onwards decreased the total area of dunes significantly and resulted in a physical separation of the larger dune entities. In parallel, loss of low-intensity agricultural practices (including grazing of livestock) and obstruction of sand dynamics due to urbanization stimulated the succession and shrub development, at the cost of the open, early-succession or pioneer dune habitats (Provoost et al. 2011). Large herbivores have been introduced in many coastal dune reserves during the last three decades to revitalize dune dynamics (Provoost et al. 2004), but might have mixed effects on local arthropod species due to intense trampling (Bonte and Maes 2008; van Klink et al. 2015; Batsleer et al. 2022b). Second and contrastingly, inland sandy soils in Flanders are acidic (Decleer 2007). In this region, large open heathland and land dune systems were heavily afforested since the 19th century (De Keersmaeker et al. 2015). Later, in the second half of the 20^th^ century—parallel to, but more severe than at the coast—the remaining open habitat patches became further build-up and hence, smaller and more fragmented. Secondary loss and fragmentation of the remaining sandy habitat patches took place, here due to acidification and eutrophication leading to a continuous grass encroachment of sparsely vegetated sand areas on lime-poor soils (Schneiders et al. 2020).

The digger wasp Bembix rostrata is a univoltine habitat-specialist associated with dynamic sandy habitats with early-successional vegetation: grey dunes in coastal areas (EU Habitats Directive habitat 2130) and dry sandy heaths and inland dunes with open grasslands in inland areas (habitats 2310 and 2330 respectively). The species occurs in Europe and Central Asia, with a northern limit reaching south Scandinavia (Bitsch et al. 1997). In several European countries, B. rostrata has declined during the 20^th^ century and is considered a Red List species in several regions in Germany and is protected in Wallonia, Belgium (Blösch 2000; Jacobs 2000; Klein and Lefeber 2004; Barbier 2007; Bogusch et al. 2021). In Belgium and the Netherlands, mostly inland populations were lost, resulting in a distribution with local strongholds at the coast and more fragmented or isolated populations present inland (Klein and Lefeber 2004). Bembix rostrata is labeled as a philopatric species that does not easily colonize vacant habitat and aggregates stay present at the same location for many consecutive years (Nielsen 1945; Larsson 1986; Bogusch et al. 2021). This presumed philopatry is likely linked to the species gregarious life-style where females base their nest choice on the presence of conspecifics (Batsleer et al. 2022a). Given the typical dynamic character of the species’ habitat (pioneer dune or other sandy vegetations), philopatry should be highly disadvantageous and eventually put the species’ at risk if new early-successional sites cannot be colonized (Bogusch et al. 2021). Hence, as for other species from early-successional habitat, good dispersal capacities would be expected despite the overall sedentary life style during breeding (e.g. Andrena vaga: Černá et al. 2013; Exeler et al. 2008). Correct information on the species’ dispersal capacity is, therefore, essential to guide future conservation strategies.

Given the species’ conservation flagship status in Belgium as emblematic ground-nesting hymenopteran (Batsleer et al. 2021), we studied connectivity through gene flow between B. rostrata populations within and between coastal and inland fragmented sandy habitats in Belgium and bordering areas. Our main question is whether and how different populations and regions are genetically connected to each other. We amplified 21 microsatellite markers and used a non-lethal sampling method with wing clips (Châline et al. 2004) to minimize the impact of sampling on the often small and/or geographically isolated populations. As the species has a haplodiploid sex-determination system, we used population genetic analyses that are mating system independent and do not incorporate a diploid population genetic model.

## 3. Material and methods

### 3.1. Study species

Bembix rostrata is a univoltine specialized, gregariously nesting digger wasp from sandy habitats with sparse vegetation (Larsson 1986; Klein and Lefeber 2004). Adults are active in summer, showing protandry: females are directly mated when emerging by the guarding males, who emerge one to five days earlier (Wiklund and Fagerström 1977; Schöne and Tengö 1981; Evans and O’Neill 2007). Females show brood care: one individual constructs one nest burrow at a time in which it progressively provisions a single larva with flies (Nielsen 1945; Field et al. 2020). An estimated of up to 5 nests are produced, each with one offspring (Larsson and Tengö 1989). There are no overlapping generations, as the species overwinters as prepupa.

### 3.2. Study sites and sampling

Sampling took place during the summers of 2018, 2020 and 2021. Samples were taken across 49 Belgian populations, three French populations and one Dutch population (Fig. 1). Detailed information about each sampling site—region, name, coordinates, sample year(s), number of samples—can be found in supplementary material S1. A large part of the Belgian coastal populations and two inland populations were sampled both in 2018 and 2020 and are used to check if samples from different years can be pooled in the subsequent analyses (see below, hierarchical Analysis of Molecular Variance; AMOVA). Only females were sampled, to solely use diploid individuals in the population genetic analyses for this haplodiploid mating system.

**Figure 1:**
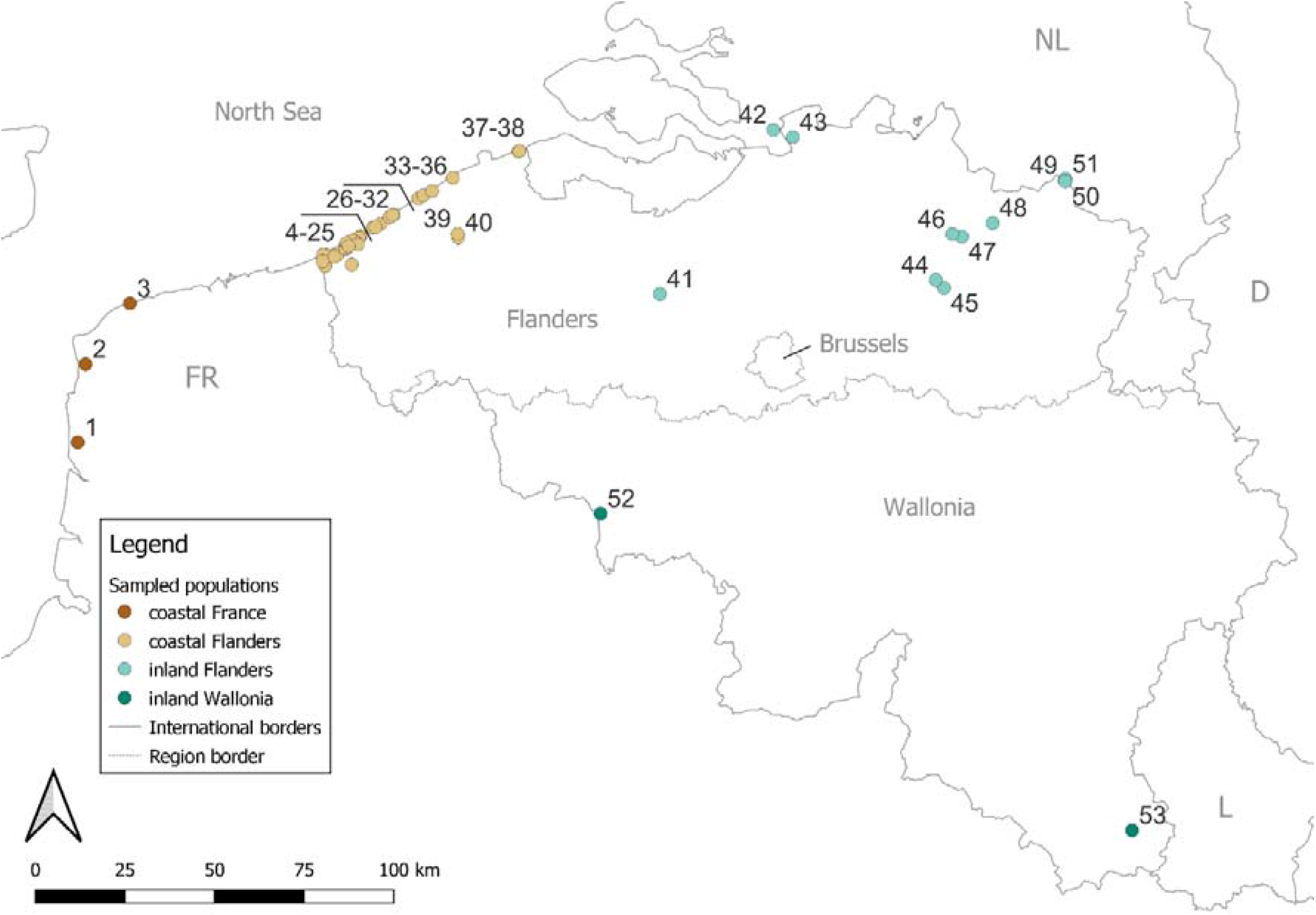
Overview map with locations of sampling. Populations 1-3 in coastal France (dark brown dots), 4-40 in coastal Flanders (light brown dots), 41-51 in inland Flanders (light green dots), 52-53 in inland Wallonia (dark green dots). Neighboring countries and the three administrative regions of Belgium are indicated: FR (France), NL (Netherlands), D (Germany), L (Luxembourg); Flanders (Flemish region), Wallonia (Walloon region) and Brussels (Brussels-capital region).

The populations were a priori divided into four regions based on geographical configuration and sampling design. In Flanders (north part of Belgium) all populations at the coast or inland known to exist at the time of sampling were covered. In Wallonia (southern part of Belgium) the two main known large populations were covered. At the French coast bordering coastal Flanders, three extra populations were sampled opportunistically, to be able to consider genetic links further along the French coast, although intermediate populations certainly exist. We considered coastal France as an a priori separate region (Fig. 1) to avoid biased interpretation of gene flow patterns due to this incomplete sampling coverage.

As it is estimated that B. rostrata has 5 nests (or equivalent number of offspring) per female (Larsson and Tengö 1989), population sizes grow slowly, especially in isolated fragments. Therefore, to minimize the impact of sampling on the populations, we used a non-lethal sampling method with wing clips, a method shown to give good-quality DNA for microsatellite PCR amplification (Châline et al. 2004). Tips of both forewings from live digger wasp individuals were cut. Both wingtips were stored in absolute ethanol, stored in a refrigerator at 4°C after sampling and transferred to a freezer (−18°C) for long-term storage. For each sample, individual coordinates of the capture position (most often a nest) were noted.

### 3.3. DNA extraction and PCR amplification

Genomic DNA was extracted from the wing tips with a protocol based on Chelex (Biorad; details in supplementary material S2). The development of species-specific microsatellites was outsourced to AllGenetics® (A Coruña, Spain), who provided 500 non-tested microsatellite primers and tested 72 of those biologically with 11 individuals. Of those, we selected 33 polymorphic microsatellites (based on polymorphism, size range, and length of repeat motif) and rearranged them with the program multiplex manager (Holleley and Geerts 2009) in 3 pairs of primer-multiplexes for PCR and amplified these for each sample. Details on the PCR-conditions and multiplexes are in supplementary material S2; characteristics of the primers are in supplementary material S3. PCR products were run on an ABI 3500 analyzer with the GeneScan-600 LIZ size standard and the electropherograms were scored using Geneious Prime (Biomatters). Samples from three recaptured individuals were blindly and randomly added to the workflow. They popped up as duplicate genotypes in the first part of the data analysis, so we were confident the scoring error was minimal.

### 3.4. Genetic data analysis

#### Hardy-Weinberg, linkage disequilibrium and null alleles

To exclude microsatellites that are uninformative or have artefacts, the assumptions of Hardy-Weinberg (HW) equilibrium and null alleles at individual loci (non-amplified alleles; Chapuis and Estoup 2007), and of no linkage disequilibrium (LD) across pairs of loci were examined before subsequent genetic analysis (Waples 2015). HW deviations, null alleles and LD deviations were calculated and examined with the R-packages pegas (Paradis 2010), PopGenReport (Adamack and Gruber 2014) and poppr (Kamvar et al. 2014) respectively, for populations that had at least 10 samples. Assumptions testing followed the general reasoning and multiple testing from Waples (2015), see supplementary material S5.

#### Hierarchical AMOVA to validate pooled samples from different years

To test if samples from populations of different years (2018, 2020 and 2021) could be pooled, a hierarchical Analysis of Molecular Variance (AMOVA) was carried out using the package poppr (Kamvar et al. 2014). To test the null hypothesis of no population structure between years, 23 populations that were sampled twice (2018 and 2020; table S1.1) were used in this test and year was hierarchically nested within population.

#### Population-level statistics

For each population, we calculated the number of private alleles (NP) with the R-package poppr (Kamvar et al. 2014). Rarefied allelic richness (AR), expected heterozygosity (H_e_), observed heterozygosity (H_o_) and inbreeding coefficient (F_IS_) were calculated with the r-package hierfstat (Goudet and Jombart 2020). These measures should only be interpreted relatively within the studied haplodiploid system.

To check the robustness of the population-level statistics in light of skewed sample sizes per population (supplementary material S1), we repeated these population-level statistics on a subsampled dataset. For this dataset, we randomly selected 10 samples if a population has more than 10 samples and omitted the two populations of sample size of 5.

#### Genetic distance

Genetic distance between populations was quantified using Nei’s standard genetic distance D_S_ (Nei 1972) calculated with the package hierfstat (Takezaki and Nei 1996; Goudet and Jombart 2020). To check which populations have on average the highest genetic distance to other populations, the mean D_S_ per population was calculated. Similar calculations for genetic differentiation indices F_ST_ and Jost’s D (Weir and Cockerham 1984; Jost 2008; Keenan et al. 2013) can be found in supplementary material S4, but are considered less suitable for haplodiploids as they are by definition based on expected diploid allele frequencies under Hardy-Weinberg equilibrium.

To check robustness of this analysis against skewed sample sizes (see previous section), we repeated the analysis on a subsampled dataset (10 samples are randomly selected if a population has more than 10 samples and the two populations of sample size of 5 were omitted).

#### Isolation-by-distance

We compared patterns of isolation-by-distance (IBD) between coastal and inland populations. Only samples from Flanders were used, where sampling was spatially covering all populations known to exist at that time. This way, a balanced distribution of geographic distances across the range of possible distances. IBD was examined with the multivariate approach distance-based redundancy analysis (dbRDA) (Diniz-Filho et al. 2013) because the suitability of Mantel tests to examine IBD is highly debated (Meirmans 2015). First, a Principal Coordinate Analysis (PCoA) to the Nei’s genetic distance D_S_ matrix was applied. The resulting PCoA-axes were then used as a response matrix in a Redundancy Analysis (RDA) to correlate them with the geographic coordinates. The adjusted coefficient of determination R² of the RDA’s (the proportion explained by the constrained axes) was used to compare the strength of IBD for coastal and inland populations. Permutations tests (n = 999) were used to check if the R² significantly differed from zero. Nei’s genetic distance D_S_ between populations was plotted against the pairwise geographical distance for the two regions. To check if the intercept and/or slope differ between the two regions, a permutation test was applied with the R package lmPerm with a maximum of 10,000 iterations (Wheeler and Torchiano 2016) with the formula D_S_ ∼ distance + region + distance:region. In general, realized gene flow is dependent on population sizes and dispersal capacity (∼N·m), with spatial configuration of populations a confounding factor. A different intercept in IBD (for the same species) for different regions would be mainly related to differences in population sizes (as N decreases, all else being equal, differentiation will increase due to total number of migrants) and a differing slope to different dispersal capacities.

#### Discriminant Analysis of Principal Components (DAPC)

A Discriminant Analysis of Principal Components (DAPC) was performed with the R-package adegenet to explore between-population structure and differentiation (Jombart 2008; Jombart et al. 2010). DAPC is a multivariate statistical approach wherein data on individual allele frequencies is first transformed using a principal component analysis (PCA) and subsequently a discriminant analysis (DA) is performed. Genetic variation is partitioned into a between-group and a within-group component, maximizing discrimination between groups (i.e. populations in this case). DAPC does not assume a population genetic model, which make it more suitable for haplodiploid mating systems than Bayesian clustering algorithms to analyze between-population structure (Jombart et al. 2010; Grünwald and Goss 2011). We performed a DAPC for all populations together and the coastal and inland populations separately. Populations are used as the a priori groups (no K-means clustering is run). A cross-validation with 1,000 replicates was performed for the three sub-analyses to retain an optimal number of PC-axes with the function xvalDapc from the R-package adegenet (Jombart 2008; Kamvar et al. 2015). We use scatterplots of the first four principal components of the DAPC analyses to visualize within and between population variation in the study area.

#### Assignment tests

To identify the most likely population of origin for all individuals based on the genetic profiles of these individuals and the populations, we performed individual assignment with GENECLASS2 tests to identify immigrant individuals or individuals that have recent immigrant ancestry (Rannala and Mountain 1997; Piry et al. 2004). The most standard method currently is to perform first-generation migrant detection (Paetkau et al. 2004). However, as this model is explicitly based on the sampling of gametes from haploid or diploid populations, we considered this method inappropriate for a haplodiploid mating system. Therefore, we used a Bayesian criterium to estimate likelihoods for each individual to originate from any of the given populations based on allele frequencies, combined with the probability computation from the same method using 10,000 Monte Carlo simulations and the expected type I error rate (alpha) set to 0.01 (Rannala and Mountain 1997). These probability computations are based on random drawing of alleles using allele frequencies directly estimated from the reference population samples and are thus mating system independent. With this method, we can identify individuals that are immigrant or have recent immigrant ancestry. However, interpretation should be done with care, as these assignment or exclusion methods are (compared to first-generation migrant detection) known to be prone to over-rejection of resident individuals and thus might overestimate gene flow (Paetkau et al. 2004; Piry et al. 2004). All individuals were considered and all populations were included as possible source populations. We made a flow chart (spatial directed network graph) in QGIS (QGIS Development Team 2020) representing the links between sampled populations and putative origin population according to the assignment tests. Arrows in such a graph start at the putative source population according to the assignment tests and end in the sampled population.

To check the robustness of the assignment analyses against skewed sample sizes, we repeated the assignment tests for a subsampled dataset (10 samples are randomly selected if a population has more than 10 samples and the two populations of sample size of 5 were omitted).

## 4. Results

In total, wing tips of 867 individuals from 53 populations were genotyped. Five microsatellite loci showed a lot of stutter in the amplification profiles which were hard to score and were therefore discarded from further analysis (AGBro486, −329, −196, −437, −298). Hardy-Weinberg (HW), linkage disequilibrium (LD) and null alleles assumption testing identified a further 7 microsatellites that were left out of the analysis (supplementary material S5): AGBro35, −57, −419 (HW and null alleles), AGBro111 (HW and LD), AGBro20, −16 (null alleles) and AGBro138 (HW, LD, null alleles). This resulted in a total of 21 microsatellite loci for further population genetics analyses. If an individual had more than 8 loci with missing data, it was discarded from the analysis beforehand (10.5%; 102 out of 969). A total of 133 alleles with an average of 6.3 alleles per locus (ranging from 3 to 14) were observed across the 21 microsatellite loci. The resulting dataset had an overall 3.48% of missing data for the 867 individuals.

Hierarchical AMOVA comparing genetic variation between populations and between years (for a dataset of 522 samples (of 867) from 23 populations sampled both in 2018 and 2020), showed that sampling year explained 0.19% of the variation (supplementary material S6). Thus, it was decided to pool the different years for the subsequent population genetics analyses (supplementary material S1).

Subsampling performed on several of the analyses (as sample sizes per population ranged between 5 and 35; supplementary material S1) showed that our results are robust for skewed sample sizes per population (details below).

### Population-level statistics

The complete table with number of private alleles (NP), rarefied allelic richness (AR), expected (H_e_) and observed (H_o_) heterozygosity and inbreeding (F_IS_) can be found in supplementary material S7. Table 1 gives a summary of these population-level statistics for coast and inland separately. Allelic richness, and expected and observed heterozygosity were in general lower in the inland populations (table 1). Inbreeding was in general high and very variable across all populations (table 1), which is expected for a haplodiploid system (Zayed 2004). From the 10 populations with the highest F_IS_, six were from the mid- and eastern part of the coast and four from inland populations, including the two Walloon populations (populations 52-53). The repeated analysis for the subsampled dataset yielded very similar results (supplementary material S7).

**Table 1:** summary table for the population-level statistics (Statistic): rarefied allelic richness (AR), expected (H_e_) and observed (H_o_) heterozygosity, and inbreeding coefficient (F_IS_). Summary calculations are for two regions: Coast (coastal France and coastal Flanders combined) and Inland (inland Flanders and inland Wallonia combined). For each statistic and region, the mean, standard deviation (SD), minimum (min) and maximum (max) of the range are given. To check the difference of a statistic between regions, a two-sided t-test was performed and t-value (t), degrees of freedom (df) and p-value are given.

| Statistic | Region | mean | SD | min | max | t | df | p-value |
| --- | --- | --- | --- | --- | --- | --- | --- | --- |
| AR | Coast | 2.61 | 0.09 | 2.41 | 2.84 | 6.21 | 14.84 | <0.001 |
|  | Inland | 2.33 | 0.15 | 2.11 | 2.61 |  |  |  |
| $H_e$ | Coast | 0.59 | 0.03 | 0.53 | 0.64 | 5.82 | 14.41 | <0.001 |
|  | Inland | 0.51 | 0.05 | 0.43 | 0.58 |  |  |  |
| $H_o$ | Coast | 0.52 | 0.06 | 0.36 | 0.62 | 3.50 | 19.38 | 0.002 |
|  | Inland | 0.45 | 0.07 | 0.33 | 0.54 |  |  |  |
| $F_{IS}$ | Coast | 0.11 | 0.09 | -0.02 | 0.36 | -0.18 | 20.60 | 0.86 |
|  | Inland | 0.12 | 0.09 | -0.03 | 0.27 |  |  |  |

### Genetic distance

Figure 2 shows the pairwise D_S_ (Nei’s standardized genetic distance) between all populations. The 10 populations with the highest mean D_S_, which have the highest average differentiation from all other populations, were inland populations, including the two Walloon populations (table S4.1). Similar figures for pairwise differentiation measures F_ST_ and Jost’s D can be found in supplementary material S4, giving similar results. Genetic distances were overall large and variable among inland populations (right upper corners Fig. 2) and small among coastal sites (left lower corner Fig. 2 and y-axis in figure 3). The genetic distances between coastal and inland regions are medium to large. The repeated analysis for the subsampled dataset yielded similar results (supplementary material S4). Extra hierarchical AMOVA’s for coastal and inland regions separately also confirmed there is more differentiation between populations inland than at the coast (supplementary material S6).

**Figure 2:**
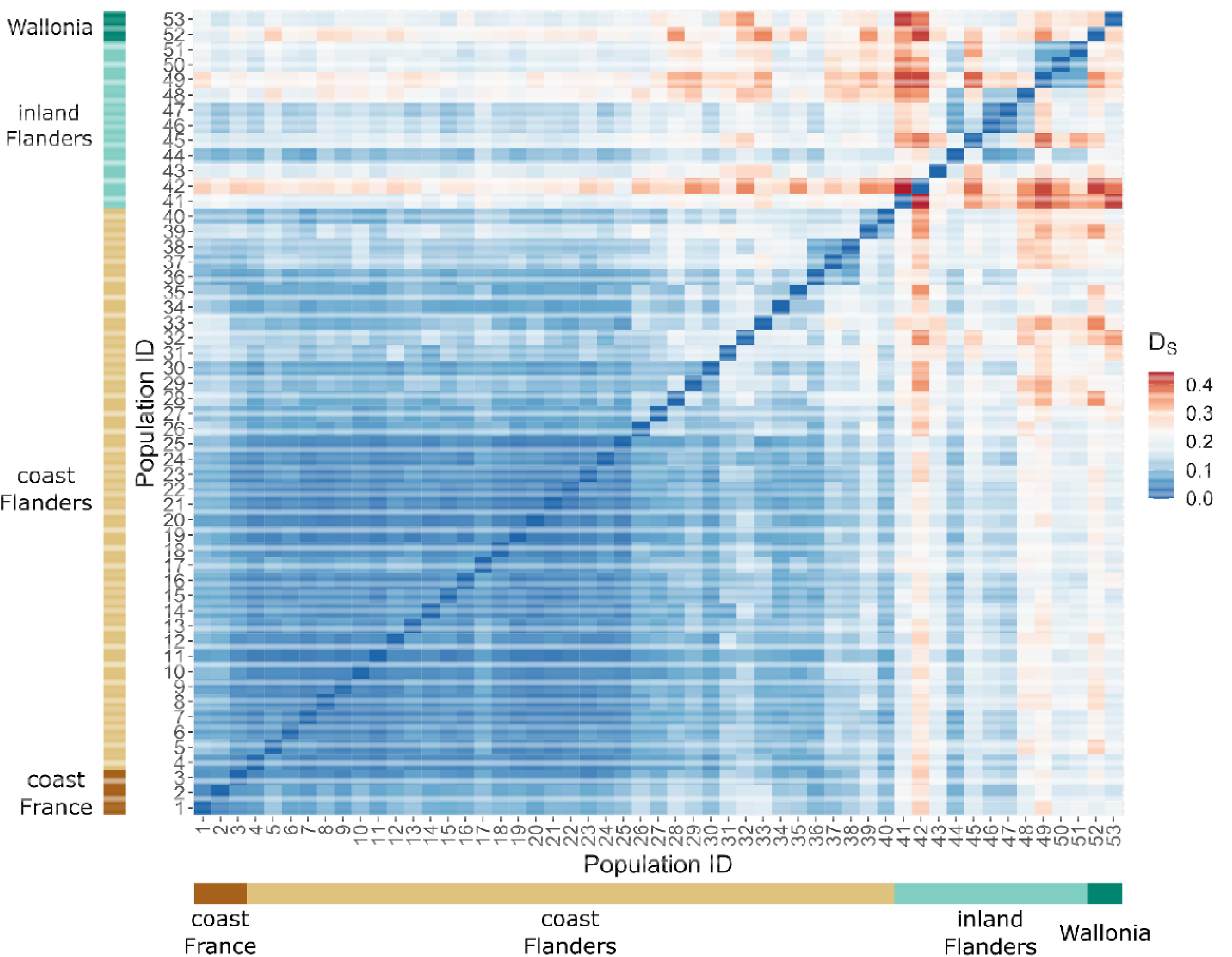
graphical matrix representation of Nei’s standardized genetic distance (D_S_): blue are low, white are mid, and red are high genetic distance values between pairwise populations. The x- and y-axes represent the population ID, subdivided in the four different regions. Genetic distances are symmetrical and consequently the matrix is mirrored along the diagonal. There are overall large genetic distances within the inland regions (right upper corner, populations 41-53) and small genetic distances within the coastal regions (left lower corner, populations 1-40). The genetic distances between coastal and inland regions (y-values 41-53 with x-values 1-40, or vice versa) are medium to large.

**Figure 3:**
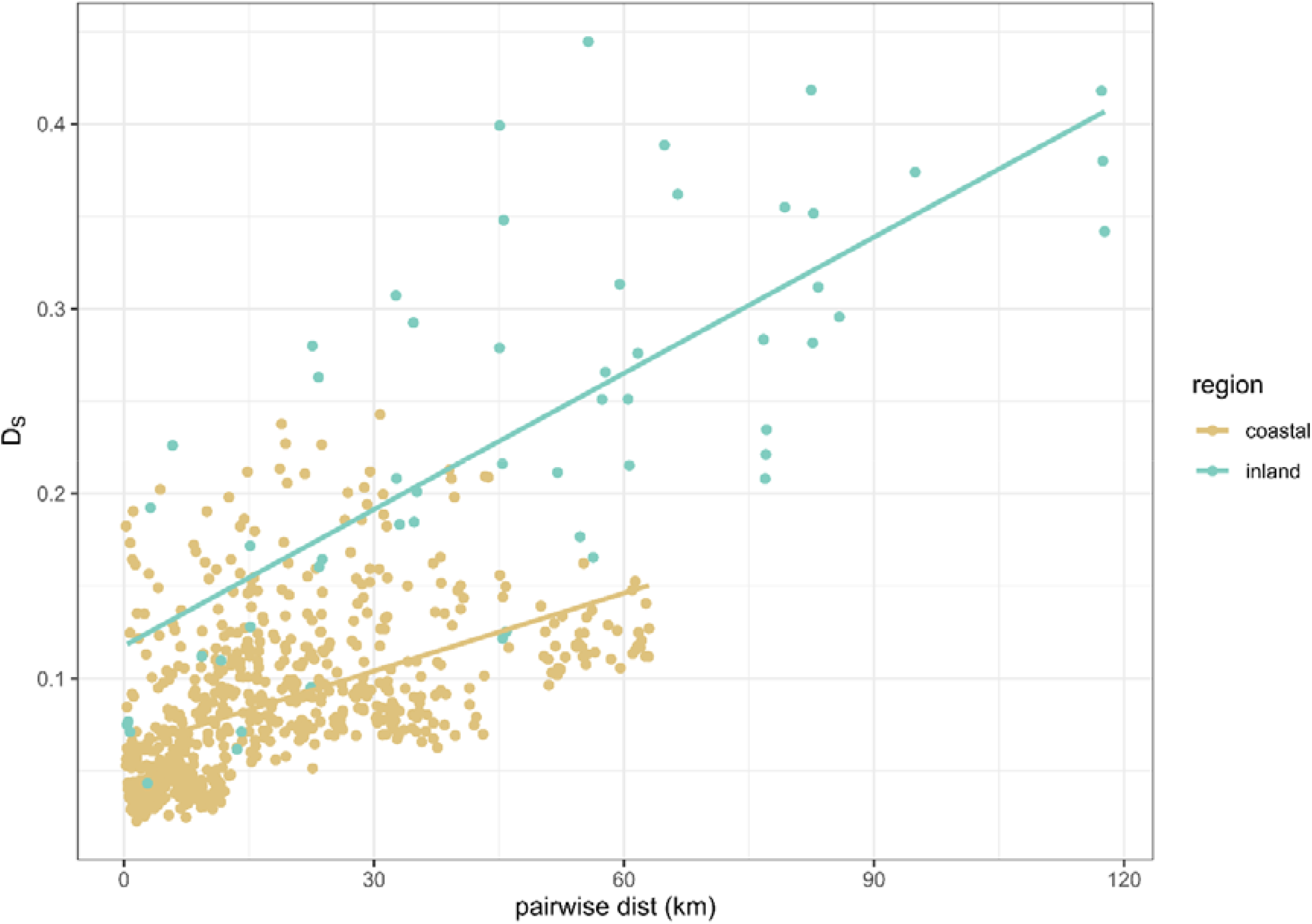
Isolation-by-distance (IBD) graph with Nei’s genetic distance (D_S_) plotted against Euclidian geographical distance (in km), separately for coastal and inland Flanders. Lines are smoothers plotted for indicating the trend, not regressions: statistical tests are done through RDA and a permutation test (see main text). These showed that the spatial genetic structure is higher for the inland regions and that both intercept and slope differ between the regions.

### Isolation-by-distance

Isolation-by-distance (IBD) is only calculated for coastal and inland Flanders as they have the most complete spatial coverage in sampling. An RDA performed with Nei’s genetic distance D_S_ and the pairwise geographical Euclidian distances indicate there is spatial genetic structure, and most strongly for the Flanders inland region: proportion explained by the constrained axes (or R²) is 22% for all populations together, 23% for the coastal region and 61% for the inland region. All explained variance is larger than zero (p=0.001; Df=2). Adjusted R² is 18%, 18% and 51% respectively. The relationship between D_S_ and geographical distance is shown in Figure 3. Both the intercept (region: p<0.001; SS=0.342 (Type III); Df=1) and slope (distance-region interaction: p<0.001; SS=0.088 (Type III); Df=1) differed significantly between regions according to the permutation test (adjusted R² of complete model was 58%). Some coastal datapoints (Fig. 3, in brown) are situated on the inland trendline. These are populations from the mid- and eastern part of the coast (populations 26-38; Fig. 1).

### Discriminant Analysis of Principal Components

Figure 4 gives scatterplots for the (Discriminant Analysis of Principal Components) DAPC analysis for the complete dataset. Scatterplots for the coastal and inland regions separately are given in supplementary material S8. The coastal Flanders populations (light brown in Fig. 4) clearly clump together, with coastal France (1-3, dark brown in Fig. 4) partially overlapping. A similar pattern can be seen in the DAPC for the coastal regions separately (Fig. S8.1A) and are in line with a pattern of isolation-by-distance. Inland populations show more between-populations structure (greens in Fig. 4; Fig. S8.1B). This is also confirmed by the separate analyses for both regions: for the coast, the first two principal components together explain 27.2% of the variation, while for inland this is 54.6%.

**Figure 4:**
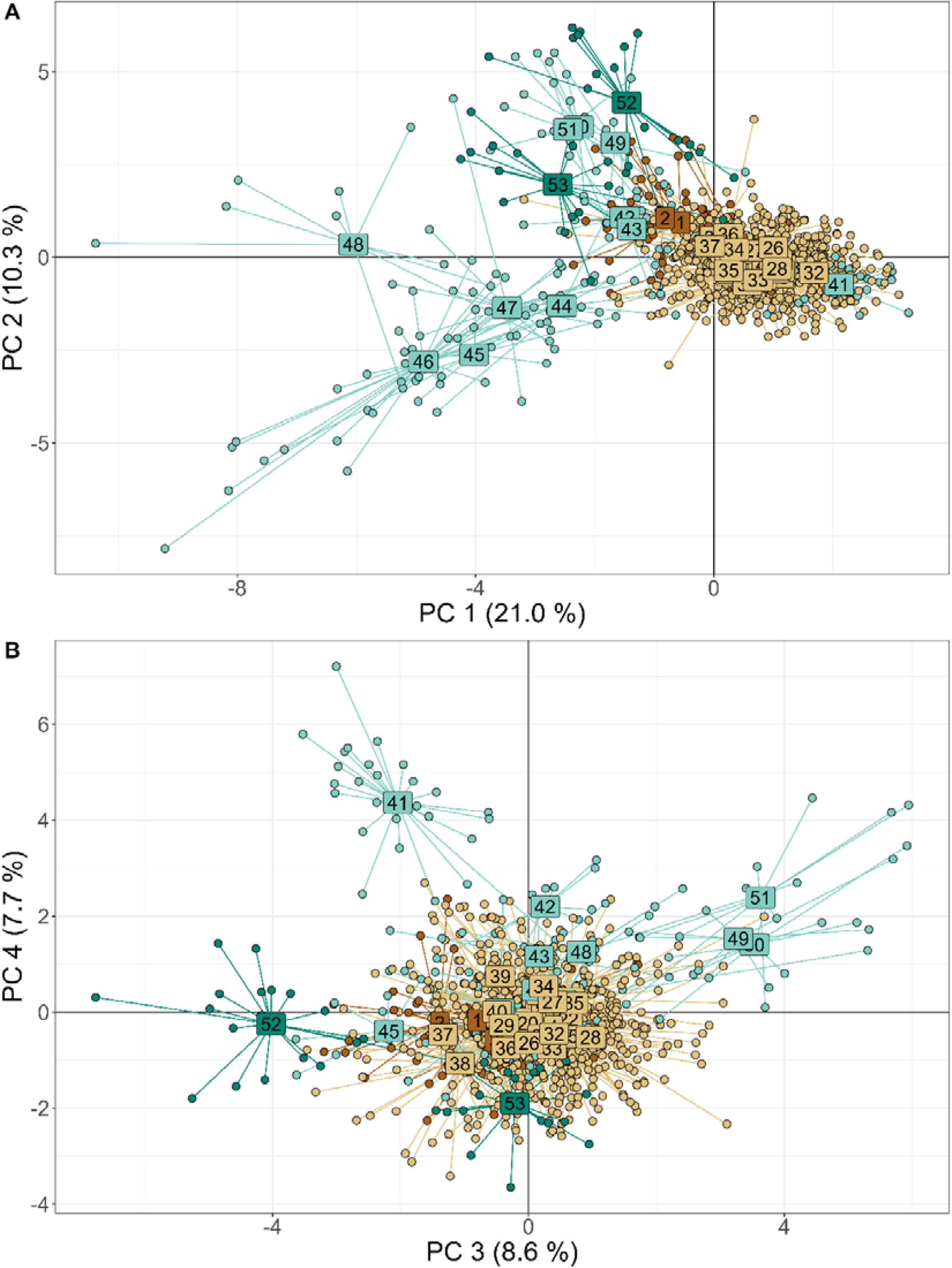
Scatterplot for DAPC results on complete dataset for A) the first two and B) the third and fourth principle components. Labels indicate population ID and point and label colors refer to the regions from figure 1: coastal France (dark brown), coastal Flanders (light brown), inland Flanders (light green), inland Wallonia (dark green). Coastal populations cluster together genetically in a large point cloud while inland regions show more genetic differentiation.

### Assignment tests

Figure 5 summarizes results of all the assignment tests per region and figure 6 depicts derived flow charts for genetic links between regions and within the inland region (not for within the coastal region as these are very numerous; Fig. 5). The assignments depict immigrants or individuals with recent immigration ancestry, probably up to two generations (Rannala and Mountain 1997), i.e. not pure first generation migrants as in Paetkau et al. (2004).

**Figure 5:**
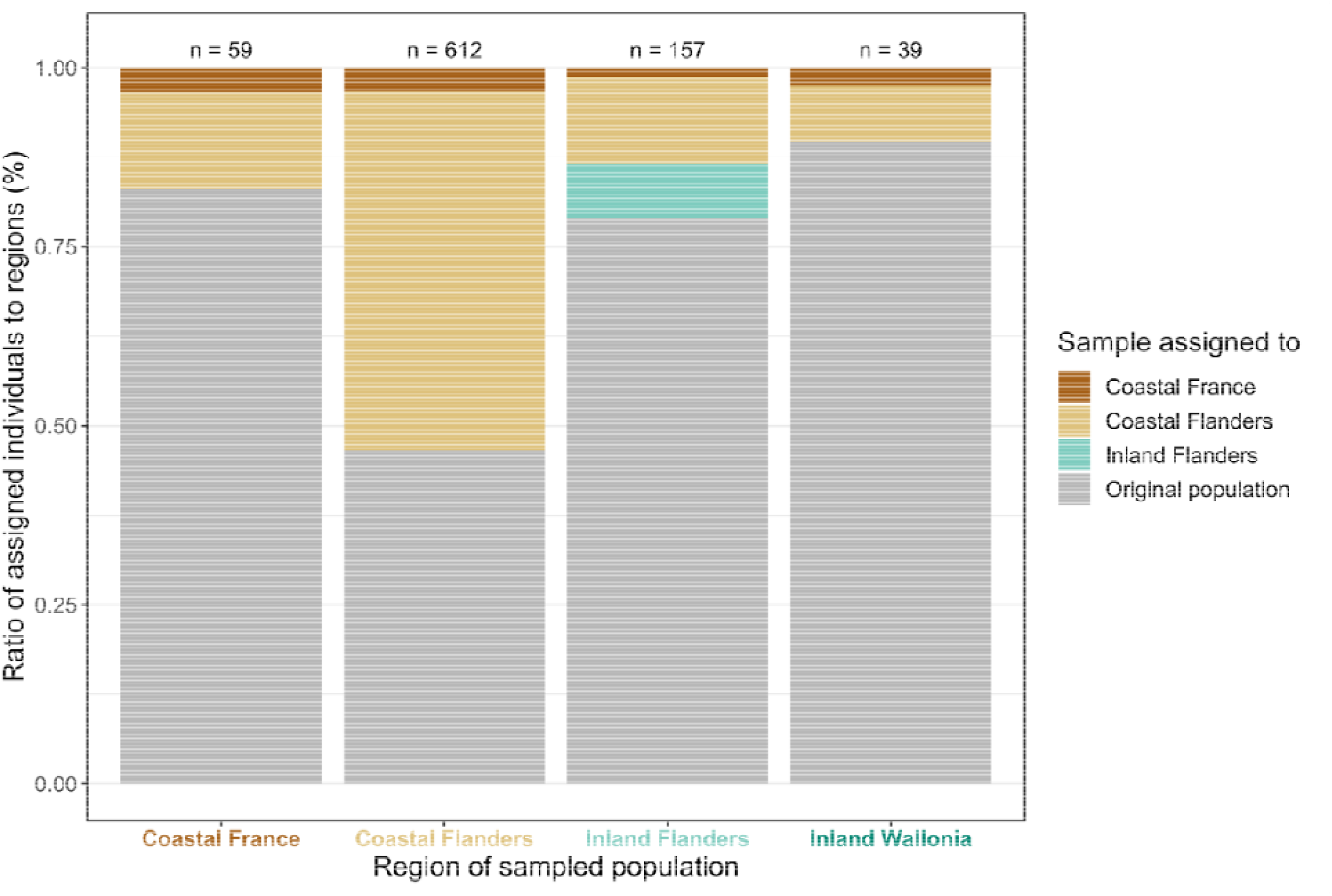
barplot of summarized results of the assignment tests for all populations from each sampled region (x-axis). If an individual was assigned to its original population where it was sampled, the barplot-area is filled with grey. If an individual was assigned to another population within the same region or to another region, the barplot-area is colored by region (dark brown: assigned to coastal France, light brown: assigned to coastal Flanders, light green: assigned to inland Flanders). Total number of samples per region (n) is indicated above each barplot. Apart from samples assigned to their original population, there were no other individuals assigned back to inland Wallonia (would have been dark green colored in barplot). Coastal Flanders has the least samples assigned back to their original populations. However, most were assigned within the region, indicating high genetic connectivity within coastal Flanders. If samples were assigned to another region, they were always from coastal regions (brown colours; Fig. 6).

**Figure 6:**
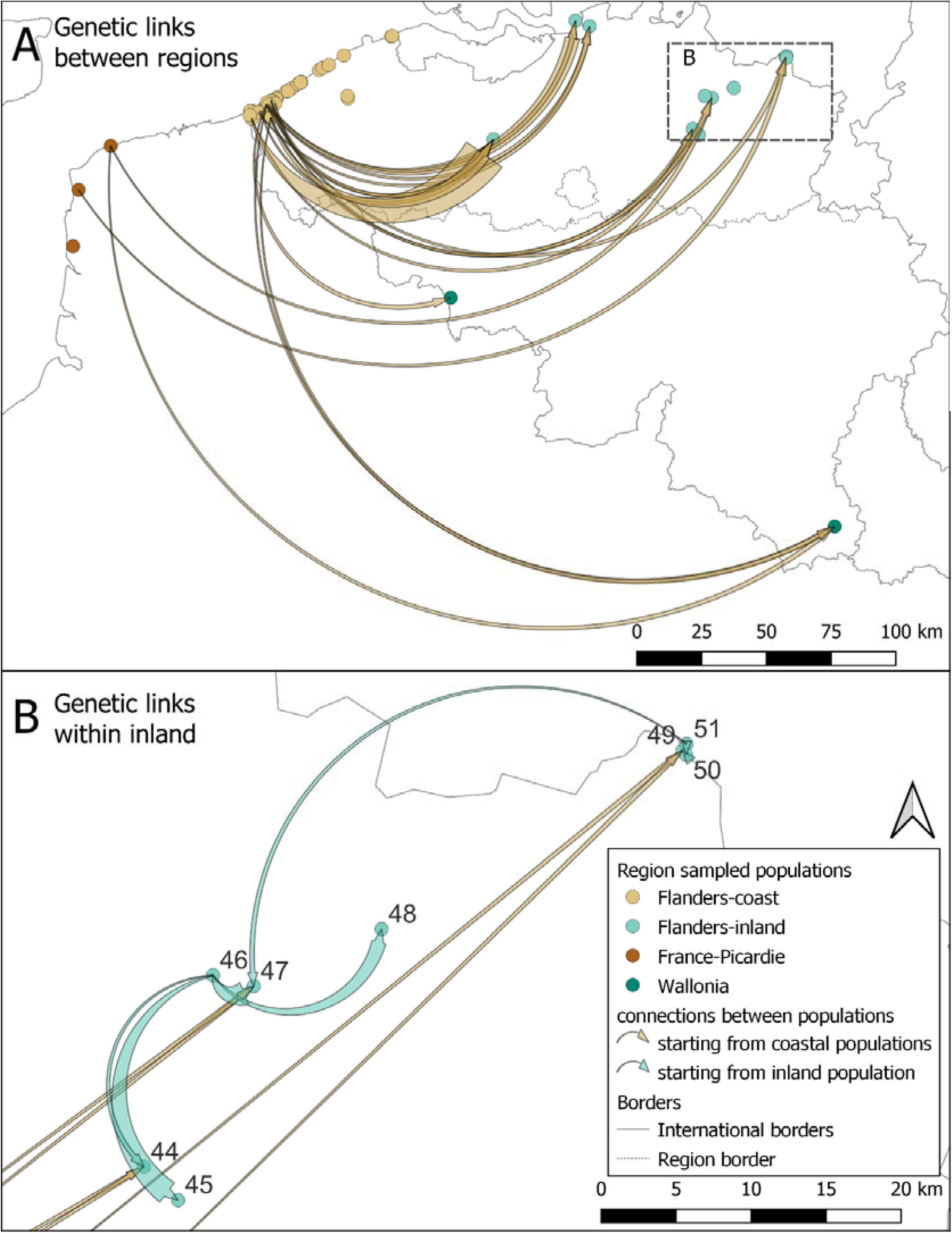
flow chart of genetic links (A) between regions and (B) within the inland region for B. rostrata according to assignment to putative source populations. Genetic links within the coastal region are not depicted as these were too numerous (Fig. 5). The links represent the number of individuals assigned to a putative origin population (start of the arrow) that were caught in the sampled population (end of the arrow). Brown arrows are links starting from the coast, green arrows start from inland populations. The thicker the end of the arrow, the higher the number of individuals assigned to the putative source. Genetic links are present from coast to inland, but not from inland to coast (A; fig. 5). Within the inland region, there are only genetic links within the cluster of populations 44 to 50 (B). The main source within inland Flanders is population 46 (Geel-Bel), which is the largest and oldest known inland population in Flanders. Individuals from other inland populations (41-43; 52, 53) are either assigned to their sampled population or are assigned to a coastal population. These source populations are predominantly from the west of coastal Flanders (A).

Three main patterns can be deduced from the assignment tests: high genetic connectivity at the coast, restricted genetic links within inland, and asymmetrical gene flow from coast to inland. Within coastal Flanders, genetic connectivity among populations is substantial (Fig. 5): a relative low number of individuals are assigned to their original sampled population (47%) but almost all are assigned within the region (97%). Within the coastal Flanders populations, populations from the west coast are the largest source of gene flow to populations at the mid- and east coast. Individuals from coastal France are mainly assigned to the region itself, but there is connectivity with coastal Flanders in both directions (Fig. 5). Inland regions have relatively high numbers of individuals assigned to the original sampled populations (79% and 90%) compared to coastal Flanders (47%). However, they also have a higher number of individuals assigned to another region (13% and 10% compared to 3% in coastal Flanders), mainly coming from the western part of coastal Flanders and France (Fig. 5 and 6). The only genetic connectivity present within the inland populations is a cluster in east Flanders (populations 44-52; Fig. 5B), wherein population 46 (Geel-Bel) seems to be a central source population for surrounding population. Thus, inland populations are genetically more isolated from each other, but there is a genetic influx from the coast, which seems to happen unidirectional from coast to inland (Fig. 5 and 6). The results of the assignment tests with the subsampled dataset are similar but have some minor differences (supplementary material S9). Nevertheless, the main conclusions—gene flow high within coast, restricted inland, asymmetrical from coast to inland— remain robustly present.

The pairwise geographical distances between sampled and assigned populations show that 75% of the distances between sampled and putative source populations lie below 20 km, with the largest distance being 320 km from coastal France to the south of Belgium (supplementary material S9, Fig. S9.3). The largest distance within Flanders between a putative source coastal population and sampled inland population is 201 km.

## 5. Discussion

Bembix rostrata populations from inland sandy regions exhibited low levels of gene flow, low genetic diversity, and high genetic differentiation, contrary to the coastal region, which has an overall high level of genetic connectivity. Asymmetrical gene flow from the coast to inland demonstrate that the species is—contrary to expectations based on its behavior and poor colonization capacity—capable of dispersing to existing populations at a distance of 200 to 300 km.

Bembix rostrata has always been considered to be a philopatric species, not able to easily colonize vacant habitat (Nielsen 1945; Larsson 1986). The retrieved pattern of genetic structure and gene flow within and across sandy regions of Belgium clearly demonstrates this does not prevent gene flow between already existing populations. The species is known to be gregarious, with on the one hand local nest choice behavior showing positive density dependence because of conspecific attraction, and on the other hand individuals making consecutive nests close to one another (Larsson 1986; Batsleer et al. 2022a). This intragenerational individual site fidelity combined with the reported low colonization capacity (Nielsen 1945; Bogusch et al. 2021), made this species a presumed poor disperser. Our results show that dispersal between existing populations is not highly restricted, especially in a well-connected, stepping stone landscape, such as in the Belgian coastal dunes. Likely, female colonization capacities are not restricted by the species’ movement capacity, but mainly by the settlement phase of dispersal—with conspecific attraction of crucial importance in B. rostrata (Batsleer et al. 2022a). When conspecifics are already present in existing populations, the settlement phase is less restricted, which can explain the pattern of gene flow between existing populations.

Alternatively, as colonization capacity in B. rostrata is clearly disconnected from gene flow, the latter may be equally or largely driven by male dispersal. Male-biased dispersal has indeed been found to be most common in other bees and wasps (Johnstone et al. 2012). Given the protandry of the species, such dispersal may be common in the period prior to female emergence, as a strategy to avoid strong (kin) competition (Bonte et al. 2012; Baguette et al. 2013). As we only sampled females, we cannot quantify a biased dispersal strategy, with for instance genetic spatial autocorrelation analyses (Banks and Peakall 2012). Because of the haplodiploid mating system, indirect analyses by the comparison of nuclear and mitochondrial markers are neither suitable because nuclear introgression is reduced relative to mitochondrial introgression (Patten et al. 2015).

Effective dispersal rates, resulting in establishment, depend on the species’ (i.e. female) capacity to move and settle, but also on the size of the source population. Colonization at short distance of vacant, newly emerging pioneer habitats remains overall much more likely than distant colonization. In addition, during periods of exceptionally suitable environmental conditions, e.g. warm summers or years with high resource abundance (nectar, prey), any local overshooting of carrying capacities may further increase the magnitude of gene flow, both in terms of extent (the threshold in density dependent dispersal; Kun and Scheuring 2006; Best et al. 2007) and spatial scale (the fatness of the dispersal kernel; Bitume et al. 2013). We hence hypothesize that the establishment of new populations may only succeed when Allee-effects are overcome by the simultaneous settlement of multiple females into a single cluster. Such demographic contributions are often overlooked in the dynamics of spatially structured populations in both population genetics and connectivity studies (Lowe et al. 2017; Drake et al. 2022). With our genetic data, we could not detect if a population was recently established or not, but at least for one population (Averbode, population 44 in Fig. 1) we know the area was only recently made suitable (ca. 2010). For this population, but also other nearby populations, assignments tests indicate that genetic connections mainly originate from large (meta)populations at the west coast and within inland from one nearby population inland (Geel-Bel, population 46, Fig. 6B). Dunes from the west coast (populations 1-25) are known to hold the highest number of old and large (meta)populations of B. rostrata in Belgium (unpubl. monitoring data; Klein and Lefeber 2004; confirmed by citizen science data from observation.org, and confirmed to a certain degree by F_IS_ results, supplementary material S7), while Geel-Bel is the single largest and oldest known population in the inland region. This indirectly confirms the very often neglected role of source population size compared to absolute dispersal potential (the dispersal kernel) for gene flow in metapopulations and their dynamics. In larger populations, the absolute number of dispersers will be larger, even if per-capita dispersal rate is constant for different population sizes. The important role of demographics could also explain the observations of Bogusch et al. (2021), who observed highly restricted local and regional colonization abilities of two small and isolated populations in the Czech Republic.

The strong impact of the size of source populations on gene flow likely also underlies the general asymmetrical gene flow from coastal to inland populations. In the well-connected, stepping stone landscape at the (west) coast, short-distance dispersal—dispersal related to routine movements of resource exploitation (Van Dyck and Baguette 2005)—results in a pattern of weak isolation-by-distance. As the genetic population structure is dominated by such short-distance dispersers, the proportionally low numbers of long-distance dispersers do not leave a detectable genetic signal within the coastal network. However, some long-distance dispersers–males or females–from the coast appear to reach inland populations and leave a proportionally larger signal of gene flow in these smaller populations. In addition, also within regions, such demographic signals are picked-up. First, populations from the west coast are the largest source of gene flow to populations at the mid- and east coast, known to hold fewer and smaller populations (unpubl. monitoring data; confirmed by F_IS_ values, supplementary material S7). Second, as mentioned previously, the largest and oldest known inland population (Geel-Bel, population 46) leaves the strongest genetic signal in the surrounding inland populations (Fig. 6B).

In addition to the demographic causes, other mutually non-exclusive mechanisms may underlie the retrieved pattern of asymmetrical gene flow. Although we consider it less likely in our case, wind has been put forward as a dominant factor for long-distance flight behavior and migration in insects (Alerstam et al. 2011; Knight et al. 2019; Leitch et al. 2021). In the focal study area, the predominant wind-direction is from coast to inland (SW and WSW), which could reinforce the demographic process at the coast. Alternatively, dispersal capacity itself could be more restricted in inland regions as well. Indeed, our isolation-by-distance results suggest that apart from the intercept, the slope differs between the regions as well. The intercept is related to population sizes (N): if N decreases, all else being equal, differentiation will increase due to genetic drift and a lower total number of migrants. The steeper slope could be—apart from the influence of spatial habitat configuration (van Strien et al. 2015)—due to dispersal capacity being more restricted in the inland region. A difference in dispersal capacity within a species can result from evolved dispersal reductions in highly fragmented landscapes (Cheptou et al. 2017) where costs of dispersal are highest or where spatio-temporal patch turnover is lowest (Bowler and Benton 2005; Bonte et al. 2012; Duputié and Massol 2013). Such conditions may indeed be more prominent for the more fragmented and locally stable populations from the inland sandy regions. Only a combination of behavioral experiments and/or quantifying physiological differences in flight metabolic performance may shed light on the likelihood of such processes (Hanski et al. 2004).

When not all possible source populations are sampled and included, assignment tests might give rise to misleading results (Rannala and Mountain 1997; Cornuet et al. 1999). In our sampling design, sampling in Flanders covered all known populations at the time and the main known large populations in Wallonia. Nevertheless, unsampled potential source populations might be present across the border in the Netherlands and France. Consequently, the populations from northern inland Flanders (populations 42, 43, 49-50) could be connected to Dutch populations and not be as isolated as our results suggest. Especially for the connections from coastal France to inland (Fig. 6), intermediate populations might be present (Bitsch et al. 1997; Barbier 2007). Therefore, the detected connections between coastal France and inland regions might not be from directly dispersing individuals, but through an indirect connection of an unsampled French population. If this is the case, the maximum distance from a direct connection within Flanders would be 201 km instead of 320 km. A second bias that can arise with the specific assignment method we used—a method suitable for haplodiploids—has been detected with a simulation study: the possibility over-rejection of resident individuals, ultimately overestimating gene flow (Paetkau et al. 2004; Piry et al. 2004). Considering our results, the absolute number of genetic links might be lower and the maximum dispersal distance an overestimation. However, as the absolute number of genetic links is also dependent on sample sizes and number of genetic markers, our interpretations are essentially relative and will still hold: more restricted gene flow in inland populations than at the coast and asymmetrical gene flow from coast to inland.

Pollinator conservation, and more specifically that of wild bees, is currently a major topic of interest to policy and science (Potts et al. 2016). A major inherent factor complicating the interpretation of population genetics results of hymenopteran species in a classical conservation genetics framework is the haplodiploid mating system. Males are haploid (unfertilized) and females are diploid, which results in non-symmetrical inheritance of genes across generations. In such a mating system, inbreeding coefficients will be inherently high and effective population sizes low due to, for instance, purging effects on deleterious alleles in haploid males (Zayed 2004, 2009). These specific attributes render metrics based on assumptions of HW-equilibrium and population genetics models difficult to apply and interpret, as different genetic processes will predominate in a haplodiploid conservation genetics framework (Zayed 2009). While classical population genetic analyses may be used if not overinterpreted (Černá et al. 2013; Sanllorente et al. 2015), we decided to report mating system independent analyses, such as a multivariate approach DAPC and classical assignment tests. The descriptive statistics provided should be interpreted with care and only be considered relatively within the study system. In our opinion, future (modelling) studies should further elucidate the potential biases for haplodiploid systems when using classical population genetics studies that are based on diploid mating systems and related assumptions. In general, integration of haplodiploids in the conservation genetics framework is lacking, although about 15% of all animal species are haplodiploid (Evans et al. 2004; Lohse and Ross 2015).

Connectivity remains difficult to generalize among species and even for different landscapes within a single species. In Flanders, the genetic connectivity of the grayling butterfly (Hipparchia semele)—also occurring in sandy habitats—was in general much more restricted and gene flow slightly higher in the inland region compared to the coast (De Ro et al. 2021). Differences in life history traits and niche are the most likely reason for the contrasting results with B. rostrata. These diverging findings for the focal region between two species from sandy habitat stress that connectivity is a trait of both species and population configuration combined—and should as such be considered in conservation policies. Additionally, for B. rostrata itself, the asymmetrical gene flow and discrepancy in connectivity between different regions were important to discuss the disconnection of gene flow from colonization capacity and consider the plausible role of demographics for the observed genetic connectivity. As such, it is crucial to combine and compare results from different regions for a single species to fully understand possible mechanisms of gene flow. It remains to be tested whether our insights on the species metapopulation structure can be scaled up towards the species’ full range. The general insight that the species’ low colonization capacity does not imply low levels of gene flow are likely to hold across other well-connected, healthy and large (meta)populations. Nature management implications discussed below are potentially helpful for B. rostrata populations across Europe, depending on the local and regional context.

### Conservation implications

Our findings have direct implications for nature management and conservation of the flagship insect species B. rostrata at both local and landscape scale. At the coast, a well-connected metapopulation occurs, while inland populations show restricted gene flow in a fragmented sandy habitat landscape. Moreover, there is an asymmetrical genetic influx from coast to inland, which we mainly interpret as being linked to the larger population sizes at the coast. The species’ poor colonization capacity, resulting in a low establishment probability, should be considered disconnected from gene flow between existing populations, as the latter seems much less restricted. To maintain the well-connected, large coastal populations, conservation should focus on local management and internal processes to ensure a constant amount of suitable habitat through time. Ideally, dunes are revitalized with aeolian (wind) dynamics at the landscape level (Provoost et al. 2011). However, in a fragmented and urbanized landscape, the current management framework focuses on grazing used as a tool to locally revitalize sand dynamics (Provoost et al. 2004). It is recommended to use a heterogenous approach for grazing and grazer type in space and time to reconcile short-term negative (trampling) and long-term positive (open dune landscape) effects of grazing on B. rostrata (Bonte 2005; Batsleer et al. 2022b). The genetically well-connected landscape and large metapopulation context ensures population recovery and persisting connectivity when implementing such a dynamic management approach.

Contrastingly, more isolated, small populations such as in the inland region, need a more cautious approach and management should consequently focus on the protection of individual populations or clusters of nearby populations. Creating extra stepping stones to increase landscape connectivity, which may already be partially present but vacant, might be less effective in the current context, as potential gene flow is not highly restricted in this species. We suggest that the primary focus should be on enlarging (source) population sizes by improving the quality of the local and directly surrounding habitat. This can be achieved by maintaining or creating open, pioneer sand dune habitat. Preventing or removing encroachment preferably happens manually, as grazers should be used with caution in small, isolated populations (Batsleer et al. 2022b). Apart from nesting resources, sufficient neighboring floral resources for both nectar and prey hunting may also be important to sustain large populations of B. rostrata (Kimoto et al. 2012; Buckles and Harmon-Threatt 2019).

## Supporting information

Supplementary materials S5 - output visualisation-HW-LD-NA

## Acknowledgements

We thank the following persons and instances for permission and access to nature reserves: Johan Lamaire, Guy Vileyn, Koen Maertens, Evy Dewulf and Klaar Meulebrouck from ANB (Agency for Nature and Forests – Flemish government); Bruno Nicolas from Eden 62 (France); Thierry Paternoster from DEMNA (Dép. de l’Étude du Milieu Naturel et Agricole - Service public de Wallonie); Rika Driessens from IWVA/Aquaduin; Rudi Delvaux from Grenspark Kalmthoutse Heide; Griet Limet from Kempens Landschap. We thank Pieter Vanormelingen from Natuurpunt to update us on the latest observations in new areas of B. rostrata. We also thank Maarten Jacobs (Sjacky) and all other friendly volunteers from Averbode Bos-en Heide (Natuurpunt) for help with searching and sampling B. rostrata in and around their nature reserve. We also than following people for assistance during sampling: Margaux Boeraeve, Ward Tamsyn, Marc Batsleer, Nadine De Schrijver, Pepijn Boeraeve and Ward Langeraert. We thank Viki Vandomme for assistance during lab work.

We thank 2 anonymous reviewers for their detailed and constructive comments that greatly improved the manuscript. We also thank Jan Van Uytvanck, Laurence Cousseau, Viktoriia Radchuk and Josep D. Asís for additional comments.

Permit numbers for collection of genetic materials and entrance of nature reserves in the study area: TREL2022990S/383 (Republique France – Ministère de la transition écologique; according to Nagoya protocol); DNF/DNEV/JPB/SLA/2020-RS-22 (Service public Wallonie - Département de la Nature et des Forêts); N69.21232 (La Défense – Direction Générale Material Resources; access to military domain in Wallonia). Permission to enter governmental nature reserves in Flanders was granted by ANB (Agency for Nature and Forests).

F.B. was supported by Research Foundation – Flanders (FWO).

## Author contributions

FB, DD, JM, AVB, DM, DB contributed to the study conception and design. Data collection was performed by FB, MG, MD, PV, FT; laboratory work and data analyses were performed by FB, MG, MD. The first draft of the manuscript was written by FB and all authors commented on previous versions of the manuscript. All authors read and approved the final manuscript.

## Data availability

Scripts and data are made available on github (https://github.com/FemkeBatsleer/PopGenBembix.git) for review and will be published on zenodo upon acceptance.

## Conflict of Interest

The authors have no conflict of interests.

## Notes

https://github.com/FemkeBatsleer/PopGenBembix.git

### Competing Interest Statement

The authors have declared no competing interest.

### Summary of Updates

Peer-review comments were incorporated

## References

Adamack AT, Gruber B (2014) PopGenReport: simplifying basic population genetic analyses in R. Methods Ecol Evol 5:384–387. https://doi.org/10.1111/2041-210X.12158

Alerstam T, Chapman JW, Bäckman J, et al (2011) Convergent patterns of long-distance nocturnal migration in noctuid moths and passerine birds. Proc R Soc B 278:3074–3080. https://doi.org/10.1098/rspb.2011.0058

Baguette M, Blanchet S, Legrand D, et al (2013) Individual dispersal, landscape connectivity and ecological networks: Dispersal, connectivity and networks. Biol Rev 88:310–326. https://doi.org/10.1111/brv.12000

Banks SC, Peakall R (2012) Genetic spatial autocorrelation can readily detect sex-biased dispersal. Mol Ecol 21:2092–2105. https://doi.org/10.1111/j.1365-294X.2012.05485.x

Barbier Y (2007) Bembix rostrata (L.) (Hymenoptera, Crabronidae) de retour en Wallonie (Belgique). Osmia 1:5–6. https://doi.org/10.47446/OSMIA1.2

Batsleer F, Maes D, Bonte D (2022a) Behavioral Strategies and the Spatial Pattern Formation of Nesting. Am Nat 199:E15–E27. https://doi.org/10.1086/717226

Batsleer F, Maes D, Uytvanck JV, et al (2021) The difficult balance between dune management and protection of the digger wasp Bembix rostrata. Can grazing in the dunes be reconciled with the conservation of invertebrates? [in Dutch]. Natuurfocus 20:100–108

Batsleer F, Van Uytvanck J, Lamaire J, et al (2022b) Rapid conservation evidence for the impact of sheep grazing on a threatened digger wasp. Insect Conserv Diversity 15:149–156. https://doi.org/10.1111/icad.12532

Best AS, Johst K, Münkemüller T, Travis JMJ (2007) Which species will succesfully track climate change? The influence of intraspecific competition and density dependent dispersal on range shifting dynamics. Oikos 116:1531–1539. https://doi.org/10.1111/j.0030-1299.2007.16047.x

Bitsch J, Barbier Y, Gayubo SF, et al (1997) Hyménoptères Sphecidae d’Europe Occidentale, Volume II. Fédération Française des Sociétés des Sciences Naturelles, Paris

Bitume EV, Bonte D, Ronce O, et al (2013) Density and genetic relatedness increase dispersal distance in a subsocial organism. Ecol Lett 16:430–437. https://doi.org/10.1111/ele.12057

Blösch M (2000) Die Grabwespen Deutschlands. Goecke & Evers, Keltern

Bogusch P, Heneberg P, Šilhán K (2021) What are the main factors limiting the distribution of Bembix rostrata (Hymenoptera: Crabronidae) at early-succession sites? J Insect Conserv 25:571–583. https://doi.org/10.1007/s10841-021-00324-9

Bonte D (2005) Anthropogenic induced changes in nesting densities of the dune-specialised digger wasp Bembix rostrata (Hymenoptera: Sphecidae). Eur J Entomol 102:809–812. https://doi.org/10.14411/eje.2005.114

Bonte D, Maes D (2008) Trampling affects the distribution of specialised coastal dune arthropods. Basic Appl Ecol 9:726–734. https://doi.org/10.1016/j.baae.2007.09.008

Bonte D, Van Dyck H, Bullock JM, et al (2012) Costs of dispersal. Biol Rev 87:290–312. https://doi.org/10.1111/j.1469-185X.2011.00201.x

Bowler DE, Benton TG (2005) Causes and consequences of animal dispersal strategies: relating individual behaviour to spatial dynamics. Biol Rev 80:205–225. https://doi.org/10.1017/S1464793104006645

Buckles BJ, Harmon-Threatt AN (2019) Bee diversity in tallgrass prairies affected by management and its effects on above- and below-ground resources. J Appl Ecol 56:2443–2453. https://doi.org/10.1111/1365-2664.13479

Cardoso P, Barton PS, Birkhofer K, et al (2020) Scientists’ warning to humanity on insect extinctions. Biol Conserv 242:108426. https://doi.org/10.1016/j.biocon.2020.108426

Černá K, Straka J, Munclinger P (2013) Population structure of pioneer specialist solitary bee Andrena vaga (Hymenoptera: Andrenidae) in central Europe: the effect of habitat fragmentation or evolutionary history? Conserv Genet 14:875–883. https://doi.org/10.1007/s10592-013-0482-y

Châline N, Ratnieks FLW, Raine NE, et al (2004) Non-lethal sampling of honey bee, Apis mellifera, DNA using wing tips. Apidologie 35:311–318. https://doi.org/10.1051/apido:2004015

Chapuis M-P, Estoup A (2007) Microsatellite Null Alleles and Estimation of Population Differentiation. Molecular Biology and Evolution 24:621–631. https://doi.org/10.1093/molbev/msl191

Cheptou P-O, Hargreaves AL, Bonte D, Jacquemyn H (2017) Adaptation to fragmentation: evolutionary dynamics driven by human influences. Phil Trans R Soc B 372:20160037. https://doi.org/10.1098/rstb.2016.0037

Cornuet J-M, Piry S, Luikart G, et al (1999) New Methods Employing Multilocus Genotypes to Select or Exclude Populations as Origins of Individuals. Genetics 153:1989–2000. https://doi.org/10.1093/genetics/153.4.1989

De Keersmaeker L, Onkelinx T, De Vos B, et al (2015) The analysis of spatio-temporal forest changes (1775–2000) in Flanders (northern Belgium) indicates habitat-specific levels of fragmentation and area loss. Landsc Ecol 30:247–259. https://doi.org/10.1007/s10980-014-0119-7

De Ro A, Vanden Broeck A, Verschaeve L, et al (2021) Occasional long-distance dispersal may not prevent inbreeding in a threatened butterfly. BMC Ecol Evo 21:224. https://doi.org/10.1186/s12862-021-01953-z

Decleer K (2007) Europees beschermde natuur in Vlaanderen en het Belgisch deel van de Noordzee. Habitattypen | Dier- en plantensoorten. Mededelingen van het Instituut voor Natuur- en Bosonderzoek INBO.M.2007.01. Instituut voor Natuur- en Bosonderzoek, Brussels

Didham RK, Basset Y, Collins CM, et al (2020) Interpreting insect declines: seven challenges and a way forward. Insect Conserv Divers 13:103–114. https://doi.org/10.1111/icad.12408

Diniz-Filho JAF, Soares TN, Lima JS, et al (2013) Mantel test in population genetics. Genet Mol Biol 36:475–485. https://doi.org/10.1590/S1415-47572013000400002

Drake J, Lambin X, Sutherland C (2022) The value of considering demographic contributions to connectivity: a review. Ecography 2022:. https://doi.org/10.1111/ecog.05552

Duputié A, Massol F (2013) An empiricist’s guide to theoretical predictions on the evolution of dispersal. Interface Focus 3:20130028. https://doi.org/10.1098/rsfs.2013.0028

European Environment Agency, Schwick C, Madriñán L, Kienast F (2011) Landscape fragmentation in Europe: joint EEA-FOEN report. Publications Office of the European Union, Luxembourg

Evans HE, O’Neill KM (2007) The sand wasps: natural history and behavior. Harvard University Press, Cambridge, MA, United States

Evans JD, Shearman DCA, Oldroyd BP (2004) Molecular basis of sex determination in haplodiploids. Trends Ecol Evol 19:1–3. https://doi.org/10.1016/j.tree.2003.11.001

Exeler N, Kratochwil A, Hochkirch A (2008) Strong genetic exchange among populations of a specialist bee, Andrena vaga (Hymenoptera: Andrenidae). Conserv Genet 9:1233–1241. https://doi.org/10.1007/s10592-007-9450-8

Field J, Gonzalez-Voyer A, Boulton RA (2020) The evolution of parental care strategies in subsocial wasps. Behav Ecol Sociobiol 74:78. https://doi.org/10.1007/s00265-020-02853-w

Goudet J, Jombart T (2020) hierfstat: Estimation and Tests of hierarchical F-Statistics

Grünwald NJ, Goss EM (2011) Evolution and Population Genetics of Exotic and Re-Emerging Pathogens: Novel Tools and Approaches. Annu Rev Phytopathol 49:249–267. https://doi.org/10.1146/annurev-phyto-072910-095246

Haddad NM, Brudvig LA, Clobert J, et al (2015) Habitat fragmentation and its lasting impact on Earth’s ecosystems. Sci Adv 1:e1500052. https://doi.org/10.1126/sciadv.1500052

Hanski I, Erälahti C, Kankare M, et al (2004) Variation in migration propensity among individuals maintained by landscape structure. Ecol Lett 7:958–966. https://doi.org/10.1111/j.1461-0248.2004.00654.x

Hanski I, Moilanen A, Gyllenberg M (1996) Minimum Viable Metapopulation Size. Am Nat 147:527–541. https://doi.org/10.1086/285864

Hanski I, Ovaskainen O (2002) Extinction Debt at Extinction Threshold. Conserv Biol 16:666–673. https://doi.org/10.1046/j.1523-1739.2002.00342.x

Harvey JA, Heinen R, Armbrecht I, et al (2020) International scientists formulate a roadmap for insect conservation and recovery. Nat Ecol Evol 4:174–176. https://doi.org/10.1038/s41559-019-1079-8

Hodgson JA, Randle Z, Shortall CR, Oliver TH (2022) Where and why are species’ range shifts hampered by unsuitable landscapes? Glob Chang Biol 28:4765–4774. https://doi.org/10.1111/gcb.16220

Holleley CE, Geerts PG (2009) Multiplex Manager 1.0: a cross-platform computer program that plans and optimizes multiplex PCR. BioTechniques 46:511–517. https://doi.org/10.2144/000113156

Jacobs H-J (2000) Rote Liste der gefährdeten Grabwespen Mecklenburg-Vorpommerns (Hymenoptera Aculeata: Sphecidae). Das Umweltministerium des Landes Mecklenburg-Vorpommern

Johnstone RA, Cant MA, Field J (2012) Sex-biased dispersal, haplodiploidy and the evolution of helping in social insects. Proc R Soc B 279:787–793. https://doi.org/10.1098/rspb.2011.1257

Jombart T (2008) adegenet: a R package for the multivariate analysis of genetic markers. Bioinformatics 24:1403–1405. https://doi.org/10.1093/bioinformatics/btn129

Jombart T, Devillard S, Balloux F (2010) Discriminant analysis of principal components: a new method for the analysis of genetically structured populations. BMC Genet 11:94. https://doi.org/10.1186/1471-2156-11-94

Jost L (2008) G_ST_ and its relatives do not measure differentiation. Mol Ecol 17:4015–4026. https://doi.org/10.1111/j.1365-294X.2008.03887.x

Kamvar ZN, Larsen MM, Kanaskie AM, et al (2015) Spatial and Temporal Analysis of Populations of the Sudden Oak Death Pathogen in Oregon Forests. Phytopathology 105:982–989. https://doi.org/10.1094/PHYTO-12-14-0350-FI

Kamvar ZN, Tabima JF, Grünwald NJ (2014) PopprJ: an R package for genetic analysis of populations with clonal, partially clonal, and/or sexual reproduction. PeerJ 2:e281. https://doi.org/10.7717/peerj.281

Keenan K, McGinnity P, Cross TF, et al (2013) diveRsityJ: An R package for the estimation and exploration of population genetics parameters and their associated errors. Methods Ecol Evol 4:782–788. https://doi.org/10.1111/2041-210X.12067

Kim KS, Sappington TW (2013) Population genetics strategies to characterize long-distance dispersal of insects. J Asia Pac Entomol 16:87–97. https://doi.org/10.1016/j.aspen.2012.11.004

Kimoto C, DeBano SJ, Thorp RW, et al (2012) Short-term responses of native bees to livestock and implications for managing ecosystem services in grasslands. Ecosphere 3:88. https://doi.org/10.1890/ES12-00118.1

Klein WF, Lefeber V (2004) Crabronidae – graafwespen. In: De wespen en mieren van Nederland (ed. by T.M.J. Peeters). Naturalis en KNNV, Utrecht, The Netherlands, pp 356–430

Knight SM, Pitman GM, Flockhart DTT, Norris DR (2019) Radio-tracking reveals how wind and temperature influence the pace of daytime insect migration. Biol Lett 15:20190327. https://doi.org/10.1098/rsbl.2019.0327

Kun Á, Scheuring I (2006) The evolution of density-dependent dispersal in a noisy spatial population model. Oikos 115:308–320. https://doi.org/10.1111/j.2006.0030-1299.15061.x

Larsson FK (1986) Increased nest density of the digger wasp Bembix rostrata as a response to parasites and predators (Hymenoptera: Sphecidae). Entomol Gener 12:71–75

Larsson FK, Tengö J (1989) It is not always good to be large; some female fitness components in a temperate digger wasp, Bembix rostrata (Hymenoptera: Sphecidae). J Kans Entomol Soc 62:490–495

Leitch KJ, Ponce FV, Dickson WB, et al (2021) The long-distance flight behavior of Drosophila supports an agent-based model for wind-assisted dispersal in insects. PNAS 118:e2013342118. https://doi.org/10.1073/pnas.2013342118

Lohse K, Ross L (2015) What haplodiploids can teach us about hybridization and speciation. Mol Ecol 24:5075–5077. https://doi.org/10.1111/mec.13393

Lowe WH, Kovach RP, Allendorf FW (2017) Population Genetics and Demography Unite Ecology and Evolution. Trends Ecol Evol 32:141–152. https://doi.org/10.1016/j.tree.2016.12.002

Maes D, Bonte D (2006) Using distribution patterns of five threatened invertebrates in a highly fragmented dune landscape to develop a multispecies conservation approach. Biological Conservation 133:490–499. https://doi.org/10.1016/j.biocon.2006.08.001

Maes D, Van Dyck H (2022) Climate-driven range expansion through anthropogenic landscapes: Landscape connectivity matters. Glob Chang Biol 28:4920–4922. https://doi.org/10.1111/gcb.16180

Meirmans PG (2015) Seven common mistakes in population genetics and how to avoid them. Mol Ecol 24:3223–3231. https://doi.org/10.1111/mec.13243

Nei M (1972) Genetic Distance between Populations. Am Nat 106:283–292

Nielsen ET (1945) Moeurs des Bembex. Spoolia Zoologica Musei Hauniensis VII. Universitetets Zoologiske Museum, København, Denmark

Paetkau D, Slade R, Burden M, Estoup A (2004) Genetic assignment methods for the direct, real-time estimation of migration rate: a simulation-based exploration of accuracy and power. Mol Ecol 13:55–65. https://doi.org/10.1046/j.1365-294X.2004.02008.x

Paradis E (2010) pegas: an R package for population genetics with an integrated-modular approach. Bioinformatics 26:419–420. https://doi.org/10.1093/bioinformatics/btp696

Patten MM, Carioscia SA, Linnen CR (2015) Biased introgression of mitochondrial and nuclear genes: a comparison of diploid and haplodiploid systems. Mol Ecol 24:5200–5210. https://doi.org/10.1111/mec.13318

Piry S, Alapetite A, Cornuet J-M, et al (2004) GENECLASS2: A Software for Genetic Assignment and First-Generation Migrant Detection. J Hered 95:536–539. https://doi.org/10.1093/jhered/esh074

Potts SG, Imperatriz-Fonseca V, Ngo HT, et al (2016) Safeguarding pollinators and their values to human well-being. Nature 540:220–229. https://doi.org/10.1038/nature20588

Provoost S, Ampe C, Bonte D, et al (2004) Ecology, management and monitoring of grey dunes in Flanders. J Coast Conserv 10:33–42. https://doi.org/10.1652/1400-0350(2004)010[0033:EMAMOG]2.0.CO;2

Provoost S, Jones MLM, Edmondson SE (2011) Changes in landscape and vegetation of coastal dunes in northwest Europe: a review. J Coast Conserv 15:207–226. https://doi.org/10.1007/s11852-009-0068-5

QGIS Development Team (2020) QGIS Geographic Information System

Rannala B, Mountain JL (1997) Detecting immigration by using multilocus genotypes. PNAS 94:9197– 9201. https://doi.org/10.1073/pnas.94.17.9197

Sanllorente O, Ruano F, Tinaut A (2015) Large-scale population genetics of the mountain ant Proformica longiseta (Hymenoptera: Formicidae). Popul Ecol 57:637–648. https://doi.org/10.1007/s10144-015-0505-2

Schneiders A, Alaerts K, Michels H, et al (2020) Natuurrapport 2020: feiten en cijfers voor een nieuw biodiversiteitsbeleid. Mededelingen van het Instituut voor Natuur- en Bosonderzoek, Nr. 2. Instituut voor Natuur-en Bosonderzoek

Schöne H, Tengö J (1981) Competition of males, courtship behaviour and chemical communication in the digger wasp Bembix rostrata (Hymenoptera, Sphecidae). Behaviour 77:44–66

Takezaki N, Nei M (1996) Genetic Distances and Reconstruction of Phylogenetic Trees From Microsatellite DNA. Genetics 144:389–399. https://doi.org/10.1093/genetics/144.1.389

Van Dyck H, Baguette M (2005) Dispersal behaviour in fragmented landscapes: Routine or special movements? Basic Appl Ecol 6:535–545. https://doi.org/10.1016/j.baae.2005.03.005

van Klink R, van der Plas F, van Noordwijk CGE (Toos), et al (2015) Effects of large herbivores on grassland arthropod diversity. Biol Rev 90:347–366. https://doi.org/10.1111/brv.12113

van Strien MJ, Holderegger R, Van Heck HJ (2015) Isolation-by-distance in landscapes: considerations for landscape genetics. Heredity 114:27–37. https://doi.org/10.1038/hdy.2014.62

Wagner DL, Grames EM, Forister ML, et al (2021) Insect decline in the Anthropocene: Death by a thousand cuts. PNAS 118:e2023989118. https://doi.org/10.1073/pnas.2023989118

Waples RS (2015) Testing for Hardy–Weinberg Proportions: Have We Lost the Plot? J Hered 106:1– 19. https://doi.org/10.1093/jhered/esu062

Weir BS, Cockerham CC (1984) Estimating F-Statistics for the Analysis of Population Structure. Evolution 38:1358–1370

Welti EAR, Joern A, Ellison AM, et al (2021) Studies of insect temporal trends must account for the complex sampling histories inherent to many long-term monitoring efforts. Nat Ecol Evol 5:589–591. https://doi.org/10.1038/s41559-021-01424-0

Wheeler B, Torchiano M (2016) lmPerm: Permutation Tests for Linear Models.

Wiklund C, Fagerström T (1977) Why do males emerge before females? A hypothesis to explain the incidence of protandry in butterflies. Oecologia 31:153–158. https://doi.org/10.1007/BF00346917

Zayed A (2004) Effective population size in Hymenoptera with complementary sex determination. Heredity 93:627–630. https://doi.org/10.1038/sj.hdy.6800588

Zayed A (2009) Bee genetics and conservation. Apidologie 40:237–262. https://doi.org/10.1051/apido/2009026

