## Supplementary materials S5 - output visualisation-HW-LD-NA for "Asymmetrical gene flow between coastal and inland dunes in a threatened digger wasp"

### Visualisation HW LD NA morethan10samplespops

Femke Batsleer

17-2-2022

```
library("dplyr")
library("tidyr")
library("tidyverse")
library("purrr")
library("adegenet")#v2.0.0 install.packages("adegenet", version = "2.0.0")
library("poppr")
library("ggplot2")
#library("genepop")
library("graph4lg")
library("pegas")
library("hierfstat")
library("PopGenReport")#v2.2.1 has to be installed
library("stringr")
library("devtools")
library("pkgload")
```

#### For populations that had at least 10 samples

General, the philosophy of Waples (2015) Journal of Heredity: Testing for Hardy-Weinberg proportions: have we lost the plot?

#### Null alleles

Was calculated (in other script 'Calculations HW LD NA.R') with null.all function from PopGenReport

First, looking at how many times freq of null alleles = 0 falls outside of confidence interval. Tables give number that they did not deviate (so the lower, the more it deviates). Second, looking at how many times abs(freq) > 0.2

First eight: 20 16 57 35 227 138 419 266 20 16 57 266 299 138 35 337 20 16 57 35 227 138 419 266 20 16 57 266 299 35 138 337

Not good: 20, 16, 57, 35, 138 (266)

35 57 419 307 138 35 57 419 307 138 266 35 57 419 307 138 35 57 419 307 138 266

Not good: 35 57 419 307 138 (266)

**conclusion: leave out 20 16 57 35 138 266 (419 307)**

```
#load data output from null allele analysis
NA_pop <- read.csv("Outputs/Output Null Alleles population morethan10samp level.csv",
                   sep=",", stringsAsFactors=FALSE)#populations (years combined)
NA_popyear <- read.csv("Outputs/Output Null Alleles popyear morethan10samp level.csv",
                       sep=",", stringsAsFactors=FALSE)#popyear (years separate)
```

```r
###function to make interpretable tables for null allele output####
null.all_tables <- function(output_test, group_var, group_name, observed=FALSE){
  group_var <- enquo(group_var)
  #add a sign symbol to indicate deviations and calcualte absolute value of observed
  null.all_df <- output_test %>% drop_na() %>%
    mutate(sign_dev = sign(sign(percentile2.5th) + sign(percentile97.5th))) %>%
    mutate(absobserved = abs(observed))

  if(observed==FALSE){#do test with significance levels for null alleles
    #calculate number of tests per locus/pop (some are NA, as alleles are sometimes fixed
    #in pops)
    testspergroup <- null.all_df %>% group_by(!!group_var) %>% summarise(n_tests=n())
    #calculate frequency of deviations and non-deviations
    tests_alldevs <- null.all_df %>% group_by(!!group_var, sign_dev) %>%
      summarise(freq=n())
    #ratio of non-deviations per group
    Nulltest_ratio <- tests_alldevs %>% filter(sign_dev==0) %>%
      left_join(testspergroup, by=group_name) %>%
      mutate(ratio_nodev = freq/n_tests)
  }
  else{#do test with observed; count the observed ones which have 0.2<abs(observed)
    testspergroup <- null.all_df %>% filter(absobserved<0.2) %>%
      group_by(!!group_var) %>% summarise(n_obs01 = n())
    tests_alldevs <- null.all_df %>% group_by(!!group_var) %>% summarise(n_tests=n())
    Nulltest_ratio <- tests_alldevs %>% left_join(testspergroup, by=group_name) %>%
      mutate(ratio_nodev = n_obs01/n_tests)
  }

  return(Nulltest_ratio)
}

###########################################################
###Looking at values where freq=0 of null alleles falls outside of confidence interval####
###########################################################
#Population level
Nulltest_pop.pop <- null.all_tables(NA_pop, group_var=pop, group_name="pop") %>%
    arrange(ratio_nodev) %>%
    rmarkdown::paged_table()
```

```
## `summarise()` has grouped output by 'pop'. You can override using the `.groups`
## argument.
```

```r
Nulltest_popyear.pop <- null.all_tables(NA_popyear, group_var=pop, group_name="pop") %>%
    arrange(ratio_nodev) %>%
    rmarkdown::paged_table()
```

```
## `summarise()` has grouped output by 'pop'. You can override using the `.groups`
## argument.
```

```r
Nulltest_pop.locus <- null.all_tables(NA_pop, group_var=locus, group_name="locus") %>%
    arrange(ratio_nodev) %>%
    rmarkdown::paged_table()
```

```
## `summarise()` has grouped output by 'locus'. You can override using the
## `.groups` argument.
```

```r
Nulltest_pop.locus
```

```r
Nulltest_popyear.locus <- null.all_tables(NA_popyear, group_var=locus,
                                          group_name="locus") %>%
  arrange(ratio_nodev) %>%
  rmarkdown::paged_table()
```

```
## `summarise()` has grouped output by 'locus'. You can override using the
## `.groups` argument.
```

```r
Nulltest_popyear.locus
```

```r
#leave out populations with lowest ratio_nodev (<0.6), so populations that already
#act strange (not in HW-equilibrium probably) are left out
weird_pops <- Nulltest_pop.pop %>% filter(ratio_nodev < 0.6)%>%
    select(pop) %>% as.vector()
(Nulltest_pop_sel.locus <- null.all_tables(filter(NA_pop, !pop %in% weird_pops$pop),
                                           group_var=locus, group_name="locus") %>%
    arrange(ratio_nodev) %>%
    rmarkdown::paged_table())
```

```
## `summarise()` has grouped output by 'locus'. You can override using the
## `.groups` argument.
```

```r
weird_popyear <- Nulltest_popyear.pop %>% filter(ratio_nodev < 0.6)%>%
    select(pop) %>% as.vector()
(Nulltest_popyear_sel.locus <- null.all_tables(filter(NA_popyear,
                                                      !pop %in% weird_popyear$pop),
                                               group_var=locus, group_name="locus") %>%
    arrange(ratio_nodev) %>%
    rmarkdown::paged_table())
```

```
## `summarise()` has grouped output by 'locus'. You can override using the
## `.groups` argument.
```

```r
##########################################################
###Looking at how many times freq null alleles > 0.2 ####
###########################################################
Nulltest_pop.pop <- null.all_tables(NA_pop, group_var=pop, group_name="pop",
                                    observed=TRUE) %>%
    arrange(ratio_nodev) %>%
    rmarkdown::paged_table()
Nulltest_popyear.pop <- null.all_tables(NA_popyear, group_var=pop, group_name="pop",
                                        observed=TRUE) %>%
    arrange(ratio_nodev) %>%
    rmarkdown::paged_table()

(Nulltest_pop.locus <- null.all_tables(NA_pop, group_var=locus, group_name="locus",
                                       observed=TRUE) %>%
    arrange(ratio_nodev) %>%
    rmarkdown::paged_table())
```

```r
(Nulltest_popyear.locus <- null.all_tables(NA_popyear, group_var=locus, group_name="locus",
                                           observed=TRUE) %>%
    arrange(ratio_nodev) %>%
    rmarkdown::paged_table())
```

```
#leave out populations with lowest ratio_nodev (<0.6)
weird_pops <- Nulltest_pop.pop %>% filter(ratio_nodev < 0.6)%>%
    select(pop) %>% as.vector()
(Nulltest_pop_sel.locus <- null.all_tables(filter(NA_pop, !pop %in% weird_pops$pop),
                                           group_var=locus, group_name="locus",
                                           observed=TRUE) %>%
    arrange(ratio_nodev) %>%
    rmarkdown::paged_table())
```

```
weird_popyear <- Nulltest_popyear.pop %>% filter(ratio_nodev < 0.6)%>%
    select(pop) %>% as.vector()
(Nulltest_popyear_sel.locus <- null.all_tables(filter(NA_popyear, !pop %in%
                                                        weird_popyear$pop),
                                           group_var=locus, group_name="locus",
                                           observed=TRUE) %>%
    arrange(ratio_nodev) %>%
    rmarkdown::paged_table())
```

##### Linkage disequilibrium

Was calculated with poppr (in other script 'Calculations HW LD NA.R')

In general, not very high deviations. *To leave out: 111* (403-110, 12-138, 337-153)

```
##Population level
LD_pop <- read.csv("Outputs/Output LD population morethan10samp level.csv", sep=",")

##pop##
LD.all_pop_df_perpop <- LD_pop %>% separate(pairloci, into=c("Locus1", "Locus2")) %>%
  filter(p.Ia<0.05) %>% group_by(pop) %>%
  summarise(n_sign=n()) %>% arrange(desc(n_sign)) #Leave out Wetteren?
LD.all_pop_df_perpop
```

```
## # A tibble: 39 x 2
##    pop               n_sign
##    <chr>              <int>
##  1 Wetteren              68
##  2 Kalmthout1            38
##  3 Keiheuvel             34
##  4 Lagland               31
##  5 Oosthoekduinen        28
##  6 Simliduinen           26
##  7 Kopberg               25
##  8 Kortenhoeff-NL        25
##  9 DuinbossenDeHaan      24
## 10 Bredene               23
## # ... with 29 more rows
```

```
LD.all_pop_df <- LD_pop %>% separate(pairloci, into=c("Locus1", "Locus2")) %>%
  filter(p.Ia<0.05) %>% group_by(Locus1, Locus2) %>%
  summarise(n_sign=n())
```

```
## `summarise()` has grouped output by 'Locus1'. You can override using the
## `.groups` argument.
```

```r
LD.all_pop_df_nowett <- LD_pop %>% separate(pairloci, into=c("Locus1", "Locus2")) %>%
  filter(p.Ia<0.05) %>%
  filter(pop != "Wetteren") %>%
  group_by(Locus1, Locus2) %>%
  summarise(n_sign=n())
```

```
## `summarise()` has grouped output by 'Locus1'. You can override using the
## `.groups` argument.
```

```r
(LD.all_pop_df_problems_nowett <- LD.all_pop_df_nowett %>% filter(n_sign>4) %>%
    #outside of CI for probability of having >2 times a significant test for 38/39
    #populations
  rmarkdown::paged_table())
```

```r
(LD.all_pop_nowett_plot <- ggplot(LD.all_pop_df_problems_nowett, aes(x=Locus1, y=Locus2,
                                                                    fill=n_sign)) +
  geom_tile() +
  scale_fill_distiller(palette = "YlOrRd", trans="reverse") +
  theme(axis.text.x = element_text(angle = 90, vjust = 0.5, hjust=1)))
```

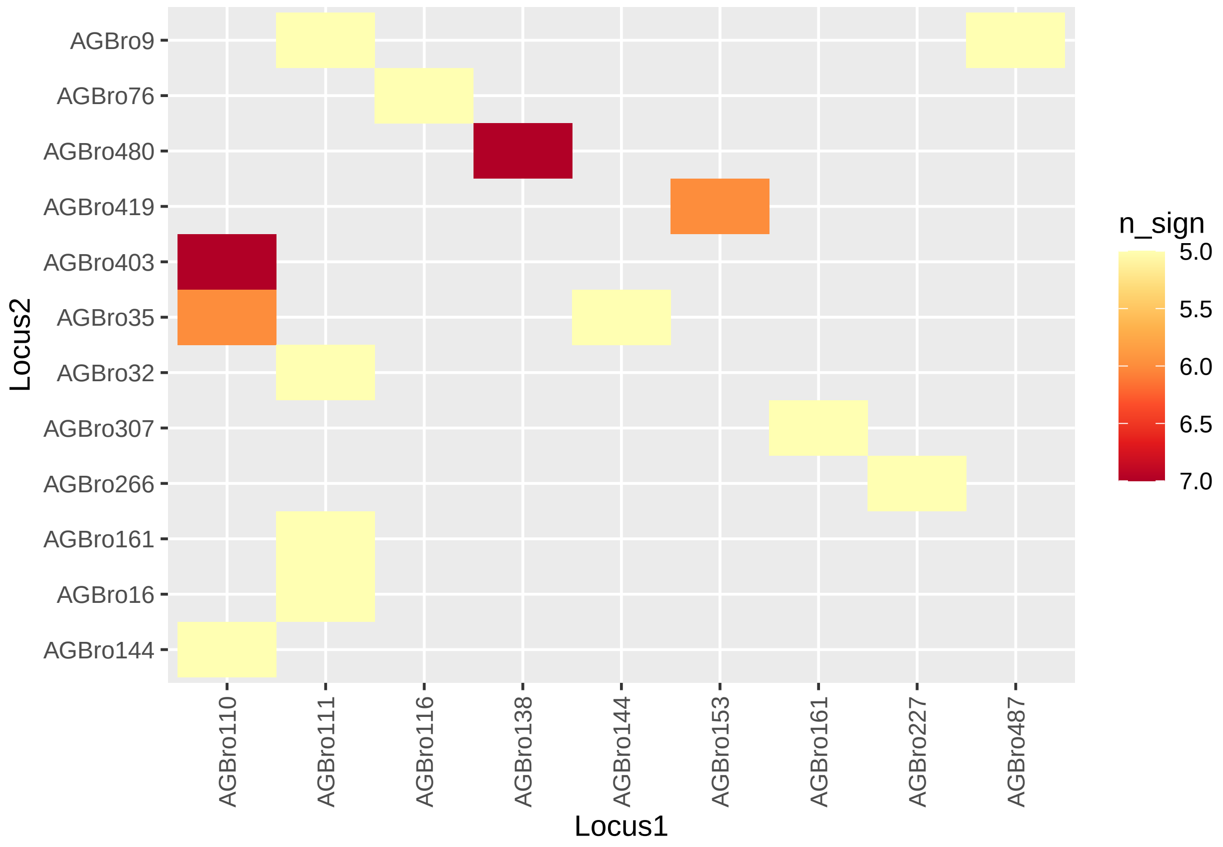


```r
(LD.all_pop_df_problems <- LD.all_pop_df %>% filter(n_sign>4) %>%
    #outside of CI for probability of having >2 times a significant test for
    #38 populations
  rmarkdown::paged_table())
```

```r
(LD.all_pop_plot <- ggplot(LD.all_pop_df_problems, aes(x=Locus1, y=Locus2, fill=n_sign)) +
  geom_tile() +
```

```
  scale_fill_distiller(palette = "YlOrRd", trans="reverse") +
  theme(axis.text.x = element_text(angle = 90, vjust = 0.5, hjust=1)))
```

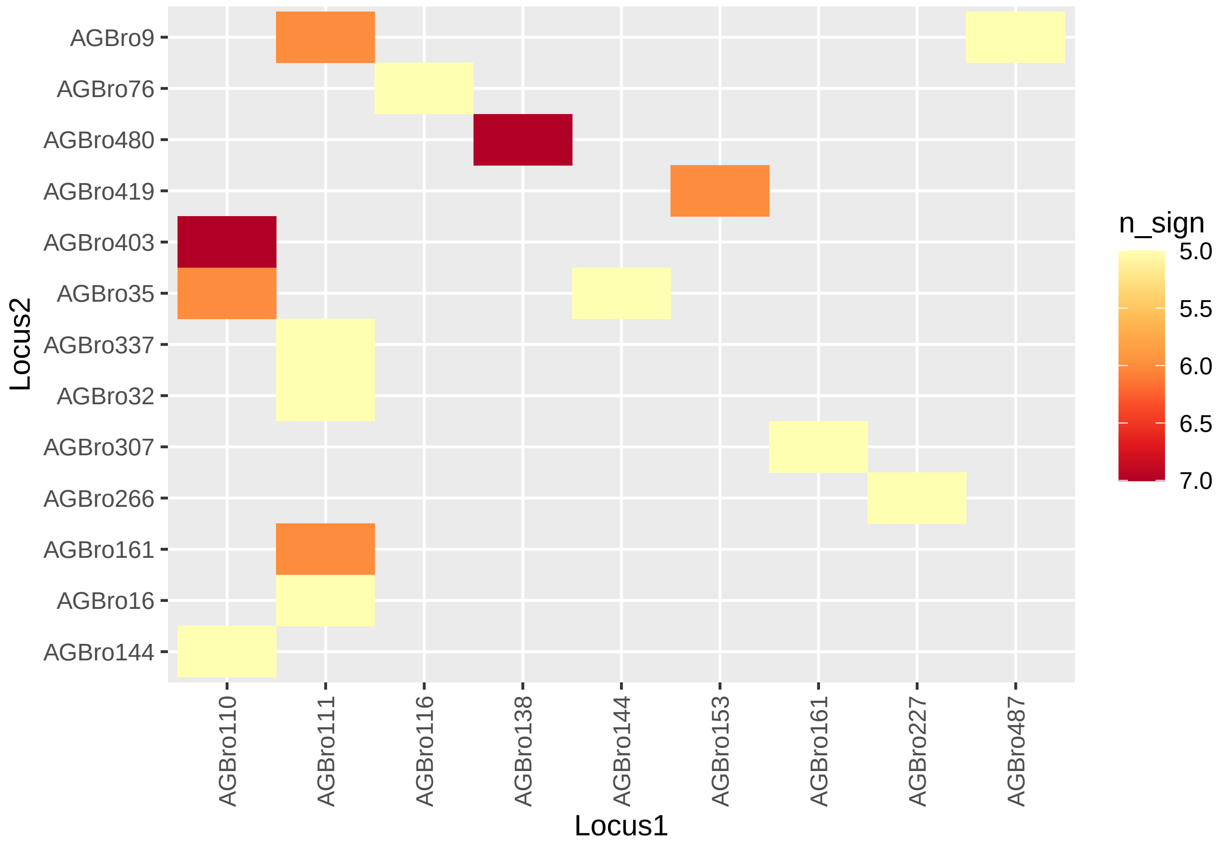


```
#Population-year level

LD_popyear <- read.csv("Outputs/Output LD popyear morethan10samp level.csv", sep=",")

##pop##
(LD.all_popyear_df_perpop <- LD_popyear %>% separate(pairloci, into=c("Locus1", "Locus2")) %>%
  filter(p.Ia<0.05) %>% group_by(pop) %>%
  summarise(n_sign=n()) %>% arrange(desc(n_sign))) #Leave out Wetteren?
```

```
## # A tibble: 41 x 2
##    pop                          n_sign
##    <chr>                         <int>
##  1 Keiheuvel2020                    41
##  2 Kalmthout12020                   35
##  3 Wetteren2020                     32
##  4 Kopberg2020                      29
##  5 Oosthoekduinen2018               27
##  6 Kortenhoeff-NL2020               26
##  7 DuinbossenDeHaan2020             25
##  8 Geel-Bel2018                     22
##  9 Vloethemveld-Zuid2020            22
## 10 Westhoek vissersdorp2018         21
## # ... with 31 more rows
```

```
LD.all_popyear_df <- LD_popyear %>% separate(pairloci,
                                             into=c("Locus1", "Locus2")) %>%
  filter(p.Ia<0.05) %>% group_by(Locus1, Locus2) %>%
  summarise(n_sign=n())
```

```
## `summarise()` has grouped output by 'Locus1'. You can override using the
## `.groups` argument.
```

```
(LD.all_popyear_df_problems <- LD.all_popyear_df %>% filter(n_sign>4) %>%
    #outside of CI for probability of having >2 times a significant test for 40/41 populations
  rmarkdown::paged_table())
```

```
(LD.all_popyear_plot <- ggplot(LD.all_popyear_df_problems, aes(x=Locus1,
                                                                y=Locus2, fill=n_sign)) +
  geom_tile() +
  scale_fill_distiller(palette = "YlOrRd", trans="reverse") +
  theme(axis.text.x = element_text(angle = 90, vjust = 0.5, hjust=1)))
```

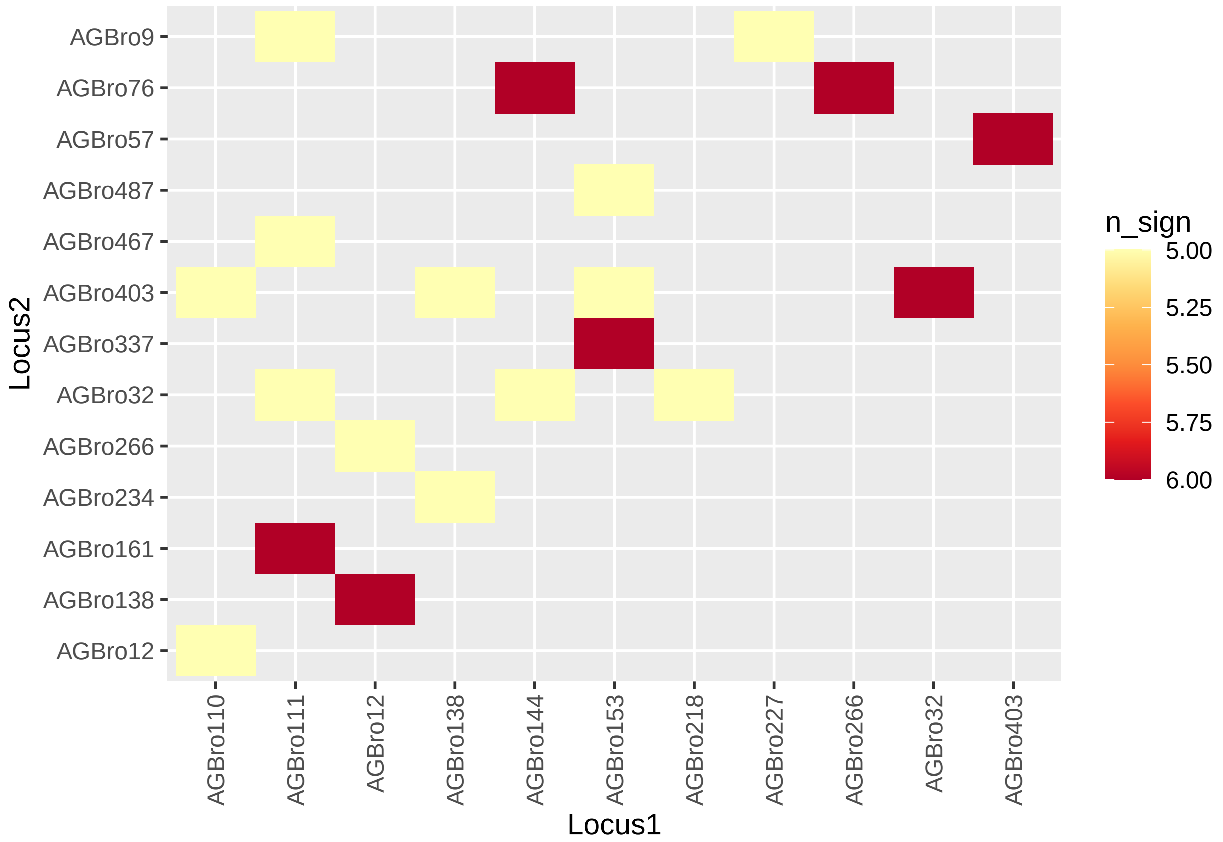


##### Hardy-Weinberg

Calculated with pegas (in other script 'Calculations HW LD NA.R')

*Large deviations: 111, 35, 419, 57,* (9) small deviations: 12, 138, 144, 307, 375, 403, 487

```
HW_pop <- read.csv("Outputs/Output HW population morethan10samp level.csv", sep=",")
HW_popyear <- read.csv("Outputs/Output HW popyear morethan10samp level.csv", sep=",")

#Add probability intervals to the plots
```

```r
#get two-sides prob-interval of 0.025<P<97.5 out of binomial distribution
#with excel: =BINOM.DIST(1(-...);18; 0.05;TRUE): 17-35 significant tests are expected


##pop##
#histogram per locus of tests
(hist_locus_HW_pop <- ggplot(HW_pop, aes(x=Pr.exact))+
  geom_histogram(bins=10)+
  geom_hline(yintercept = 4, linetype="dashed")+ #for 39 pops or tests per locus (#number)
  facet_wrap(~ locus))
```

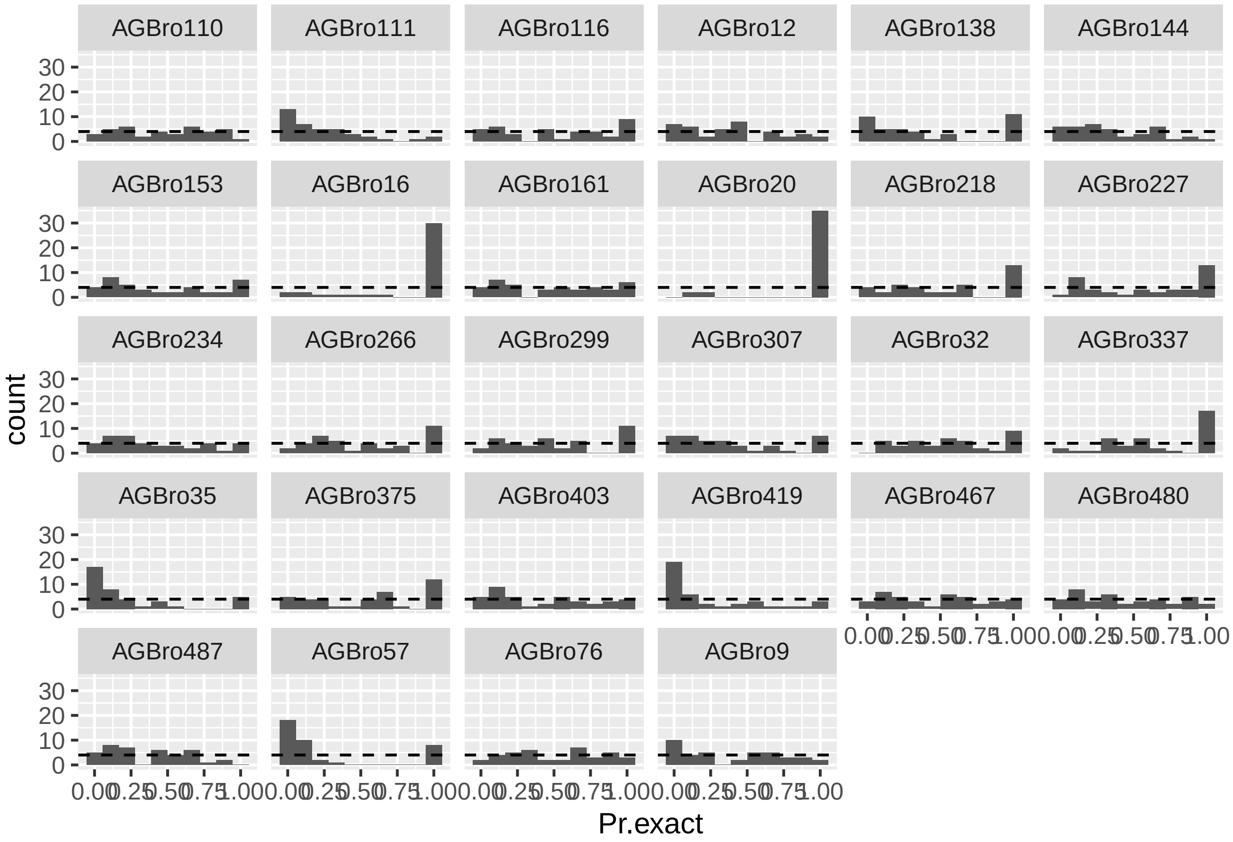


```r
#histogram per pop of tests
(hist_pop_HW_pop <- ggplot(HW_pop, aes(x=Pr.exact))+
  geom_histogram(bins=10)+
  geom_hline(yintercept = 3, linetype="dashed")+ #for 28 loci or test per pop
  facet_wrap(~ pop))
```

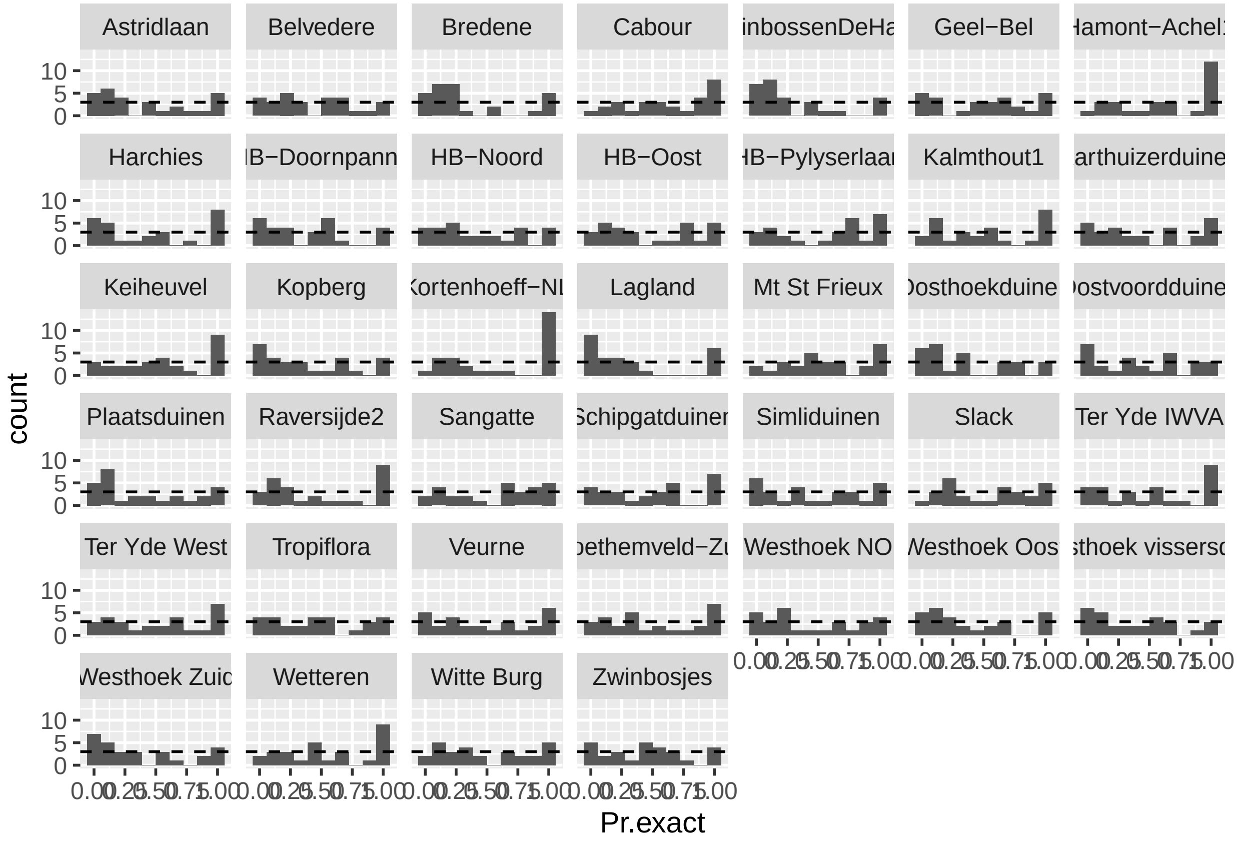


```
##popyear##
#histogram per locus of tests
(hist_locus_HW_popyear <- ggplot(HW_popyear, aes(x=Pr.exact))+
  geom_histogram(bins=10)+
  geom_hline(yintercept = 4, linetype="dashed")+ #for 41 pops or tests per locus (#number)
  facet_wrap(~ locus))
```

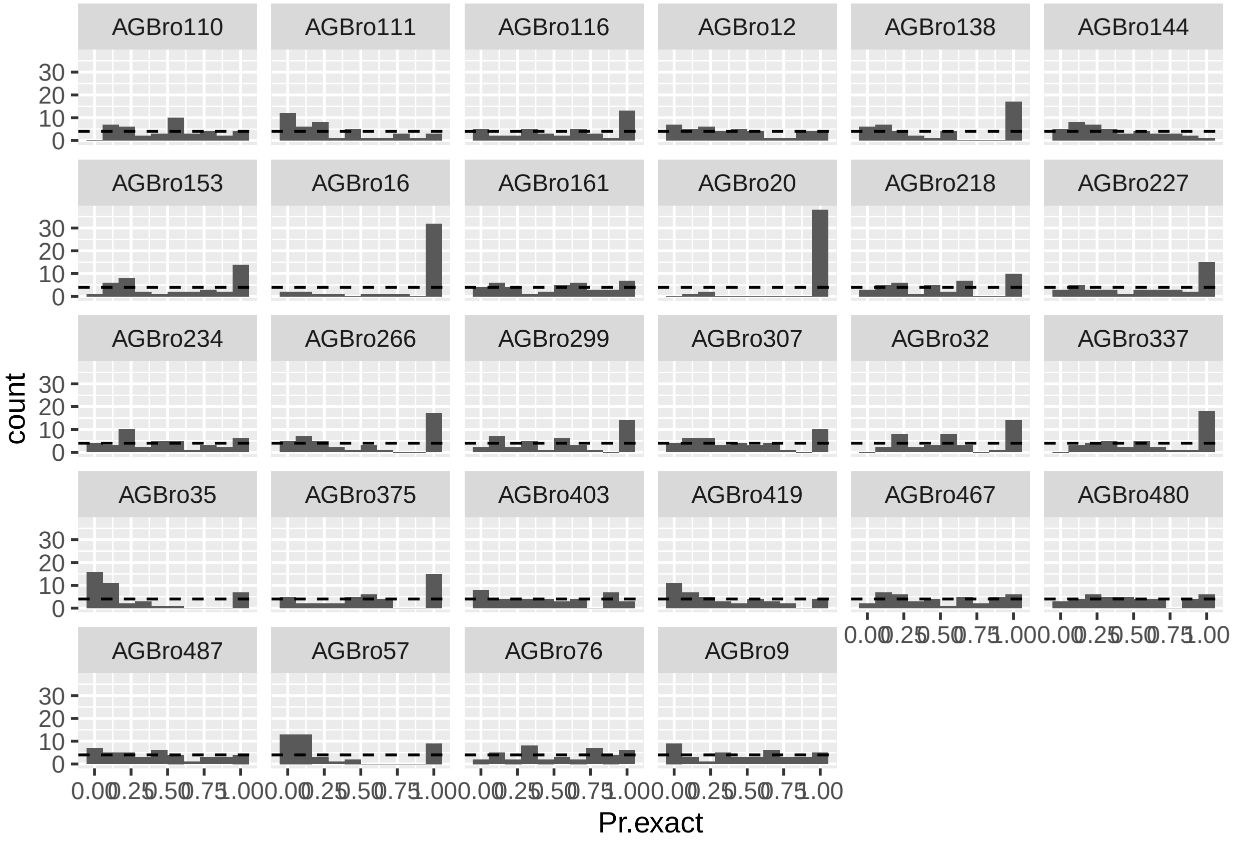


```r
#histogram per pop of tests
(hist_pop_HW_popyear <- ggplot(HW_popyear, aes(x=Pr.exact))+
  geom_histogram(bins=10)+
  geom_hline(yintercept = 3, linetype="dashed")+ #for 28 loci or test per pop
  facet_wrap(~ pop))
```

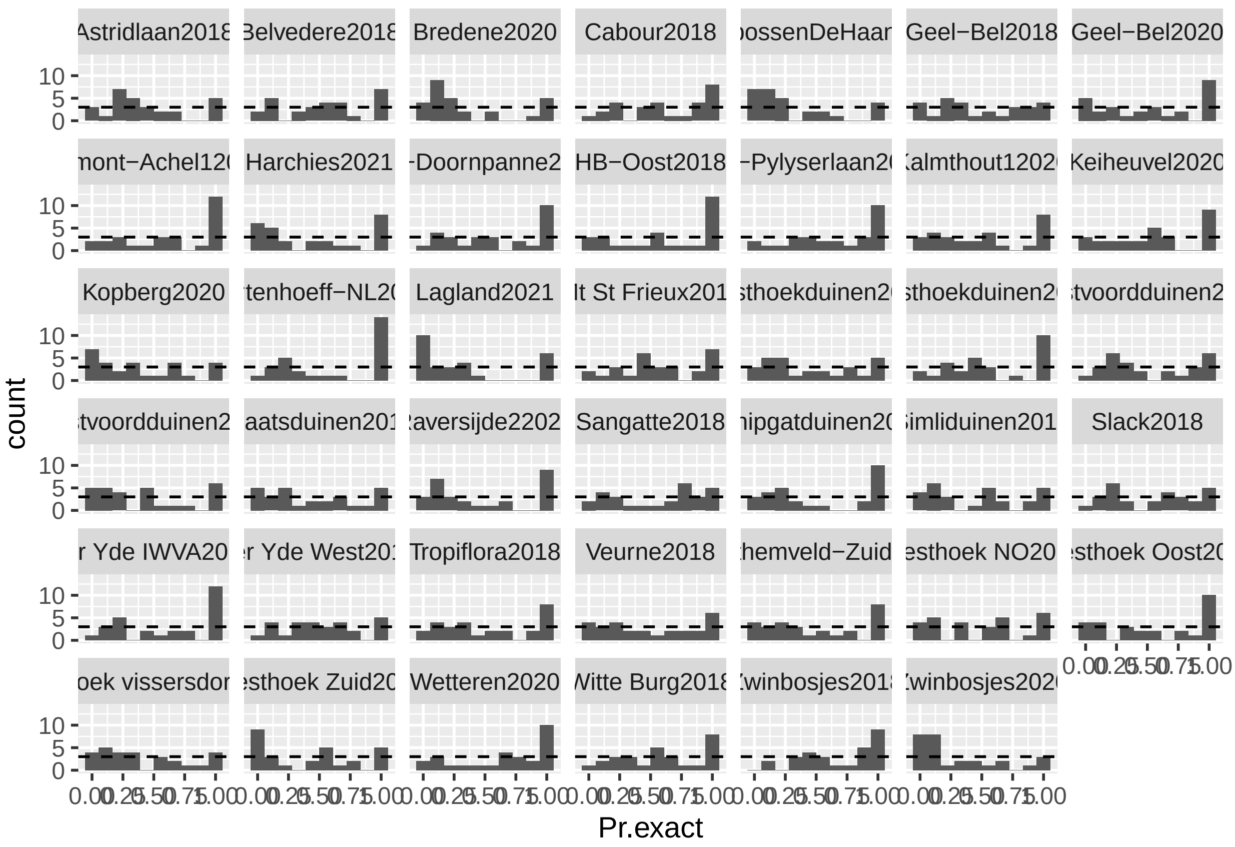


#### Conclusion

I will leave out: **35 57 419 (HW, NA) 111 (HW LD) 20 16 (NA) 138 (HW NA LD)**, these had large deviations and/or were recurring in the different tests.
