## Supplementary materials S6-S9 for "Asymmetrical gene flow between coastal and inland dunes in a threatened digger wasp"

### S6. AMOVA tables

The results of the hierarchical Analysis of Molecular Variance (AMOVA) analysis for populations which were sampled across two years are given in table S6.1. Analyses were performed with the package poppr. Only populations that were sampled across two years were considered for this analysis.

Table S6.1: results of the AMOVA analysis for population samples across the two sampling years. Df: degrees of freedom, sigma: variance.

| **Source of variation** | **Df** | **Sum of Squares** | **Sigma** | **% of variation** |
| --- | --- | --- | --- | --- |
| Between populations | 22 | 665.30 | 0.375 | 3.106 |
| Between year within population | 23 | 302.62 | 0.023 | 0.193 |
| Between samples within year | 476 | 6041.40 | 1.022 | 8.466 |
| Within samples | 522 | 5558.52 | 10.649 | 88.235 |

The results of the AMOVA analyses to check variation between and within populations, for both coastal and inland region separately, are in given in tables S6.2-5.3. All samples were taken into account for this analysis.

Table S6.2: results of the AMOVA analysis for population samples at the coast. Df: degrees of freedom, sigma: variance.

| **Source of variation in coastal region** | **Df** | **Sum of Squares** | **Sigma** | **% of variation** |
| --- | --- | --- | --- | --- |
| Between populations | 39 | 790.57 | 0.208 | 1.706 |
| Between samples within population | 631 | 8414.31 | 1.355 | 11.114 |
| Within samples | 671 | 7129.82 | 10.626 | 87.180 |

Table S6.3: results of the AMOVA analysis for population samples inland. Df: degrees of freedom, sigma: variance.

| **Source of variation in inland region** | **Df** | **Sum of Squares** | **Sigma** | **% of variation** |
| --- | --- | --- | --- | --- |
| Between populations | 12 | 571.75 | 1.324 | 14.826 |
| Between samples within population | 183 | 1565.75 | 0.949 | 10.625 |
| Within samples | 196 | 1305.00 | 6.658 | 74.549 |

### S7. Population level statistics

Population level statistics for all sampled population are given in table S7.1. Figures S7.1-S7.3 give a visualization of these data for all populations.

Table S7.1: population level statistics for all sampled populations (table S1.1). Following abbreviations are used: Region, population number (ID; see map main manuscript), given population name (Population name), number of samples (n), number of private alleles (NP), rarefied allelic richness (AR), expected heterozygosity (H_e_), observed heterozygosity (H_o_) and inbreeding coefficient (F_IS_). Standard errors (SE) are given for AR, H_e_, H_o_ and F_IS_.

| **Region** | **ID** | **Population**  **name** | **n** | **NP** | **AR ± SE** | **H_e_ ± SE** | **H_o_ ± SE** | **F_IS_ ± SE** |
| --- | --- | --- | --- | --- | --- | --- | --- | --- |
| France-Picardie | 1 | Mt St Frieux | 19 | 1 | 2.523 ± 0.153 | 0.547 ± 0.037 | 0.559 ± 0.041 | -0.022 ± 0.026 |
|  | 2 | Slack | 20 | 1 | 2.605 ± 0.141 | 0.587 ± 0.031 | 0.585 ± 0.038 | 0.003 ± 0.035 |
|  | 3 | Sangatte | 20 | 0 | 2.630 ± 0.160 | 0.588 ± 0.029 | 0.555 ± 0.033 | 0.057 ± 0.027 |
| Flanders-coast | 4 | Westhoek Zuid | 21 | 0 | 2.623 ± 0.148 | 0.584 ± 0.032 | 0.520 ± 0.035 | 0.105 ± 0.042 |
|  | 5 | Cabour | 10 | 0 | 2.769 ± 0.160 | 0.63 ± 0.028 | 0.619 ± 0.033 | 0.004 ± 0.046 |
|  | 6 | Veurne | 20 | 0 | 2.570 ± 0.145 | 0.586 ± 0.03 | 0.498 ± 0.032 | 0.149 ± 0.028 |
|  | 7 | Tropiflora | 29 | 1 | 2.628 ± 0.157 | 0.575 ± 0.036 | 0.535 ± 0.038 | 0.072 ± 0.032 |
|  | 8 | Westhoek vissersdorp | 23 | 0 | 2.511 ± 0.138 | 0.563 ± 0.03 | 0.494 ± 0.036 | 0.126 ± 0.036 |
|  | 9 | Westhoek NO | 30 | 1 | 2.657 ± 0.162 | 0.594 ± 0.032 | 0.547 ± 0.034 | 0.083 ± 0.021 |
|  | 10 | Westhoek Oost | 18 | 0 | 2.647 ± 0.162 | 0.587 ± 0.034 | 0.558 ± 0.048 | 0.054 ± 0.057 |
|  | 11 | Oosthoekduinen | 29 | 0 | 2.711 ± 0.154 | 0.613 ± 0.028 | 0.584 ± 0.034 | 0.047 ± 0.040 |
|  | 12 | Belvedere | 18 | 0 | 2.670 ± 0.147 | 0.604 ± 0.029 | 0.515 ± 0.038 | 0.151 ± 0.042 |
|  | 13 | HB-Noord | 16 | 0 | 2.686 ± 0.164 | 0.594 ± 0.039 | 0.545 ± 0.051 | 0.110 ± 0.054 |
|  | 14 | HB-Doornpanne | 17 | 0 | 2.659 ± 0.154 | 0.589 ± 0.036 | 0.501 ± 0.033 | 0.137 ± 0.035 |
|  | 15 | HB-Oost | 16 | 1 | 2.725 ± 0.156 | 0.617 ± 0.028 | 0.576 ± 0.030 | 0.057 ± 0.041 |
|  | 16 | HB-Pylyserlaan | 18 | 0 | 2.661 ± 0.16 | 0.595 ± 0.033 | 0.551 ± 0.045 | 0.087 ± 0.049 |
|  | 17 | Schipgatduinen | 16 | 0 | 2.478 ± 0.16 | 0.539 ± 0.037 | 0.490 ± 0.030 | 0.063 ± 0.040 |
|  | 18 | Witte Burg | 19 | 0 | 2.629 ± 0.142 | 0.599 ± 0.028 | 0.577 ± 0.024 | 0.023 ± 0.036 |
|  | 19 | Astridlaan | 24 | 0 | 2.688 ± 0.153 | 0.603 ± 0.030 | 0.555 ± 0.034 | 0.071 ± 0.046 |
|  | 20 | Plaatsduinen | 27 | 0 | 2.630 ± 0.147 | 0.590 ± 0.030 | 0.513 ± 0.036 | 0.140 ± 0.035 |
|  | 21 | Oostvoordduinen | 30 | 0 | 2.634 ± 0.136 | 0.594 ± 0.027 | 0.580 ± 0.033 | 0.021 ± 0.039 |
|  | 22 | Ter Yde West | 13 | 0 | 2.614 ± 0.147 | 0.595 ± 0.029 | 0.582 ± 0.036 | 0.005 ± 0.058 |
|  | 23 | Ter Yde IWVA | 19 | 0 | 2.581 ± 0.13 | 0.582 ± 0.027 | 0.576 ± 0.027 | -0.009 ± 0.045 |
|  | 24 | Karthuizerduinen | 18 | 0 | 2.614 ± 0.147 | 0.585 ± 0.031 | 0.503 ± 0.036 | 0.136 ± 0.052 |
|  | 25 | Simliduinen | 27 | 0 | 2.621 ± 0.143 | 0.590 ± 0.030 | 0.560 ± 0.024 | 0.032 ± 0.030 |
|  | 26 | Sint-Laureins1 | 9 | 0 | 2.655 ± 0.163 | 0.602 ± 0.033 | 0.502 ± 0.036 | 0.141 ± 0.065 |
|  | 27 | Sint-Laureins2 | 7 | 0 | 2.842 ± 0.176 | 0.64 ± 0.036 | 0.528 ± 0.049 | 0.173 ± 0.060 |
|  | 28 | Warandeduinen Middelkerke | 7 | 0 | 2.718 ± 0.148 | 0.612 ± 0.034 | 0.517 ± 0.042 | 0.125 ± 0.070 |
|  | 29 | Raversijde1 | 8 | 0 | 2.634 ± 0.162 | 0.597 ± 0.042 | 0.369 ± 0.045 | 0.361 ± 0.069 |
|  | 30 | Raversijde2 | 10 | 0 | 2.576 ± 0.136 | 0.597 ± 0.03 | 0.470 ± 0.038 | 0.211 ± 0.054 |
|  | 31 | Raversijde3 | 5 | 0 | 2.442 ± 0.18 | 0.529 ± 0.058 | 0.395 ± 0.05 | 0.201 ± 0.072 |
|  | 32 | Raversijde4 | 5 | 0 | 2.724 ± 0.188 | 0.616 ± 0.036 | 0.604 ± 0.059 | 0.018 ± 0.084 |
|  | 33 | Fort Napoleon | 9 | 0 | 2.502 ± 0.139 | 0.556 ± 0.036 | 0.517 ± 0.041 | 0.053 ± 0.064 |
|  | 34 | Spanjaardduinen Oostende | 8 | 0 | 2.696 ± 0.161 | 0.617 ± 0.035 | 0.451 ± 0.042 | 0.264 ± 0.055 |
|  | 35 | Bredene | 10 | 1 | 2.513 ± 0.157 | 0.576 ± 0.034 | 0.364 ± 0.044 | 0.359 ± 0.069 |
|  | 36 | Duinbossen DeHaan | 12 | 0 | 2.575 ± 0.134 | 0.603 ± 0.027 | 0.403 ± 0.034 | 0.333 ± 0.050 |
|  | 37 | Zwinbosjes-grazed | 9 | 0 | 2.448 ± 0.151 | 0.544 ± 0.045 | 0.481 ± 0.048 | 0.098 ± 0.054 |
|  | 38 | Zwinbosjes | 35 | 0 | 2.551 ± 0.152 | 0.567 ± 0.035 | 0.504 ± 0.034 | 0.108 ± 0.022 |
|  | 39 | Vloethemveld-Noord | 9 | 0 | 2.412 ± 0.13 | 0.528 ± 0.040 | 0.458 ± 0.037 | 0.102 ± 0.047 |
|  | 40 | Vloethemveld-Zuid | 11 | 0 | 2.535 ± 0.153 | 0.555 ± 0.042 | 0.483 ± 0.042 | 0.123 ± 0.038 |
| Flanders-inland | 41 | Wetteren | 28 | 0 | 2.127 ± 0.129 | 0.443 ± 0.046 | 0.446 ± 0.049 | 0.005 ± 0.030 |
|  | 42 | Kortenhoeff-NL | 10 | 0 | 2.178 ± 0.14 | 0.446 ± 0.046 | 0.392 ± 0.044 | 0.091 ± 0.048 |
|  | 43 | Kalmthout | 12 | 0 | 2.387 ± 0.151 | 0.518 ± 0.042 | 0.432 ± 0.042 | 0.171 ± 0.054 |
|  | 44 | Averbode | 9 | 0 | 2.607 ± 0.177 | 0.579 ± 0.044 | 0.466 ± 0.049 | 0.182 ± 0.065 |
|  | 45 | Arendschot | 9 | 0 | 2.355 ± 0.148 | 0.523 ± 0.042 | 0.435 ± 0.043 | 0.106 ± 0.078 |
|  | 46 | Geel-Bel | 31 | 1 | 2.525 ± 0.143 | 0.547 ± 0.036 | 0.510 ± 0.039 | 0.071 ± 0.040 |
|  | 47 | Kopberg | 21 | 0 | 2.433 ± 0.145 | 0.522 ± 0.039 | 0.431 ± 0.049 | 0.220 ± 0.062 |
|  | 48 | Keiheuvel | 10 | 0 | 2.433 ± 0.148 | 0.528 ± 0.038 | 0.479 ± 0.045 | 0.079 ± 0.064 |
|  | 49 | Hamont-Achel2 | 8 | 0 | 2.379 ± 0.142 | 0.545 ± 0.042 | 0.486 ± 0.053 | 0.114 ± 0.060 |
|  | 50 | Hamont-Achel3 | 9 | 0 | 2.306 ± 0.112 | 0.523 ± 0.033 | 0.521 ± 0.038 | -0.010 ± 0.055 |
|  | 51 | Hamont-Achel1 | 10 | 0 | 2.35 ± 0.123 | 0.532 ± 0.032 | 0.540 ± 0.042 | -0.025 ± 0.055 |
| Wallonia | 52 | Harchies | 19 | 2 | 2.114 ± 0.134 | 0.433 ± 0.049 | 0.336 ± 0.040 | 0.217 ± 0.051 |
|  | 53 | Lagland | 20 | 1 | 2.154 ± 0.124 | 0.450 ± 0.037 | 0.330 ± 0.041 | 0.269 ± 0.057 |

#### Results for subsampled dataset

A summary table for the population-level statistics for the subsample dataset is given in table S7.2. Population level statistics for the subsampled dataset for all sampled population are given in table S7.3 (similar to table S7.1). Figures S7.1-S7.3 give a visualization of these data for all populations.

Table S7.2: as table 1 in main text but for the subsampled populations. Summary table for the population-level statistics (Statistic): rarefied allelic richness (AR), expected (H_e_) and observed (H_o_) heterozygosity, and inbreeding coefficient (F_IS_). Summary calculations are for two regions: Coast (coastal France and coastal Flanders combined) and Inland (inland Flanders and inland Wallonia combined). For each statistic and region, the mean, standard deviation (SD), minimum (min) and maximum (max) of the range are given. To check the difference of the statistics between regions, a two-sided t-test was performed and t-value (t), degrees of freedom (df) and p-value are given.

| **Statistic** | **Region** | **Mean** | **SD** | **Min** | **Max** | **t** | **df** | **p-value** |
| --- | --- | --- | --- | --- | --- | --- | --- | --- |
| AR | Coast | 3.06 | 0.13 | 2.81 | 3.33 | 6.14 | 14.81 | <0.001 |
|  | Inland | 2.65 | 0.23 | 2.29 | 3.05 |  |  |  |
| H_e_ | Coast | 0.59 | 0.03 | 0.53 | 0.64 | 5.64 | 14.00 | <0.001 |
|  | Inland | 0.51 | 0.05 | 0.41 | 0.58 |  |  |  |
| H_o_ | Coast | 0.52 | 0.06 | 0.36 | 0.62 | 3.61 | 20.24 | 0.002 |
|  | Inland | 0.45 | 0.06 | 0.34 | 0.54 |  |  |  |
| F_IS_ | Coast | 0.11 | 0.10 | -0.04 | 0.37 | 0.22 | 26.76 | 0.83 |
|  | Inland | 0.10 | 0.08 | -0.03 | 0.22 |  |  |  |

S7.3: as table S7.1 - population level statistics with the subsampled dataset for all sampled populations (table S1.1). Details are with table S7.1. NA values are for two populations left out of the subsampled dataset (had 5 samples each).

| **Region** | **ID** | **Population**  **name** | **n** | **NP** | **AR ± SE** | **H_e_ ± SE** | **H_o_ ± SE** | **F_IS_ ± SE** |
| --- | --- | --- | --- | --- | --- | --- | --- | --- |
| France-Picardie | 1 | Mt St Frieux | 10 | 1 | 3.040 ± 0.232 | 0.565 ± 0.035 | 0.566 ± 0.045 | 0.003 ± 0.042 |
|  | 2 | Slack | 10 | 0 | 2.874 ± 0.184 | 0.566 ± 0.033 | 0.562 ± 0.049 | -0.002 ± 0.066 |
|  | 3 | Sangatte | 10 | 0 | 3.144 ± 0.278 | 0.603 ± 0.031 | 0.570 ± 0.035 | 0.046 ± 0.040 |
| Flanders-coast | 4 | Westhoek Zuid | 10 | 0 | 2.936 ± 0.243 | 0.557 ± 0.042 | 0.510 ± 0.053 | 0.086 ± 0.069 |
|  | 5 | Cabour | 10 | 0 | 3.250 ± 0.248 | 0.630 ± 0.028 | 0.619 ± 0.033 | 0.004 ± 0.046 |
|  | 6 | Veurne | 10 | 0 | 2.990 ± 0.215 | 0.596 ± 0.036 | 0.467 ± 0.037 | 0.205 ± 0.042 |
|  | 7 | Tropiflora | 10 | 0 | 3.086 ± 0.248 | 0.587 ± 0.039 | 0.510 ± 0.047 | 0.144 ± 0.045 |
|  | 8 | Westhoek vissersdorp | 10 | 0 | 3.001 ± 0.245 | 0.570 ± 0.038 | 0.528 ± 0.043 | 0.061 ± 0.054 |
|  | 9 | Westhoek NO | 10 | 1 | 3.159 ± 0.284 | 0.601 ± 0.034 | 0.565 ± 0.048 | 0.070 ± 0.053 |
|  | 10 | Westhoek Oost | 10 | 0 | 3.120 ± 0.258 | 0.610 ± 0.032 | 0.611 ± 0.058 | -0.013 ± 0.097 |
|  | 11 | Oosthoekduinen | 10 | 0 | 3.172 ± 0.265 | 0.598 ± 0.036 | 0.597 ± 0.051 | 0.008 ± 0.062 |
|  | 12 | Belvedere | 10 | 0 | 3.114 ± 0.239 | 0.597 ± 0.030 | 0.574 ± 0.041 | 0.037 ± 0.050 |
|  | 13 | HB-Noord | 10 | 0 | 3.165 ± 0.234 | 0.583 ± 0.044 | 0.529 ± 0.056 | 0.107 ± 0.062 |
|  | 14 | HB-Doornpanne | 10 | 0 | 3.296 ± 0.247 | 0.617 ± 0.036 | 0.530 ± 0.044 | 0.152 ± 0.050 |
|  | 15 | HB-Oost | 10 | 0 | 3.176 ± 0.249 | 0.611 ± 0.030 | 0.546 ± 0.037 | 0.093 ± 0.056 |
|  | 16 | HB-Pylyserlaan | 10 | 0 | 3.101 ± 0.242 | 0.610 ± 0.032 | 0.552 ± 0.049 | 0.102 ± 0.056 |
|  | 17 | Schipgatduinen | 10 | 0 | 2.935 ± 0.241 | 0.555 ± 0.038 | 0.517 ± 0.032 | 0.033 ± 0.042 |
|  | 18 | Witte Burg | 10 | 0 | 3.070 ± 0.229 | 0.604 ± 0.031 | 0.618 ± 0.038 | -0.040 ± 0.055 |
|  | 19 | Astridlaan | 10 | 0 | 3.107 ± 0.276 | 0.594 ± 0.035 | 0.515 ± 0.038 | 0.124 ± 0.052 |
|  | 20 | Plaatsduinen | 10 | 0 | 2.993 ± 0.227 | 0.570 ± 0.037 | 0.524 ± 0.037 | 0.073 ± 0.037 |
|  | 21 | Oostvoordduinen | 10 | 0 | 2.975 ± 0.223 | 0.593 ± 0.025 | 0.561 ± 0.038 | 0.042 ± 0.064 |
|  | 22 | Ter Yde West | 10 | 0 | 3.030 ± 0.234 | 0.585 ± 0.031 | 0.575 ± 0.036 | -0.004 ± 0.059 |
|  | 23 | Ter Yde IWVA | 10 | 0 | 2.956 ± 0.208 | 0.592 ± 0.029 | 0.539 ± 0.034 | 0.064 ± 0.064 |
|  | 24 | Karthuizerduinen | 10 | 0 | 2.977 ± 0.228 | 0.582 ± 0.035 | 0.539 ± 0.036 | 0.053 ± 0.053 |
|  | 25 | Simliduinen | 10 | 1 | 3.270 ± 0.263 | 0.621 ± 0.033 | 0.562 ± 0.037 | 0.091 ± 0.039 |
|  | 26 | Sint-Laureins1 | 9 | 0 | 3.149 ± 0.252 | 0.602 ± 0.033 | 0.502 ± 0.036 | 0.141 ± 0.065 |
|  | 27 | Sint-Laureins2 | 7 | 0 | 3.328 ± 0.272 | 0.640 ± 0.036 | 0.528 ± 0.049 | 0.173 ± 0.060 |
|  | 28 | Warandeduinen Middelkerke | 7 | 0 | 3.226 ± 0.220 | 0.612 ± 0.034 | 0.517 ± 0.042 | 0.125 ± 0.070 |
|  | 29 | Raversijde1 | 8 | 0 | 3.065 ± 0.222 | 0.597 ± 0.042 | 0.369 ± 0.045 | 0.361 ± 0.069 |
|  | 30 | Raversijde2 | 10 | 0 | 2.950 ± 0.196 | 0.597 ± 0.030 | 0.470 ± 0.038 | 0.211 ± 0.054 |
|  | 31 | Raversijde3 | NA | NA | NA | NA | NA | NA |
|  | 32 | Raversijde4 | NA | NA | NA | NA | NA | NA |
|  | 33 | Fort Napoleon | 9 | 0 | 2.913 ± 0.196 | 0.556 ± 0.036 | 0.517 ± 0.041 | 0.053 ± 0.064 |
|  | 34 | Spanjaardduinen Oostende | 8 | 0 | 3.132 ± 0.229 | 0.617 ± 0.035 | 0.451 ± 0.042 | 0.264 ± 0.055 |
|  | 35 | Bredene | 10 | 1 | 2.938 ± 0.238 | 0.576 ± 0.034 | 0.364 ± 0.044 | 0.359 ± 0.069 |
|  | 36 | Duinbossen DeHaan | 10 | 0 | 2.916 ± 0.196 | 0.606 ± 0.025 | 0.384 ± 0.032 | 0.365 ± 0.051 |
|  | 37 | Zwinbosjes-grazed | 9 | 0 | 2.808 ± 0.205 | 0.544 ± 0.045 | 0.481 ± 0.048 | 0.098 ± 0.054 |
|  | 38 | Zwinbosjes | 10 | 0 | 2.968 ± 0.260 | 0.557 ± 0.040 | 0.515 ± 0.048 | 0.071 ± 0.059 |
|  | 39 | Vloethemveld-Noord | 9 | 0 | 2.811 ± 0.181 | 0.528 ± 0.040 | 0.458 ± 0.037 | 0.102 ± 0.047 |
|  | 40 | Vloethemveld-Zuid | 10 | 0 | 2.994 ± 0.211 | 0.563 ± 0.041 | 0.478 ± 0.040 | 0.140 ± 0.037 |
| Flanders-inland | 41 | Wetteren | 10 | 0 | 2.507 ± 0.147 | 0.478 ± 0.043 | 0.487 ± 0.051 | -0.018 ± 0.047 |
|  | 42 | Kortenhoeff-NL | 10 | 0 | 2.478 ± 0.171 | 0.446 ± 0.046 | 0.392 ± 0.044 | 0.091 ± 0.048 |
|  | 43 | Kalmthout | 10 | 0 | 2.725 ± 0.207 | 0.514 ± 0.043 | 0.413 ± 0.038 | 0.173 ± 0.049 |
|  | 44 | Averbode | 9 | 0 | 3.049 ± 0.258 | 0.579 ± 0.044 | 0.466 ± 0.049 | 0.182 ± 0.065 |
|  | 45 | Arendschot | 9 | 1 | 2.690 ± 0.208 | 0.523 ± 0.042 | 0.435 ± 0.043 | 0.106 ± 0.078 |
|  | 46 | Geel-Bel | 10 | 0 | 2.898 ± 0.194 | 0.547 ± 0.038 | 0.467 ± 0.046 | 0.138 ± 0.068 |
|  | 47 | Kopberg | 10 | 0 | 2.847 ± 0.228 | 0.529 ± 0.047 | 0.472 ± 0.053 | 0.101 ± 0.062 |
|  | 48 | Keiheuvel | 10 | 1 | 2.849 ± 0.214 | 0.528 ± 0.038 | 0.479 ± 0.045 | 0.079 ± 0.064 |
|  | 49 | Hamont-Achel2 | 8 | 0 | 2.629 ± 0.176 | 0.545 ± 0.042 | 0.486 ± 0.053 | 0.114 ± 0.060 |
|  | 50 | Hamont-Achel3 | 9 | 0 | 2.570 ± 0.147 | 0.523 ± 0.033 | 0.521 ± 0.038 | -0.010 ± 0.055 |
|  | 51 | Hamont-Achel1 | 10 | 0 | 2.611 ± 0.164 | 0.532 ± 0.032 | 0.540 ± 0.042 | -0.025 ± 0.055 |
| Wallonia | 52 | Harchies | 10 | 2 | 2.310 ± 0.152 | 0.414 ± 0.047 | 0.339 ± 0.039 | 0.137 ± 0.056 |
|  | 53 | Lagland | 10 | 1 | 2.286 ± 0.184 | 0.418 ± 0.048 | 0.337 ± 0.054 | 0.224 ± 0.083 |

#### Visualizations of population level statistics

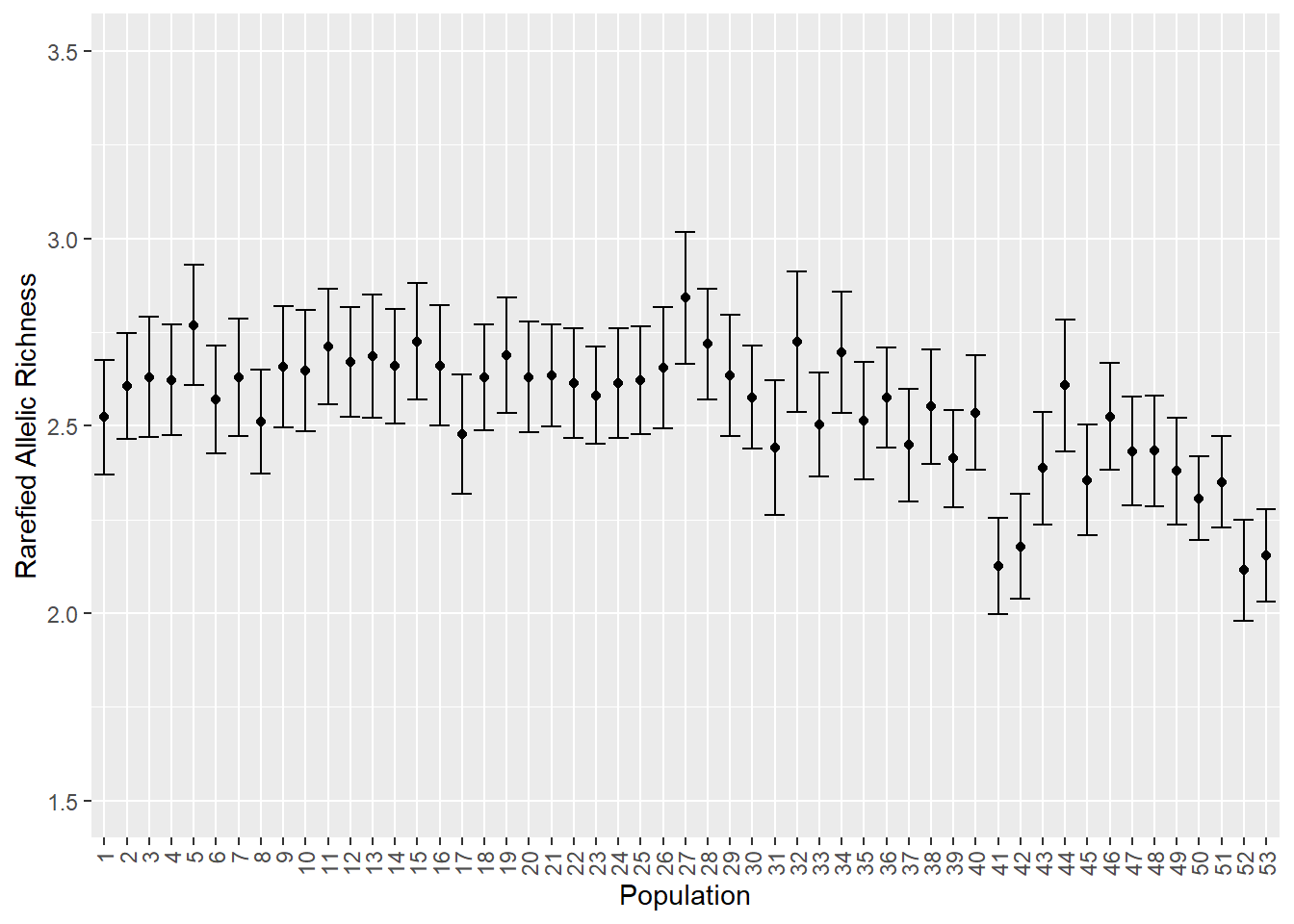

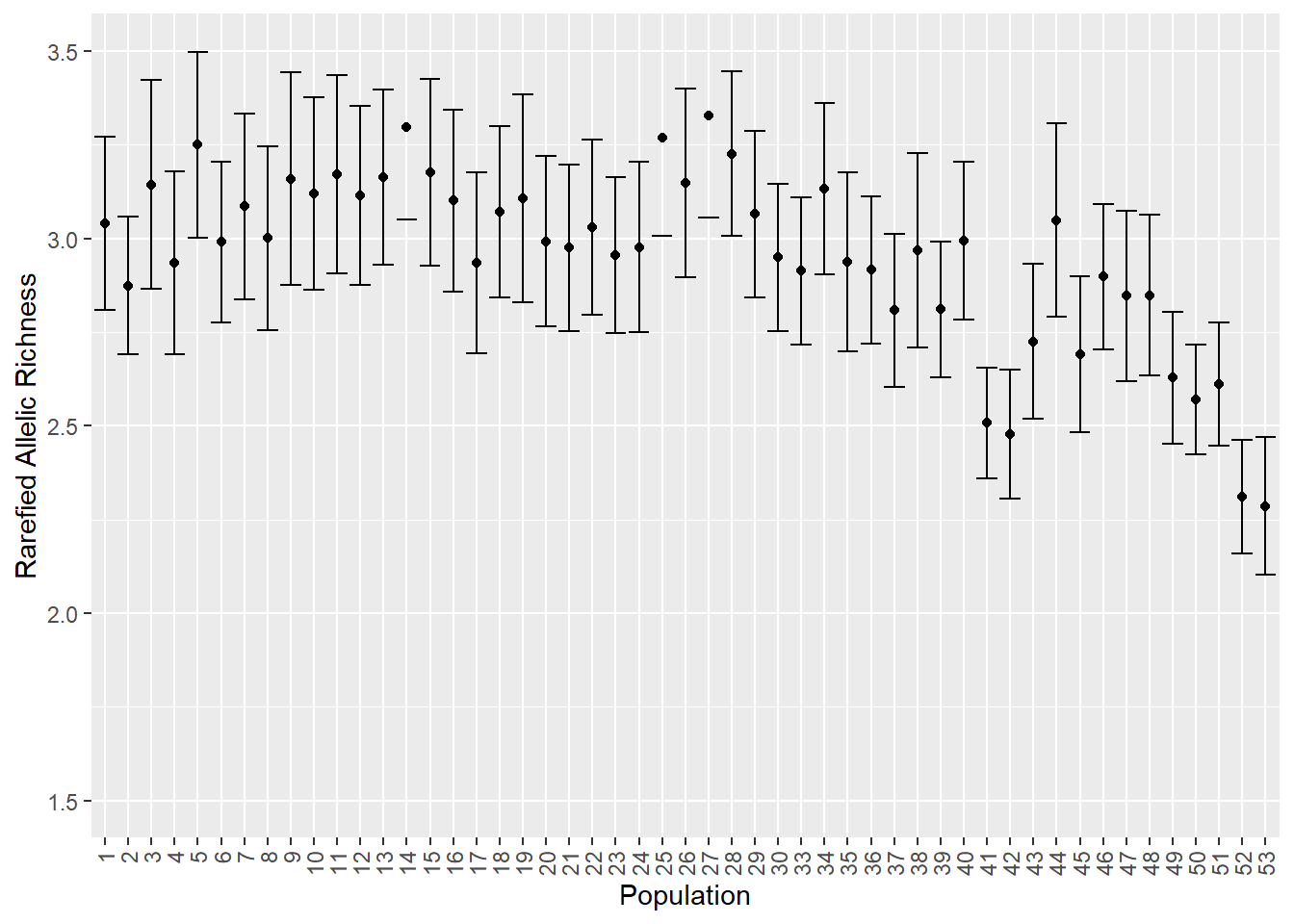

Figure S7.1: visualization of allelic richness for all populations (table S7.1 and table S7.3). Left for complete dataset, right for subsampled dataset. Error bars are standard errors.

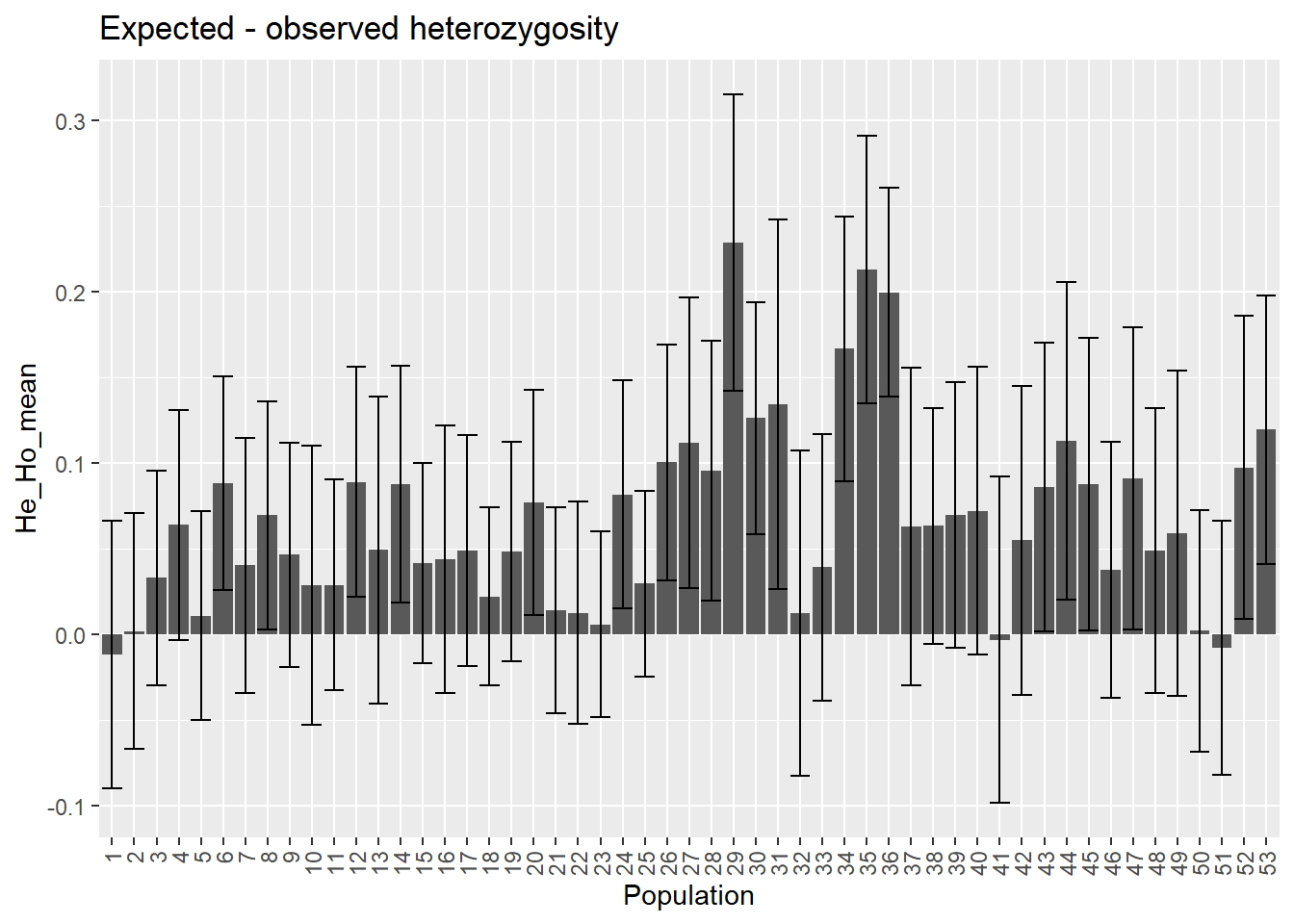

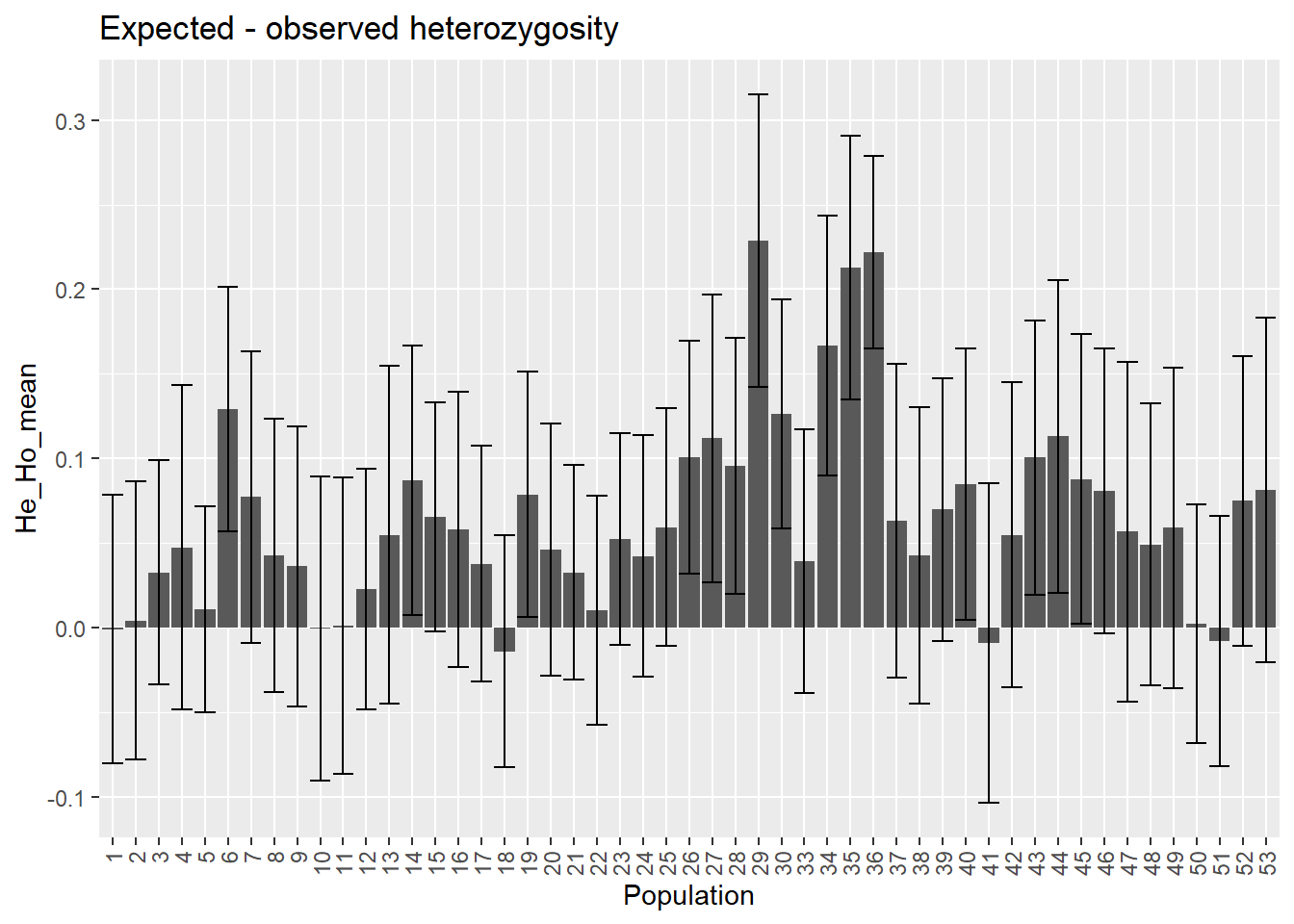

Figure S7.2: visualization of H_e_-H_o_ for all populations (table S7.1 and table S7.3). Left for complete dataset, right for subsampled dataset. Error bars are standard errors.

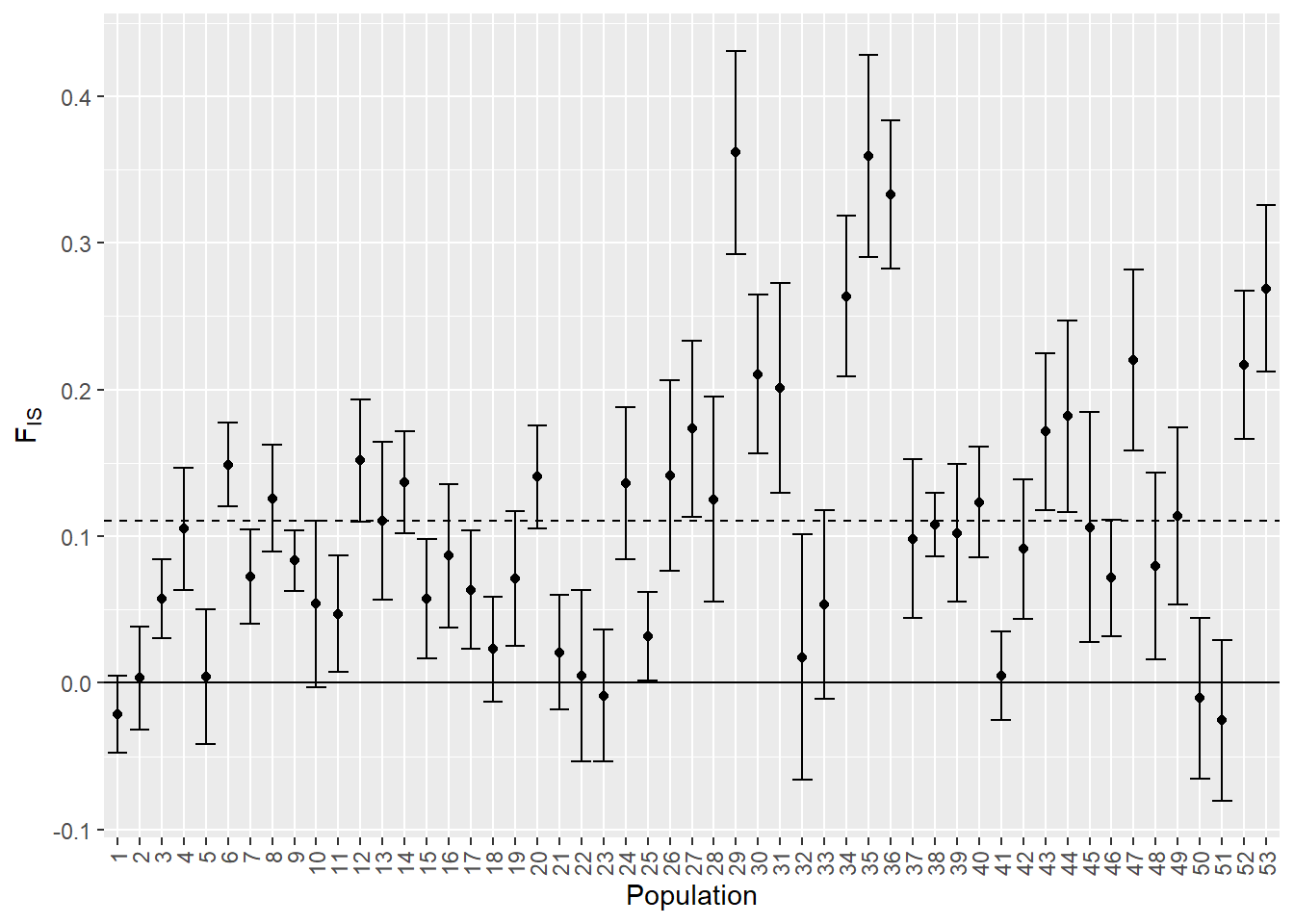

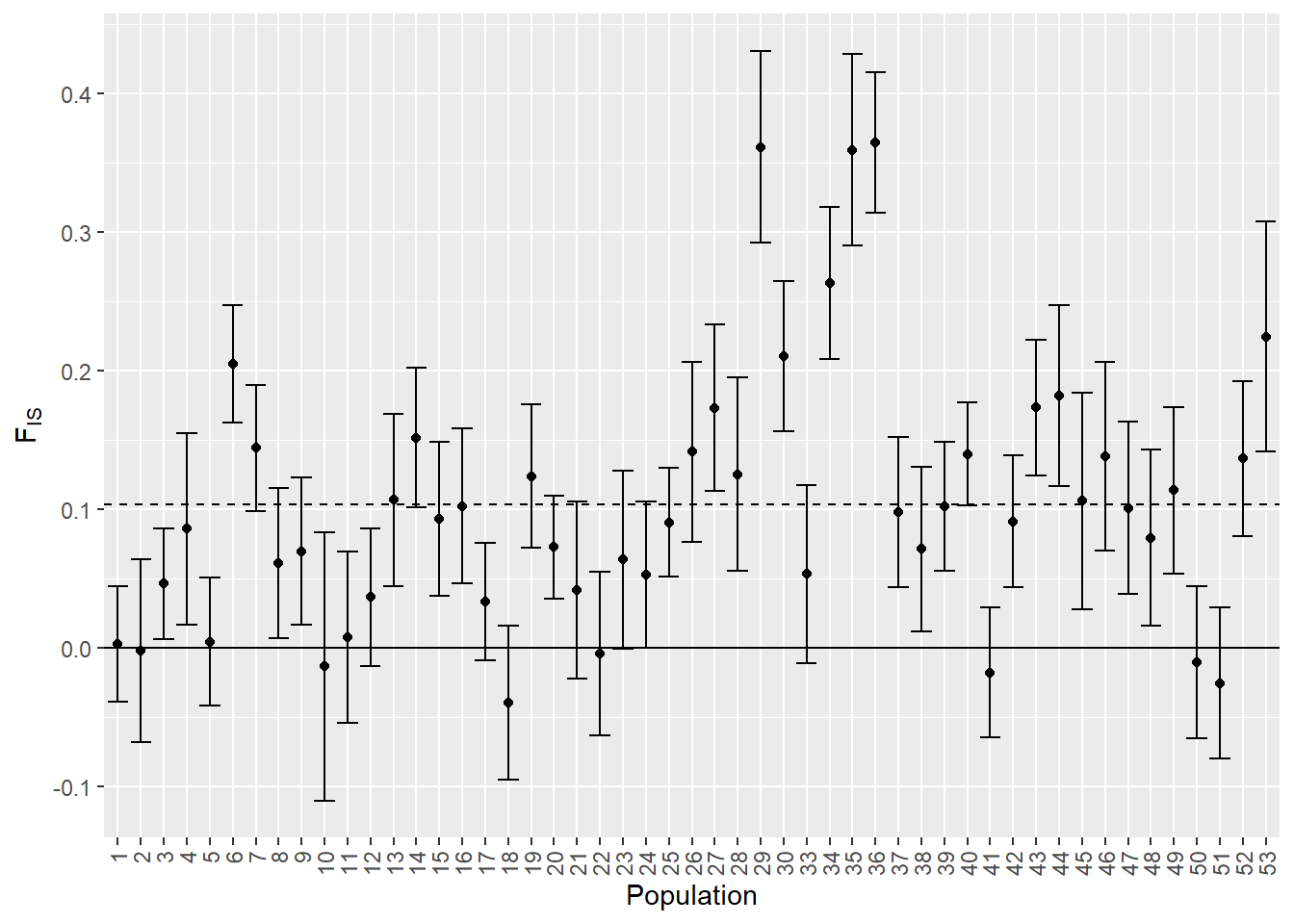

Figure S7.3: visualization of F_IS_ for all populations (table S7.1 and table S7.3). Left for complete dataset, right for subsampled dataset. Error bars are standard errors.

### S8. DAPC scatterplots

#### Coastal and inland regions separately

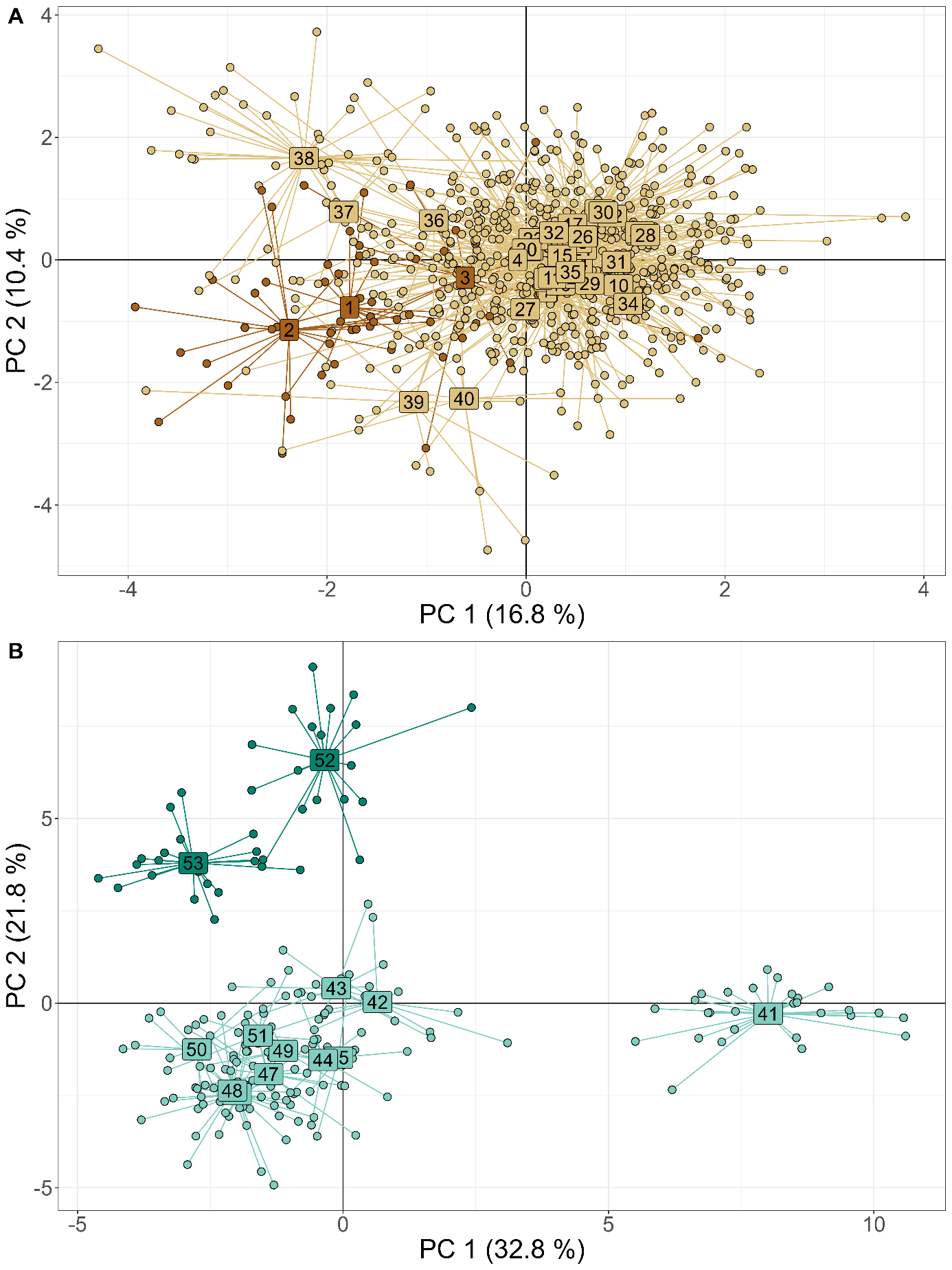

Figure S8.1: DAPC scatterplots for coastal (A) and inland (B) regions separately. Numbers are population IDs and are centered in the point cloud of each population.

### S9. Extra information assignment tests

#### Assignment tests for subsampled dataset

To test the robustness of results of the assignment tests and the derived conclusions, we ran the same assignment tests as described in the main manuscript, but for a subsampled dataset (see XX; XX main manuscript). Table S9 and figure S4 are parallel to figures 5 and 6 from the main manuscript, but now for the subsampled dataset.

Results are similar to the results with the complete dataset in the main manuscript (fig. 5 and 6). Differences are due to either rare links that are not detected anymore (because some random individuals are not present anymore in the dataset) or because the putative source populations have now a more restricted set of alleles and variation present (which makes assigning more difficult). Nevertheless, the general interpretations still holds with the results with this subsampled dataset: more restricted gene flow within inland, high gene flow within coast, asymmetrical gene flow from coast to inland. Examples of some detailed differences are: 1) no genetic links from coastal France to inland. This is probably because the pool of the putative source populations (in France) is now much more constrained and less varied than in the full dataset. As coastal France is well-connected to the coast in Belgium, which holds many more populations, individuals are more easily assigned to these populations in Belgium. 2) There are in general less links, which is as expected from a more restricted dataset. Consequently, some rare links (such as from population 51 to 47; 46 to 47) are not detected anymore. 3) Population 41 has less arrows and a much less thick arrow arriving from coastal populations, which is probably because of a mixture of both main reasons (individuals randomly omitted from the focal population and the variation in alleles in the putative source populations).

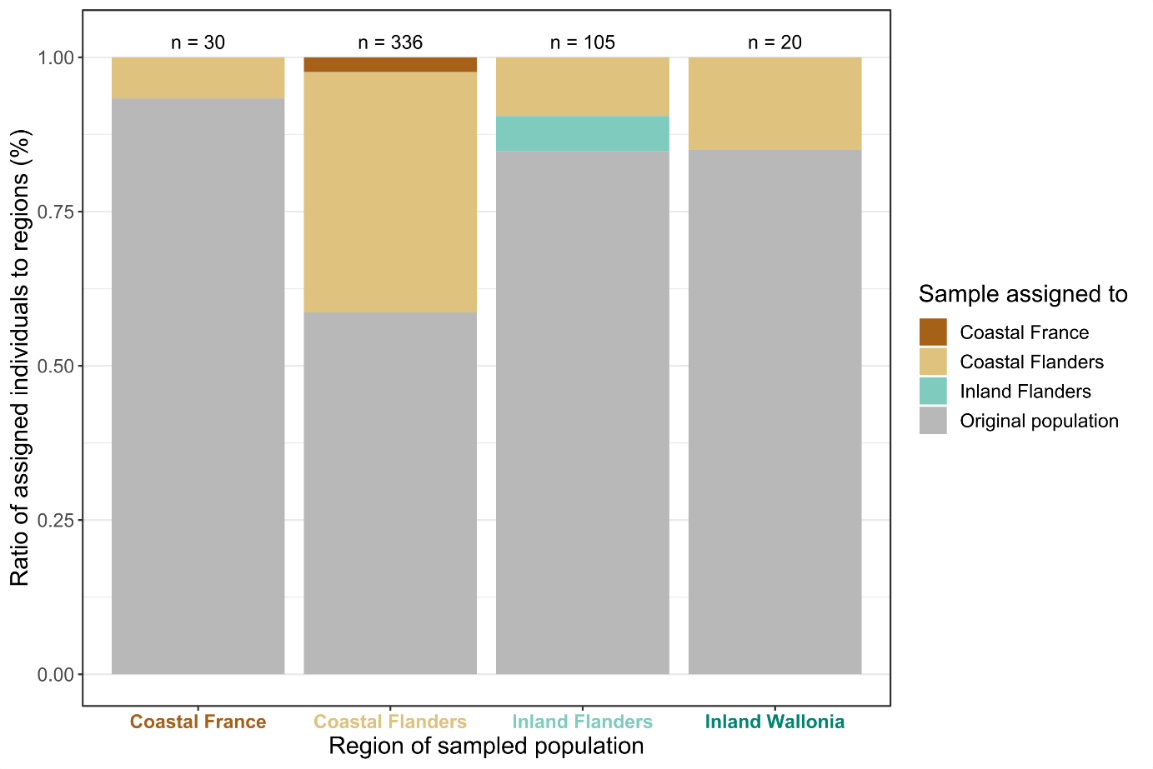

Figure S9.1: barplot of summarized results with the subsampled dataset of the assignment tests for all populations from each sampled region (x-axis), parallel to figure 5 in the main manuscript. If an individual was assigned to its original population where it was sampled, the barplot-area is filled with grey. If an individual was assigned to another population within the same region or to another region, the barplot-area is colored by region (dark brown: assigned to coastal France, light brown: assigned to coastal Flanders, light green: assigned to inland Flanders). Total number samples per region (n) is indicated above each barplot.

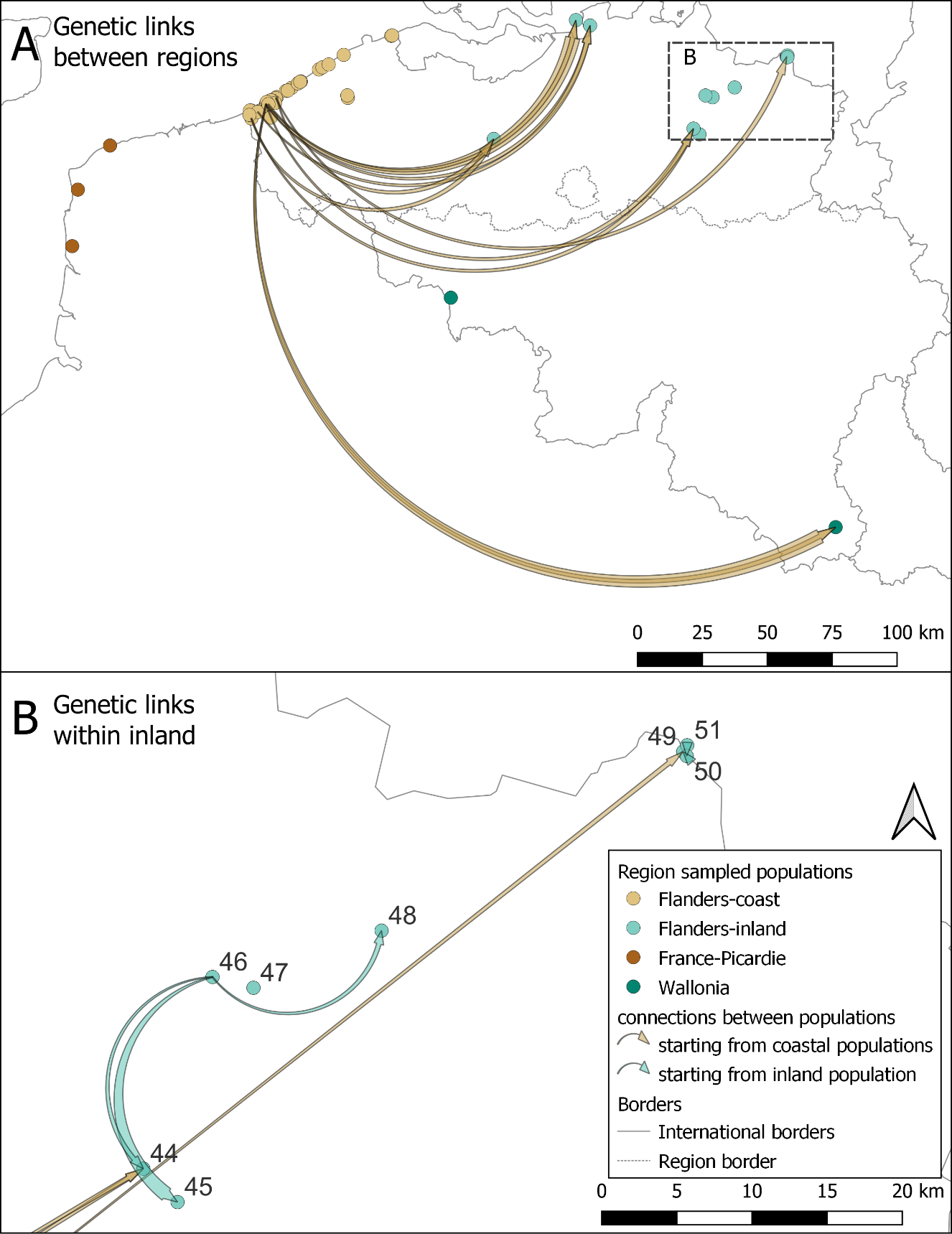

Figure S9.2: flow chart of genetic links (A) between regions (B) within the inland region for *B. rostrata* according to assignment to putative source populations with the subsampled dataset; parallel to figure 5 in the main text. Genetic links within the coastal region are not depicted as these were too numerous (table S9). The links represent the number of individuals assigned to a putative origin population (start of the arrow) that were caught in the sampled population (end of the arrow). Brown arrows are links starting from the coast, green arrows start from inland populations. The thicker the end of the arrow, the higher the number of individuals assigned to the putative source. Genetic links are present from coast to inland, but not from inland to coast (A). Within the inland region, there are only genetic links within a cluster of populations 44 to 50 (B). Individuals from other inland populations (41-43; 52, 53) are either assigned to their sampled population or are assigned to a coastal population (A).

#### Derived dispersal kernel

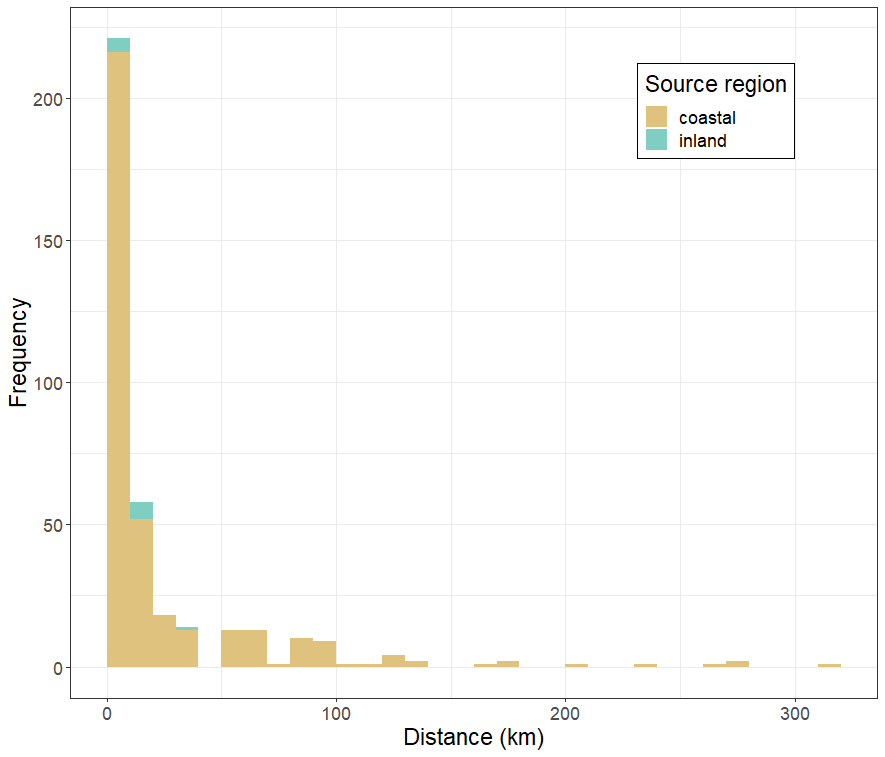

Figure S9.3: histogram of pairwise distances of genetic connections based on assignment tests, depicting an indirectly derived dispersal kernel. Colors depict region of source population: coastal (Flanders and France) and inland (Flanders; population from inland Wallonia were not detected as source populations for any individual sampled in another population).
