## Supplementary materials S1-S4 for "Asymmetrical gene flow between coastal and inland dunes in a threatened digger wasp"

### S1. Details of sampled populations

Table S1.1: details of sampled populations: region, population number (ID; see map main manuscript), given population name (Population name), X- and Y-coordinates (X, Y) in coordinate system Belgian Lambert 72 (epsg:31370), sample year(s) (Year), number of samples (n) and total number of samples used in analyses per population, see AMOVA analysis (n tot).

| **Region** | **ID** | **Population name** | **X** | **Y** | **Year** | **n** | **n tot** |
| --- | --- | --- | --- | --- | --- | --- | --- |
| Coastal France | 1 | Mt St Frieux | -45055.836 | 147290.022 | 2018 | 19 | 19 |
|  | 2 | Slack | -42871.872 | 169089.634 | 2018 | 20 | 20 |
|  | 3 | Sangatte | -30478.698 | 186092.799 | 2018 | 20 | 20 |
| Coastal Flanders | 4 | Westhoek Zuid | 23274.352 | 197883.930 | 2018  2020 | 12  9 | 21 |
|  | 5 | Cabour | 24000.043 | 196525.259 | 2018 | 10 | 10 |
|  | 6 | Veurne | 31418.656 | 196887.771 | 2018 | 20 | 20 |
|  | 7 | Tropiflora | 24109.489 | 198094.010 | 2018  2020 | 20  9 | 29 |
|  | 8 | Westhoek vissersdorp | 23467.294 | 199618.259 | 2018  2020 | 20  3 | 23 |
|  | 9 | Westhoek NO | 24120.869 | 199559.082 | 2018  2020 | 22  8 | 30 |
|  | 10 | Westhoek Oost | 24446.785 | 199084.816 | 2018  2020 | 10  8 | 18 |
|  | 11 | Oosthoekduinen | 26503.156 | 199039.926 | 2018  2020 | 19  10 | 29 |
|  | 12 | Belvedere | 27469.461 | 199611.067 | 2018  2020 | 10  8 | 18 |
|  | 13 | HB-Noord | 29615.521 | 201772.772 | 2018  2020 | 7  9 | 16 |
|  | 14 | HB-Doornpanne | 29652.166 | 202069.447 | 2018  2020 | 10  7 | 17 |
|  | 15 | HB-Oost | 29672.837 | 201505.012 | 2018  2020 | 10  6 | 16 |
|  | 16 | HB-Pylyserlaan | 29915.987 | 201394.673 | 2018  2020 | 10  8 | 18 |
|  | 17 | Schipgatduinen | 29945.549 | 202880.064 | 2018  2020 | 12  4 | 16 |
|  | 18 | Witte Burg | 30561.564 | 202010.765 | 2018  2020 | 12  7 | 19 |
|  | 19 | Astridlaan | 31560.066 | 203551.845 | 2018  2020 | 17  7 | 24 |
|  | 20 | Plaatsduinen | 32343.888 | 203286.675 | 2018  2020 | 18  9 | 27 |
|  | 21 | Oostvoordduinen | 33104.253 | 202483.973 | 2018  2020 | 20  10 | 30 |
|  | 22 | Ter Yde West | 32486.049 | 203868.703 | 2018  2020 | 10  3 | 13 |
|  | 23 | Ter Yde IWVA | 33151.347 | 203994.353 | 2018  2020 | 10  9 | 19 |
|  | 24 | Karthuizerduinen | 33449.436 | 204338.378 | 2018  2020 | 9  9 | 18 |
|  | 25 | Simliduinen | 33814.709 | 204776.506 | 2018  2020 | 20  7 | 27 |
|  | 26 | Sint-Laureins1 | 37336.964 | 207156.451 | 2020 | 9 | 9 |
|  | 27 | Sint-Laureins2 | 38042.734 | 207317.394 | 2020 | 7 | 7 |
|  | 28 | Warandeduinen Middelkerke | 39485.855 | 208330.732 | 2020 | 7 | 7 |
|  | 29 | Raversijde1 | 41944.728 | 210035.728 | 2020 | 8 | 8 |
|  | 30 | Raversijde2 | 42218.098 | 210273.838 | 2020 | 10 | 10 |
|  | 31 | Raversijde3 | 42839.355 | 210691.179 | 2020 | 5 | 5 |
|  | 32 | Raversijde4 | 43074.002 | 210795.354 | 2020 | 5 | 5 |
|  | 33 | Fort Napoleon | 50050.322 | 215428.560 | 2020 | 9 | 9 |
|  | 34 | Spanjaardduinen Oostende | 51403.973 | 216230.489 | 2020 | 8 | 8 |
|  | 35 | Bredene | 53808.105 | 217367.810 | 2020 | 10 | 10 |
|  | 36 | Duinbossen DeHaan | 59600.497 | 221053.034 | 2020 | 12 | 12 |
|  | 37 | Zwinbosjes-grazed | 77921.134 | 228387.482 | 2020 | 9 | 9 |
|  | 38 | Zwinbosjes | 78243.444 | 228544.867 | 2018  2020 | 20  15 | 35 |
|  | 39 | Vloethemveld-Noord | 60996.017 | 205437.804 | 2020 | 9 | 9 |
|  | 40 | Vloethemveld-Zuid | 61176.550 | 204480.049 | 2020 | 11 | 11 |
| Inland Flanders | 41 | Wetteren | 117432.840 | 188634.139 | 2018  2020 | 8  20 | 28 |
|  | 42 | Kortenhoeff-NL | 149108.508 | 234460.777 | 2020 | 10 | 10 |
|  | 43 | Kalmthout | 154547.132 | 232375.786 | 2020 | 12 | 12 |
|  | 44 | Averbode | 194424.342 | 192633.330 | 2020 | 9 | 9 |
|  | 45 | Arendschot | 196699.590 | 190406.561 | 2020 | 9 | 9 |
|  | 46 | Geel-Bel | 199027.845 | 205395.375 | 2018  2020 | 20  11 | 31 |
|  | 47 | Kopberg | 201751.857 | 204674.642 | 2020 | 21 | 21 |
|  | 48 | Keiheuvel | 210274.429 | 208479.656 | 2020 | 10 | 10 |
|  | 49 | Hamont-Achel2 | 230347.333 | 220397.920 | 2020 | 8 | 8 |
|  | 50 | Hamont-Achel3 | 230584.834 | 220105.267 | 2020 | 9 | 9 |
|  | 51 | Hamont-Achel1 | 230612.992 | 220822.730 | 2020 | 10 | 10 |
| Inland Wallonia | 52 | Harchies | 100876.462 | 127411.662 | 2021 | 19 | 19 |
|  | 53 | Lagland | 249207.562 | 38937.742 | 2021 | 20 | 20 |

### S2. DNA-extraction and PCR protocol

#### DNA-extraction protocol for wing tips *Bembix rostrata*

Eppendorfs with samples dried to the air were put in liquid nitrogen for a few minutes and homogenized with a crusher. 50 μL Chelex and 10 μL proteinase K were added and the samples were incubated (56°C) overnight. The boiling and centrifuge steps were performed the next morning.

Day 1

- Put a wing tip in a 1.5mL ep, let it dry for several minutes in the open air (to let the ethanol evaporate)
- Put the ep for 5’ in liquid nitrogen
- Crush the wing tip firmly with a crusher
- Pipet 50 µL 6% Chelex InstaGene Matrix solution (Biorad) (while it is on the magnetiser) into the ep, while rinsing off the crusher
- Add 10 µL proteinase K (N600mAU/ml, Qiagen)
- Incubate overnight at 56°C (+ soft shaking)

Day 2

- Vortex (+ minicentrifuge) to get the drops of the lid
- 15’ 99°C
- Vortex (+ minicentrifuge) to get the drops of the lid
- Centrifuge 3’ at 14 000 rpm and store in the fridge awaiting PCR-amplification

#### PCR protocol

For the primer mix, a ratio of ‘3 : 1 : 3’ ‘Forward primer : Reverse primer (tailed) : oligonucleotides dye (FAM, VIC, NED, PET)’ is used (1μM concentration). Table S2 gives the partition of primers into 3 pairs of multiplexes: for each series/plate of samples, 6 PCRs were run (6 multiplexes: 1.1, 1.2, 2.1, 2.2, 3.1, 3.2). These PCR products were diluted and pairwise combined (1.1+1.2, 2.1+2.2, 3.1+3.2) before sending the PCR products to the ABI analyzer.

We worked with a total volume in each well of a PCR-plate of 5 μl:

- 2 μl Qiagen MultiPlex (Qiagen® Multiplex PCR kit cat. No. 206143)
- 2 μl microsatellite primer mix
- 1 μl of DNA

If the tissue material for the DNA extraction was very small, the 2 μl of DNA was air dried by putting it on a block heater (37°C) for at least 5 hours. This way, the concentration of DNA was increased in the PCR-volume.

PCR conditions:

- 95°C for 15 min
- 35 cycles of

94°C for 30 sec

57°C for 90 sec

72°C for 60 sec

- 60°C for 30 min
- 4°C for ∞

Table S2.1: Partitioning of primers into 3 pairs multiplexes: 6 PCRs are run for each sample, but 3 PCRs products are used for the ABI analyzer.

| 1.1 | | | | | | | 2.1 | | | | | | 3.1 | | | |  |  |
| --- | --- | --- | --- | --- | --- | --- | --- | --- | --- | --- | --- | --- | --- | --- | --- | --- | --- | --- |
| 3 µL | F110 | F16 | F486 | F9 | F227 | F57 | 3 µL | F76 | F144 | F234 | F32 | F35 | 6 µL | F12 | F138 | F161 |  |  |
| 1 µL | R110 | R16 | R486 | R9 | R227 | R57 | 1 µL | R76 | R144 | R234 | R32 | R35 | 2 µL | R12 | R138 | R161 |  |  |
| 9 µL | 6-FAM M13-21 | | | VIC M13-21 | | | 9 µL | 6-FAM M13-21 | | |  |  | 3 µL | F329 | F111 |  |  |  |
| Dilution post PCR: 2**x** | | | | | | | 6 µL | VIC M13-21 | |  |  |  | 1 µL | R329 | R111 |  |  |  |
|  |  |  |  |  |  |  | Dilution post PCR: **4x** | | | | | | 12 µL | 6-FAM | VIC M13-21 |  |  |  |
|  |  |  |  |  |  |  |  |  |  |  |  |  | Dilution post PCR: 4**x** | | | |  |  |
| 1.2 | | | |  |  |  | 2.2 | | | | | | 3.2 | | | | | |
| 6 µL | F403 | F299 |  |  |  |  | 6 µL | F20 |  |  |  |  | 6 µL | F337 |  |  |  |  |
| 2 µL | R403 | R299 |  |  |  |  | 2 µL | R20 |  |  |  |  | 2 µL | R337 |  |  |  |  |
| 3 µL | F116 | F375 | F467 |  |  |  | 3 µL | F437 | F487 | F298 | F266 | F153 | 3 µL | F196 | F419 | F480 | F218 | F307 |
| 1 µL | R116 | R375 | R467 |  |  |  | 1 µL | R437 | R487 | R298 | R266 | R153 | 1 µL | R196 | R419 | R480 | R218 | R307 |
| 12 µL | NED M13 Moda | | |  |  |  | 9 µL | NED M13 Moda | | |  |  | 6 µL | NED M13 Moda | |  |  |  |
| 9 µL | PET T7 |  |  |  |  |  | 12 µL | PET T7 | | |  |  | 15 µL | PET T7 | | | |  |
| Dilution post PCR: **4x** | | | |  |  |  | Dilution post PCR: 2**x** | | | | | | Dilution post PCR: **3x** | | | | | |

### S3. List of primers

Table S3.1: Characteristics of the 33 microsatellites used in our study for *Bembix rostrata*. A full list of the newly developed (by AllGenetics®, A Coruña, Spain) remaining non-tested (and tested but discarded) microsatellite loci can be obtained from the contacting author. Annealing temperature used for all primers is 57°C. Five loci had a lot of stutter in the amplification profiles and were discarded from the analysis (AGBro486, -329, -196, -437, -298). Following abbreviations are used: the multiplex mix in which the marker was included (Primer Mix) and fluorescent label used (Label), the observed fragment length range (Size range), the number of alleles observed (No. alleles), observed and expected heterozygosity (Ho and He), if they are used in the genetic analyses after assumption testing (Used) and both primer sequences (F = forward, R = reverse).

| **Locus** | **Repeat motif** | **No repeats** | **Primer Mix** | **Label** | **Size range (bp)** | **No. of alleles** | **Ho** | **He** | **Used** | **Primer sequences (5’-3’)** |
| --- | --- | --- | --- | --- | --- | --- | --- | --- | --- | --- |
| AGBro110 | AAG | 7 | 1.1 | FAM | 95-124 | 7 | 0.58 | 0.6189 | y | F: GCCATCACGTTTACAGCCAC  R: AGAGGTGGTAGTGCTGGAGA |
| AGBro16 | AGGC | 7 | 1.1 | FAM | 188-208 | 5 | 0.1437 | 0.1573 | n | F: CTCCGCGTACTATTTCCGCT  R: TAACAGCGTGGTTTCCGGAA |
| AGBro486 | AG | 11 | 1.1 | FAM | 366-414 | / | / | / | n | F: ACAAACGTTTCACGGTACTTGT  R: GGCAGGAGGACATCGTTGAT |
| AGBro9 | AG | 14 | 1.1 | VIC | 90-130 | 14 | 0.7089 | 0.8144 | y | F: GAGTGAGAGGGAGGCAGAGA  R: TACGTGCCTGAGGAAACGAC |
| AGBro227 | AACG | 6 | 1.1 | VIC | 144-180 | 8 | 0.3978 | 0.4198 | y | F: CAAGCATGGCCAGTCTGTTT  R: TGGCTACTGTGGGCTCACTA |
| AGBro57 | AG | 7 | 1.1 | VIC | 225-235 | 2 | 0.114 | 0.2806 | n | F: GCACCGGGACACTGTCTTAT  R: GGGTCACTCGATCGACGTTT |
| AGBro403 | AACG | 10 | 1.2 | NED | 122-180 | 12 | 0.6876 | 0.7435 | y | F: TTATCCCGATCGCTTGGCAT  R: GCTCGACGCTTCATCGATTAAC |
| AGBro116 | AG | 8 | 1.2 | NED | 213-235 | 5 | 0.4787 | 0.5597 | y | F: CTCCTCTCCATACGACGCAC  R: TTGGCAGTAGAACGAGGACC |
| AGBro375 | ACAG | 8 | 1.2 | NED | 267-279 | 3 | 0.3719 | 0.4279 | y | F: CGAAGTTCCGCATTACCTTGC  R: CCTGACAGGTGCTCACGTATT |
| AGBro299 | AC | 8 | 1.2 | PET | 100-123 | 5 | 0.4092 | 0.4418 | y | F: GACGTAAGGGCGAAGAACGT  R: GCATTCCGTGCGAGTGAATA |
| AGBro467 | AAG | 7 | 1.2 | PET | 158-202 | 11 | 0.6525 | 0.7317 | y | F: GTCCAGAGAAGGTATGAGAGGG  R: CGTGACGTAATATCCGGGCA |
| AGBro76 | AAG | 7 | 2.1 | FAM | 100-122 | 7 | 0.5356 | 0.5813 | y | F: AATTGTGCCGAAACTTGGCC  R: TCGTTGCAAGTGTCGTGACA |
| AGBro144 | ACGG | 7 | 2.1 | FAM | 152-192 | 8 | 0.6386 | 0.745 | y | F: GCCGTTTATCCGTCCATCCA  R: CCTCTCATATCGGTGCCTCC |
| AGBro234 | AG | 8 | 2.1 | FAM | 270-282 | 4 | 0.5425 | 0.605 | y | F: CGTGCTCCACCCGAAATTCT  R: CAGCTGCAGTTCGATGATCG |
| AGBro32 | AAGC | 5 | 2.1 | VIC | 133-149 | 4 | 0.4853 | 0.4955 | y | F: GCGGCTGGTATCTGATCCAA  R: CTAGCTGTCTGCCTACCTGC |
| AGBro35 | AAG | 5 | 2.1 | VIC | 229-240 | 2 | 0.1405 | 0.3423 | n | F: AAGTTCTCACGAAACCGCCT  R: GGGCCACCAGATTCTTACCC |
| AGBro437 | AG | 7 | 2.2 | NED | 104-110 | / | / | / | n | F: CGATAGCAAGCACGCGAGG  R: GAGGGTAAACCACGAGGAGC |
| AGBro487 | ACG | 12 | 2.2 | NED | 146-176 | 6 | 0.5191 | 0.6037 | y | F: GGGAGAGTTCGCGAAGGTAC  R: CCTTCAGAAATGCTTGTCGTTGT |
| AGBro298 | AG | 10 | 2.2 | NED | 230-306 | / | / | / | n | F: AGCTTGTTGGACGCGTAAGA  R: GGCGATCGACATTTCAGTCA |
| AGBro20 | AAT | 6 | 2.2 | PET | 309-320 | 3 | 0.0759 | 0.1001 | n | F: TCTGATTGGACCGTTCGTCG  R: ACGTGTCTGATCGTGCTTGT |
| AGBro266 | AT | 6 | 2.2 | PET | 112-122 | 4 | 0.3306 | 0.4099 | y | F: CGCGAACATTAAGCACCGAA  R: ATACCGTCACGACAGAGCCA |
| AGBro153 | AG | 7 | 2.2 | PET | 160-170 | 4 | 0.4447 | 0.5075 | y | F: TAGCTCAGCCTCTACCGACC  R: TGAACGAGAACGGCGTACAG |
| AGBro161 | AAAG | 6 | 3.1 | FAM | 248-263 | 4 | 0.4679 | 0.556 | y | F: TCGCCGTAAGACCTTCGTAC  R: GGTATGCGGTCTTCCTGGTG |
| AGBro329 | ACC | 6 | 3.1 | FAM | 85-133 | / | / | / | n | F: TCCCTCTTCGTTCCTCTCCT  R: CTCGGCGAAAGATAGCACGG |
| AGBro111 | AG | 8 | 3.1 | FAM | 162-184 | 9 | 0.5723 | 0.6913 | n | F: TGTACCAATCCGGCCTTTGT  R: AACGTACGGTGGATTAGCCG |
| AGBro12 | AG | 8 | 3.1 | VIC | 198-220 | 10 | 0.5872 | 0.6902 | y | F: GTGCCGTAATTCGACGAACG  R: ATCTCGTAACGTTCCTCGCC |
| AGBro138 | AAG | 6 | 3.1 | VIC | 95-112 | 3 | 0.1869 | 0.3075 | n | F: ACTGCCGTACCTGTAGCTTC  R: CTTTCACGCTTCGCACATGT |
| AGBro196 | AG | 14 | 3.2 | NED | 126-158 | / | / | / | n | F: ATGGCGAAGGAAACGGTCTT  R: CTCCCTCGCGTATTTCTCCT |
| AGBro419 | AG | 10 | 3.2 | NED | 248-264 | 6 | 0.398 | 0.5849 | n | F: TGTGACCAGTGGTAACCCAT  R: TGCACCCACTGTCCATATAGC |
| AGBro337 | AGC | 6 | 3.2 | PET | 113-130 | 5 | 0.3928 | 0.4102 | y | F: CGACGGGACCCAATTCATCG  R: ACCATCCTCTTTCTACCGCC |
| AGBro480 | AAAGC | 7 | 3.2 | PET | 163-183 | 4 | 0.5074 | 0.5932 | y | F: TCGTTTACTGTCGCAAATGACC  R: TTGCTTCTCTTCGCTCCACT |
| AGBro218 | AG | 6 | 3.2 | PET | 299-315 | 4 | 0.4393 | 0.4919 | y | F: CAATACCGTCAACTCACCCGA  R: CTGACACCTGACGGATAGCC |
| AGBro307 | AAG | 6 | 3.2 | PET | 220-236 | 4 | 0.3323 | 0.451 | y | F: GTCGCAGCTGATAGCCAAGT  R: ACGACTTATGTCCACGTGGA |

### S4. Genetic differentiation measures

Figures S4.1-4.2 are similar to figure 2 in the main manuscript with Nei’s standardized genetic distance (D_S_), but for the genetic differentiation measures F_ST_ (Weir and Cockerham 1984) and D (Jost 2008). Genetic differentiation measures were calculated with the R package diveRsity (Keenan et al. 2013). Unbiased confidence intervals of 95% were calculated from 1,000 bootstraps.

Tables S4.1-4.3 list the 10 populations with the highest genetic distances and differentiation values, which have the highest average differentiation from all other populations.

These two differentiation measures give similar results as the pairwise Nei’s standardized genetic distance used in the main manuscript: differentiation values are overall high among inland sampling sites, and low among coastal sites. Values are medium to high between coastal and inland regions.

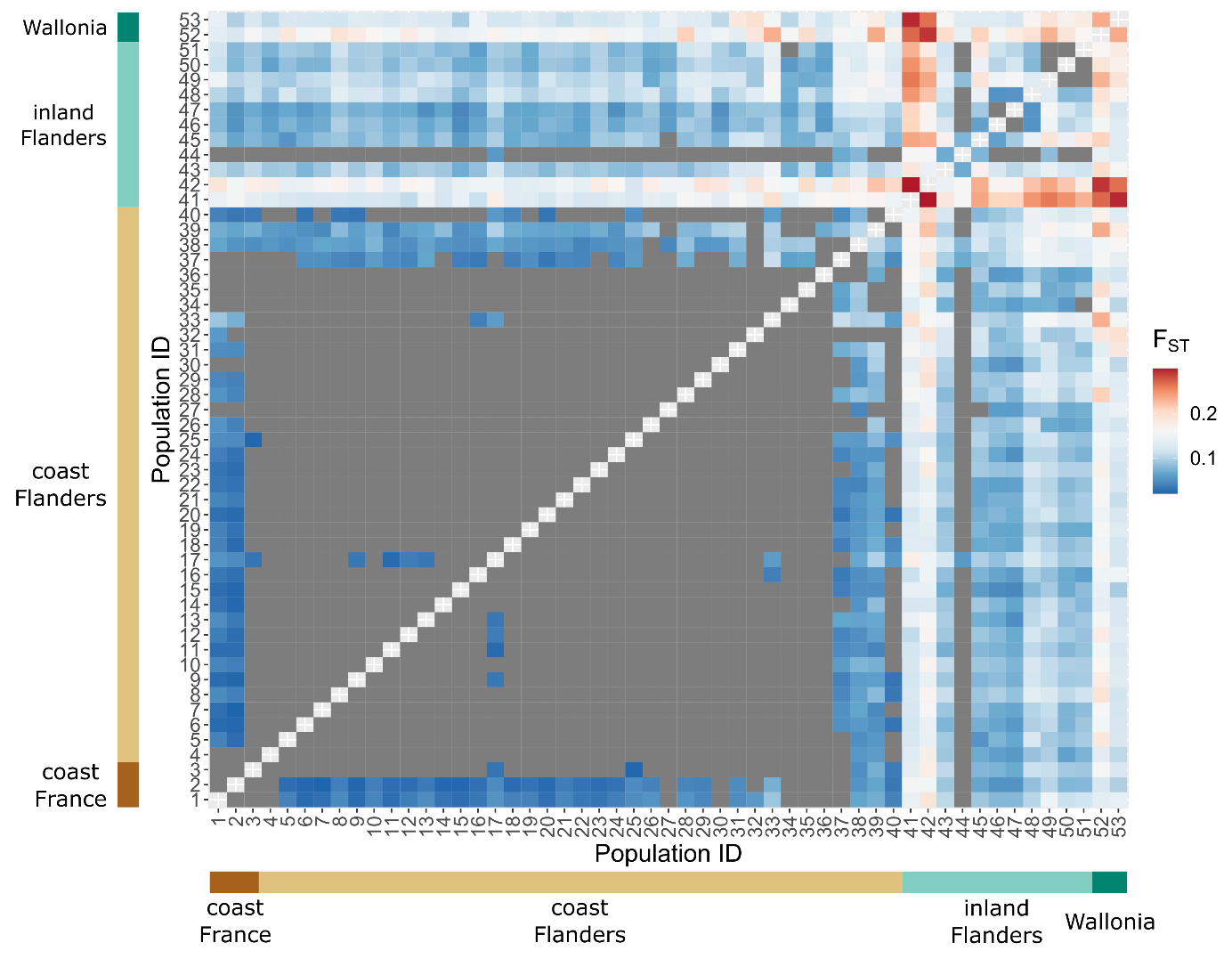
Figure S4.1: graphical matrix representation of pairwise genetic differentiation measure F_ST_: blue are low, white are mid, and red are high pairwise genetic differentiation values between populations. Grey areas are non-significant pairwise genetic differentiation values (0 was included in the 95% confidence intervals of 95% were calculated from 1,000 bootstraps). The x- and y-axes represent the population ID, subdivided in the four different regions. Genetic distances are symmetrical and consequently the matrix is mirrored along the diagonal. There is overall high genetic differentiation within the inland regions (right upper corner) and low genetic differentiation within the coastal regions (left lower corner). The genetic distances between coastal and inland regions are medium to high.

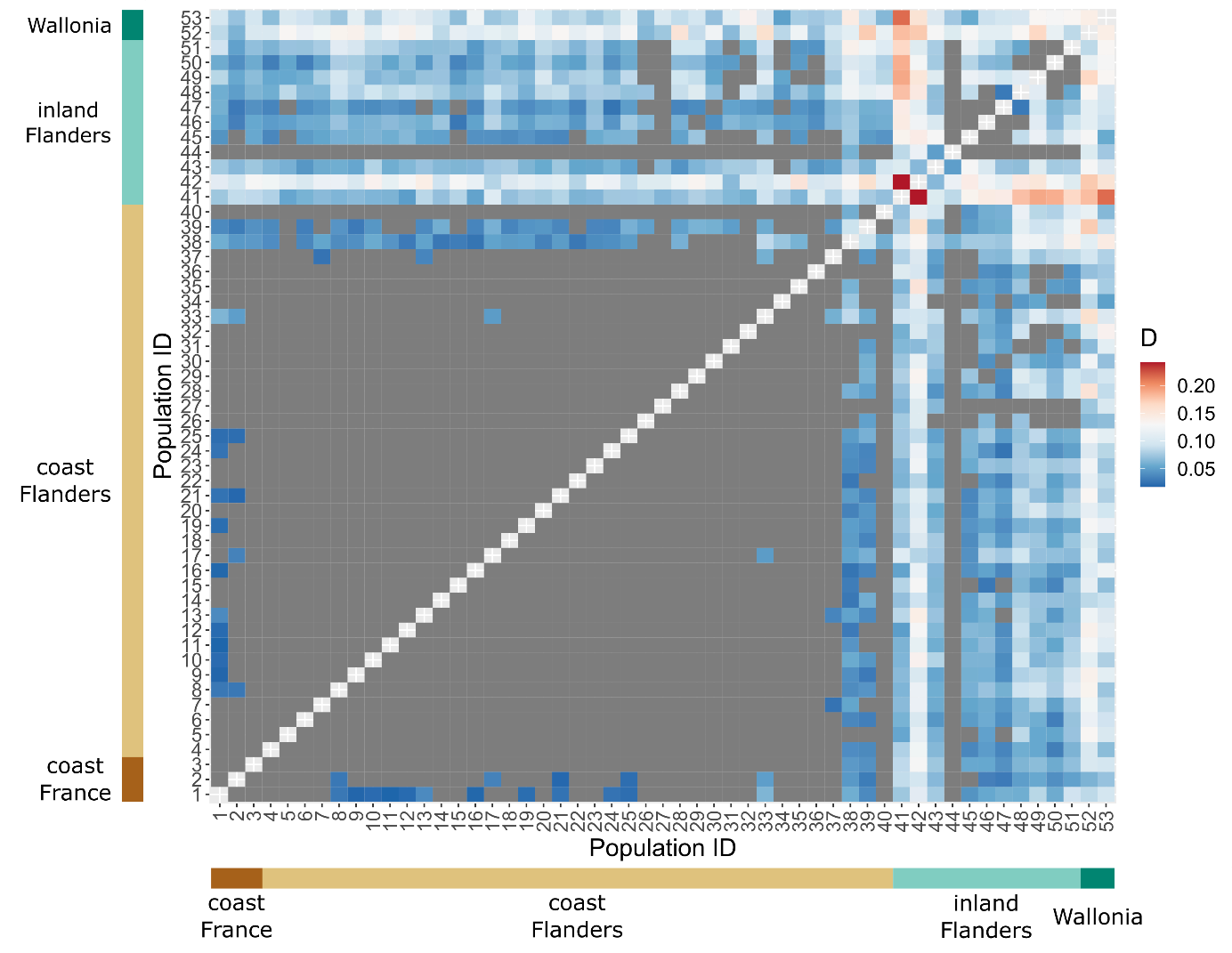
Figure S4.2: graphical matrix representation of pairwise genetic differentiation measure Jost’s D: blue are low, white are mid, and red are high pairwise genetic differentiation values between populations. Grey areas are non-significant pairwise genetic differentiation values (0 was included in the 95% confidence intervals of 95% were calculated from 1,000 bootstraps). The x- and y-axes represent the population ID, subdivided in the four different regions. Genetic distances are symmetrical and consequently the matrix is mirrored along the diagonal. There is overall high genetic differentiation within the inland regions (right upper corner) and low genetic differentiation within the coastal regions (left lower corner). The genetic distances between coastal and inland regions are medium to high.

Table S4.1: the 10 populations with highest averaged Nei’s distance values (Mean(D_S_)). ID is population ID from figure 1 and table S1.1, SD(D_S_) is the standard deviation of mean(D_S_).

| **Region** | **Population name** | **ID** | **Mean(D_S_)** | **SD(D_S_)** |
| --- | --- | --- | --- | --- |
| Flanders-inland | Kortenhoeff-NL | 42 | 0.291 | 0.069 |
| Flanders-inland | Hamont-Achel2 | 49 | 0.257 | 0.075 |
| Wallonia | Harchies | 52 | 0.256 | 0.063 |
| Flanders-inland | Keiheuvel | 48 | 0.230 | 0.065 |
| Flanders-inland | Wetteren | 41 | 0.229 | 0.082 |
| Wallonia | Lagland | 53 | 0.215 | 0.065 |
| Flanders-inland | Arendschot | 45 | 0.207 | 0.069 |
| Flanders-inland | Kalmthout | 43 | 0.201 | 0.048 |
| Flanders-inland | Hamont-Achel1 | 51 | 0.200 | 0.061 |
| Flanders-inland | Halomt-Achel3 | 52 | 0.197 | 0.061 |

Table S4.2: the 10 populations with highest averaged F_ST_ values (Mean(F_ST_)). ID is population ID from table S1.1, SD(F_ST_) is the standard deviation of mean(F_ST_).

| **Region** | **Population** | **ID** | **Mean(F_ST_)** | **SD(F_ST_)** |
| --- | --- | --- | --- | --- |
| Flanders-inland | Kortenhoeff-NL | 42 | 0.173 | 0.039 |
| Wallonia | Harchies | 52 | 0.171 | 0.035 |
| Flanders-inland | Wetteren | 41 | 0.161 | 0.049 |
| Wallonia | Lagland | 53 | 0.142 | 0.039 |
| Flanders-inland | Hamont-Achel2 | 49 | 0.119 | 0.047 |
| Flanders-inland | Keiheuvel | 48 | 0.116 | 0.039 |
| Flanders-inland | Kalmthout | 43 | 0.106 | 0.029 |
| Flanders-inland | Arendschot | 45 | 0.101 | 0.043 |
| Flanders-inland | Hamont-Achel1 | 51 | 0.1 | 0.04 |
| Flanders-inland | Hamont-Achel3 | 50 | 0.098 | 0.043 |

Table S4.3: the 10 populations with highest averaged Jost’s D values (Mean(D)). ID is population ID from table S1.1, SD(D) is the standard deviation of mean(D).

| **Region** | **Population** | **ID** | **Mean(D)** | **SD(D)** |
| --- | --- | --- | --- | --- |
| Flanders-inland | Kortenhoeff-NL | 42 | 0.119 | 0.028 |
| Wallonia | Harchies | 52 | 0.116 | 0.028 |
| Wallonia | Lagland | 53 | 0.097 | 0.029 |
| Flanders-inland | Wetteren | 41 | 0.094 | 0.045 |
| Flanders-inland | Keiheuvel | 48 | 0.079 | 0.030 |
| Flanders-inland | Hamont-Achel2 | 49 | 0.077 | 0.036 |
| Flanders-inland | Hamont-Achel1 | 51 | 0.073 | 0.032 |
| Flanders-inland | Kalmthout | 43 | 0.070 | 0.019 |
| Flanders-inland | Hamont-Achel3 | 50 | 0.061 | 0.032 |
| Flanders-inland | Geel-Bel | 46 | 0.061 | 0.026 |

#### Results for genetic distance and differentiation measures for subsampled dataset

Figures S4.3-4.5 are also similar to figure 2 and the two previous figures, but for subsampled populations (dataset with per population 10 randomly drawn samples when the population had more than 10 samples, see main manuscript). When compared, these figures show qualitatively the same results as for the full dataset. More detailed results (e.g. as tables S4.1-S4.3) can be found in the online code.

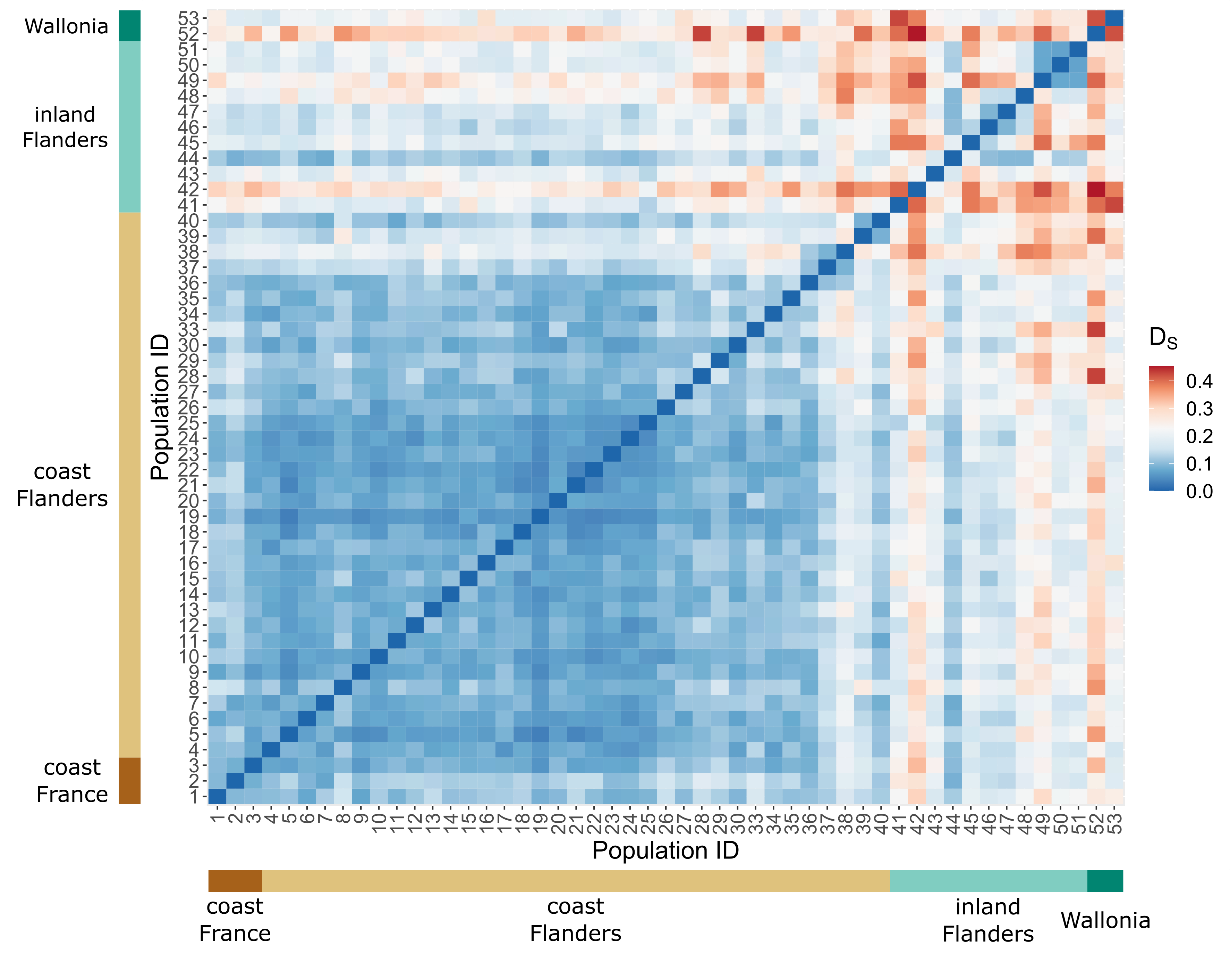

Figure S4.3: Nei’s standardized genetic distance (D_S_) as in fig. 2 in the main manuscript, but for the subsampled dataset (max 10 samples randomly drawn per population from full dataset).

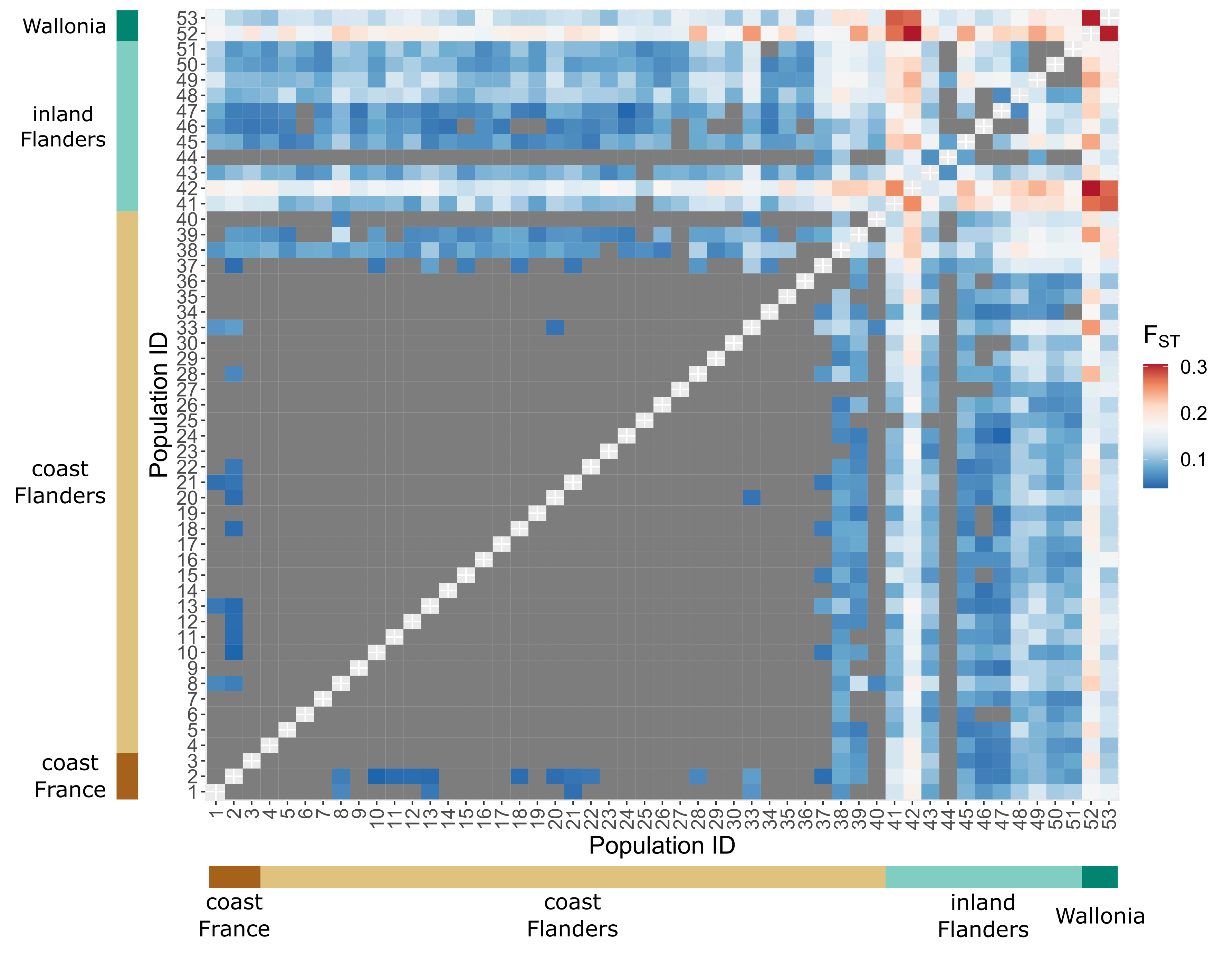

Figure S4.4: pairwise genetic differentiation measure F_ST_ as in fig. S1, but for the subsampled dataset (max 10 samples randomly drawn per population from full dataset).

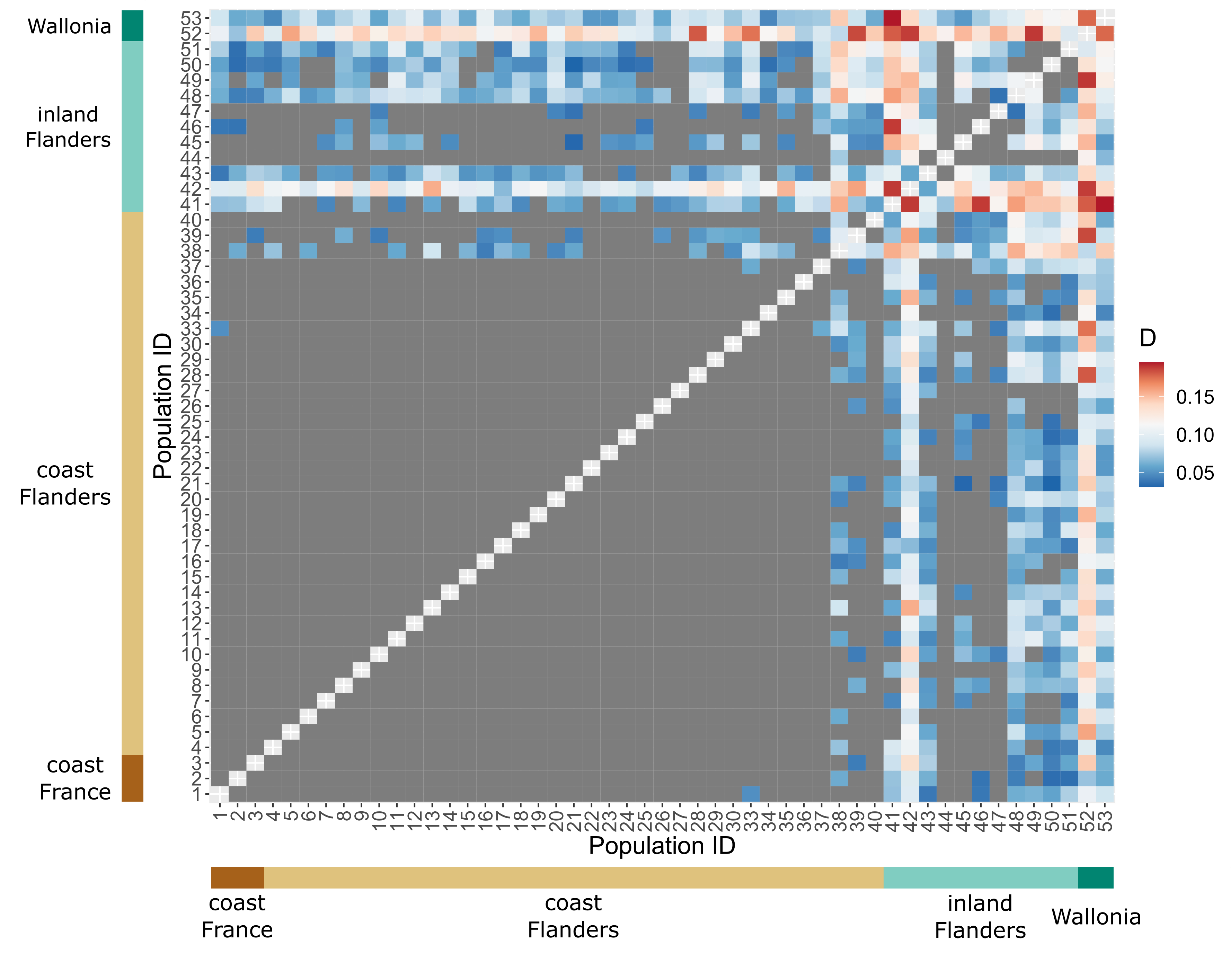

Figure S4.5: pairwise genetic differentiation measure Jost’s D as in fig. S2, but for the subsampled dataset (max 10 samples randomly drawn per population from full dataset).
